# Environmentally-mediated selection parallels population divergence across a chimpanzee subspecies contact zone

**DOI:** 10.1101/2024.07.26.605171

**Authors:** Matthew W. Mitchell, Walker Alexander, Dana V. Mitchell, Adam H. Freedman, Janina Dordel, Ryan J. Harrigan, Ahmet Sacan, Fabrice Kentatchime, Bryan S. Featherstone, Ekwoge E. Abwe, Paul R. Sesink Clee, Abwe E. Abwe, Sabrina Locatelli, Bethan J. Morgan, Bernard Fosso, Roger Fotso, Sarah A. Tishkoff, Evan E. Eichler, Nicola M. Anthony, Thomas B. Smith, Mary Katherine Gonder

**Affiliations:** Department of Biology, Drexel University, Philadelphia, PA, USA; Coriell Institute for Medical Research, Camden, NJ, USA; School of Biomedical Engineering, Science and Health Systems, Drexel University, Philadelphia, PA, USA; Faculty of Arts and Sciences Informatics Group, Harvard University, Cambridge, MA, USA; Center for Tropical Research, Institute of Environment and Sustainability, University of California, Los Angeles, CA, USA; Department of Ecology and Conservation Biology, Texas A&M University, College Station, TX, USA; San Diego Zoo Wildlife Alliance, Escondido, CA, USA; Cameroon Biodiversity Association, Douala, Cameroon; Institut de Recherche pour le Développement (IRD), Maladies Infectieuses et Vecteurs: Ecologie, Génétique, Evolution et Contrôle (MIVEGEC) (IRD 224-CNRS 5290-Université de Montpellier), Montpellier, France; School of Natural Sciences, University of Stirling, Stirling, UK; Wildlife Conservation Society Cameroon, Yaoundé, Cameroon; Departments of Genetics and Biology, University of Pennsylvania, Philadelphia, PA, USA; Department of Genome Sciences, University of Washington School of Medicine, Seattle, WA, USA; Howard Hughes Medical Institute, University of Washington, Seattle, WA, USA; Department of Biological Sciences, University of New Orleans, New Orleans, LA, USA; Department of Ecology and Evolutionary Biology, University of California, Los Angeles, CA, USA

## Abstract

Species evolve from populations with ancestor-descendant relationships in a bifurcating process shaped by geography, gene flow, genetic drift, and natural selection leading to local adaptation to prevailing environmental and ecological conditions. Building on this foundational understanding, we explored local adaptation in chimpanzees (*Pan troglodytes*) at a key geographical intersection in Cameroon where the two main chimpanzee phylogenetic lineages converge. The Nigeria-Cameroon chimpanzee (*P. t. ellioti*) and central chimpanzee (*P. t. troglodytes*) last shared a common ancestor about 500 thousand years ago, with occasional gene flow between them. The evolutionary processes driving their prolonged separation are not fully understood, but neutral evolutionary mechanisms alone cannot account for the observed divergence pattern. Cameroon is often referred to as ‘Africa in miniature’ because the Gulf of Guinea Forest, Congo Basin Forest, and savanna converge there, forming an ecotone. Thus, this contact zone between subspecies in Cameroon provides a unique natural laboratory that enabled us to investigate how environmental variation and natural selection shape divergence in chimpanzees. We developed a genome-wide panel of single-nucleotide polymorphisms (SNPs) in 112 wild chimpanzees sampled in multiple habitats across this contact zone. We augmented SNP discovery by sequencing eight new chimpanzee genomes from Cameroon and analyzing them with previously published chimpanzee genomes. We found that *P. t. ellioti* and *P. t. troglodytes* diverged from one another around 478,000 years ago and occasionally exchange migrants. We identified 1,690 unique SNPs across 905 genes associated with 31 environmental variables that describe the habitat. These genes are involved in essential biological processes, including immune response, neurological development, behavior, and dietary adaptations. This study highlights the importance of understanding the geographical context of natural selection, paving the way for future studies to interpret evidence for genetic variation with phenotypic traits and deepening our understanding of how populations diverge in response to environmental pressures.

**Author Summary:** We investigated how local adaptation contributes to shaping the diversification of chimpanzee subspecies at the geographical convergence point for the two major branches of the chimpanzee phylogenetic tree. We analyzed genome-wide SNP genotypes of 112 chimpanzees sampled from natural communities located in this understudied area. We used tiered methods that identified 905 genes subject to selection, each associated with one or more of 31 environmental predictors describing the habitat. We found strong signals of selection in immune response genes that separate *P. t. troglodytes* from *P. t. ellioti*, highlighting the important role of different pathogen histories in their evolution. We also found evidence of selection in genes associated with neurological development, behavior, and diet, that separate both the subspecies and populations of *P. t. ellioti* that occupy different niches. These findings suggest that ecological and cultural factors may also contribute to shaping the diversification of chimpanzees across the contact zone.

## Introduction

Species consist of populations of reproductively compatible individuals with ancestor–descendant relationships that evolve through time [1]. Speciation may result from various factors. It may have a geographical dimension ranging from allopatry to sympatry with varying degrees of gene flow among populations, genetic drift, and natural selection [2]. Ecological factors often play a decisive role in this process through the local adaptation of populations to prevailing environmental conditions [3]. The fusion of genomics with ecological modeling has advanced the ability to identify loci under environmental selection. It contributes to understanding how species adapt to specific habitats and its impact on speciation [4, 5]. While this link has been studied in many taxa [6], it has been an especially strong focus in studies of human evolution. Human populations have adapted to a multitude of environments [5], disease landscapes [4, 7, 8], and diets including the ability to digest milk into adulthood [9], fatty acid digestion [10], foraging practices in tropical African rainforests [11], cereal-rich diets [12], and persistence in high-altitude environments [13–15].

By comparison, the factors that contribute to shaping the evolution of non-human great ape species are poorly understood. Genomic tools have contributed substantially to resolving the evolutionary relationships and histories of great ape species, subspecies, and some populations [16–20]. However, these studies generally assumed that neutral evolutionary processes (i.e., genetic drift) largely explain the partitioning of genetic variation in great apes. In particular, population genetic structure has been presented as evidence for allopatric speciation in ‘Pleistocene Refugia,’ among gorillas [21], isolation across conspicuous geographic boundaries like rivers [21–23], and separation on different islands [20]. However, a growing body of evidence supports the hypothesis that local adaptation due to natural selection occupies an essential role in shaping the patterning of genetic variation and speciation in great apes [19, 24–26].

Among the great apes, chimpanzees (*Pan troglodytes*) have been particularly well-studied, including analysis of genomes from a representative sample of captive individuals [16, 18, 19] and population genetic studies of natural populations [22–24, 27]. The overall picture from these studies is that the species originated in western equatorial Africa about 1mya. By 500kya in the Middle Pleistocene, two lineages began to diverge from the ancestral *Pan* population: a western lineage composed of the subspecies *P. t. verus* and *P. t. ellioti*, and a central/eastern lineage comprising *P. t. troglodytes* and *P. t. schweinfurthii* (**Fig. 1a**). Major rivers, lakes, and the Dahomey Gap are thought to have acted as dispersal barriers that separate the subspecies to different degrees and timescales, potentially leading to allopatric speciation among chimpanzee subspecies. Among these dispersal barriers, the Sanaga River in Cameroon stands out (**Fig. 1b**). It separates the chimpanzee phylogenetic tree into its two main branches yet remains permeable to occasional gene flow between *P. t. ellioti* and *P. t. troglodytes* [18, 24, 27]. The Sanaga River has likely enabled some degree of allopatric divergence due to genetic drift but the role that natural selection may have played in separating these chimpanzee subspecies remains unknown.

**Fig 1.**
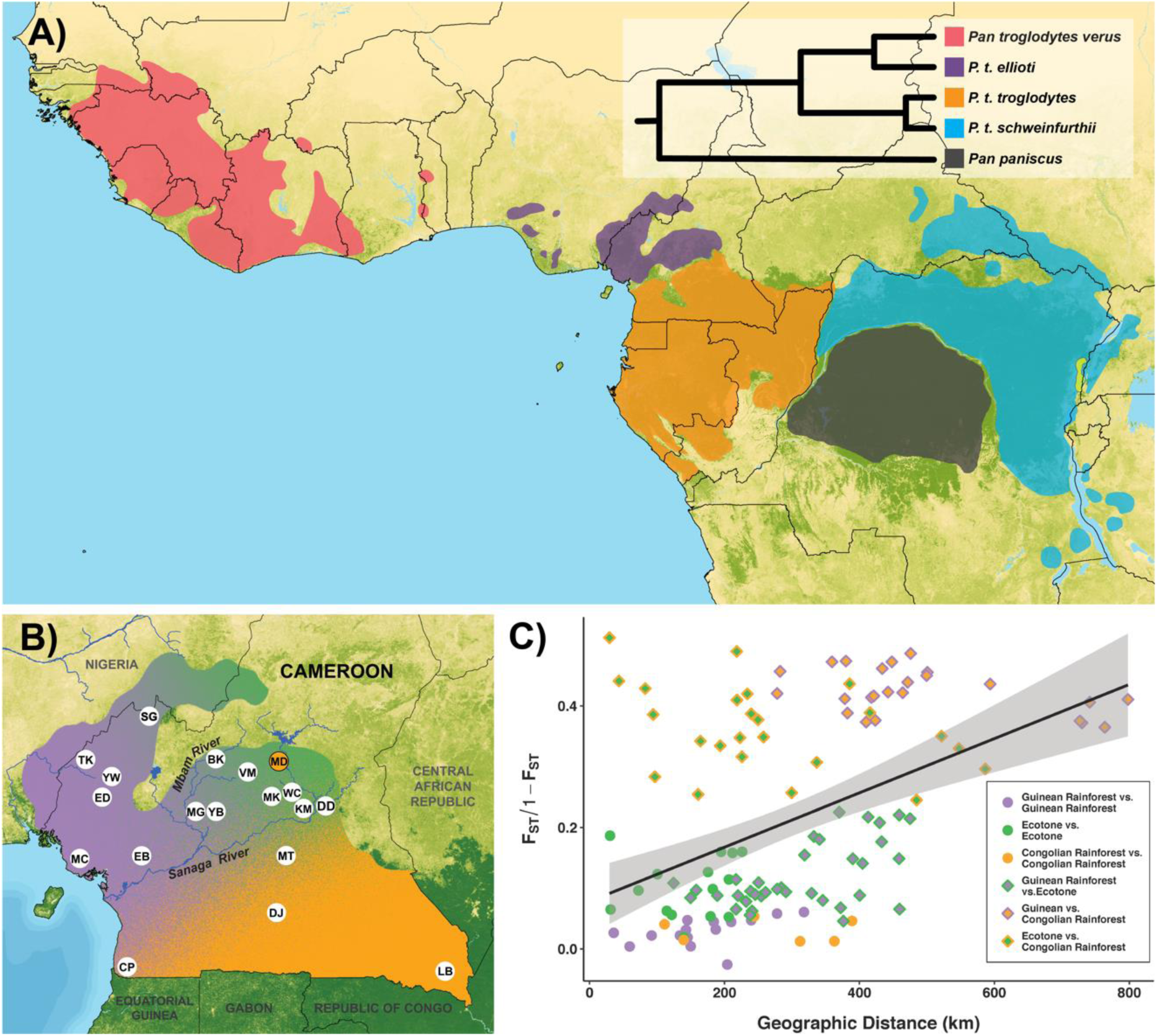
Chimpanzee evolutionary history across Africa and population structure in Cameroon. (A) Distribution and phylogeny of the genus *Pan*. (B) Sampling locations of wild chimpanzee populations in Cameroon overlaid on spatial interpolation of population structure using SNPs from wild chimpanzees. The ‘MD’ sampling location is shaded orange to signify the presence of a *P. t. ellioti*/*P. t. troglodytes* F1 hybrid (CMMD06). (C) Isolation-by-environment in wild chimpanzees in Cameroon. Correlation between ‘linearized *F*_ST_’ and geographic distance (km) generated using SNPs from wild chimpanzees. Solid circles represent pairs of sampling locations from the same habitat. Dual-colored diamonds represent pairs of sampling locations from different habitats. Colors correspond to chimpanzee population origin: purple – *P. t. ellioti* (Rainforest), green – *P. t. ellioti* (Ecotone), and orange – *P. t. troglodytes*.

Natural selection has numerous opportunities to contribute to genetic divergence that may vary between subspecies or populations in different habitats. Life history traits and pathogen defense stand out as likely candidates for establishing among-population divergence due to local adaptation Among these, the role of pathogens is best understood. Differences in pathogen presence and prevalence have long been associated with genotypic differences among great apes, especially chimpanzees. For instance, wild chimpanzee populations are infected to different degrees with several disease-causing pathogens, including malaria [28], Ebola [29], and viruses like simian immunodeficiency virus (SIV) [30]. In the case of SIV and similar viruses, it is relatively well established that these pathogens have exerted selective pressure on chimpanzees, particularly the central and eastern subspecies [31–33]. Interestingly, Cameroon is a unique disease landscape for chimpanzees, especially concerning the puzzling distribution of SIVcpz. Unlike *P. t. troglodytes* and *P. t. schweinfurthii*, SIVcpz has not been found in *P. t. ellioti* or *P. t. verus*, despite extensive sampling [34–36] (**Fig. 1a**).

Secondly, each chimpanzee subspecies occupies a distinct set of environmental niches [37], creating opportunities for adaptation to local environmental conditions. Although little is known about the links among genotypes, phenotypes, and environmental conditions, chimpanzees in arid environments are more efficient in salt removal than their counterparts in more humid forested environments [38]. However, the role of local adaptation to specific environments remains largely unexplored yet is perhaps the most intriguing avenue of investigation in their evolution. Chimpanzees, like humans, have complex social systems and behaviors and maintain diverse cultural traditions [39]. Similarly, cultural variation among chimpanzee communities may lead to localized gene-culture co-evolution, potentially facilitating adaptation to diverse habitats [37] that are vulnerable to human encroachment [40]. Habitat variation and resource availability, specifically food types, are also known to affect chimpanzee socioecological patterns directly [41], yet whether this variation translates into heritable genetic differences remains speculative.

We investigated how local adaptation has influenced the evolution of chimpanzees in Cameroon, a key region where the western and central/eastern lineages of chimpanzees converge. Despite the wealth of research on the contributions of neutral evolutionary processes to the genetic variation found in wild chimpanzees, the contribution of natural selection remains a significant knowledge gap that our study aimed to fill. We employed a two-tier approach to identify genic regions under selection from a comprehensive analysis of natural chimpanzee communities sampled intensively across Cameroon. First, we used whole-genome sequencing (WGS) data from 24 previously published chimpanzee genomes [16], along with eight newly sequenced genomes of individuals from Cameroon to create and annotate a map of genomic regions under natural selection from this expanded sample of complete genomes of chimpanzees originating from Cameroon. Second, we used the analysis of this expanded sample of genomes to create a genome-wide panel of ancestry-informative putatively neutral SNPs, as well as SNPs that fell within signals of positive selection (inferred with the WGS data) and, thus, were good candidates for performing tests to assess local adaptation. We genotyped these SNPs in 112 wild chimpanzees sampled across multiple habitats in Cameroon, encircling the contact zone between *P. t. ellioti* and *P. t. troglodytes*, and that represent the diversity of habitats occupied by chimpanzees across the contact zone [42], including the northern extent of the Congo Basin Forest, the lowland and montane Gulf of Guinea Forest, and the forest/savanna ecotone that bridges these two forest ecosystems. Finally, we used these SNP panels to investigate the relationship between individual SNPs and a suite of SNPs representing the genome to understand the relationship between allele frequencies and environmental variability. Our objective was to assess whether environmental pressures from differing ecologies have influenced allele frequency variation across these wild populations.

## Results

We used WGS data from 24 previously published chimpanzee genomes [16], along with eight newly sequenced genomes of individuals from Cameroon from the Limbe Wildlife Center referred to hereafter as ‘captive chimpanzees’ (**Fig. S1** and **Table S1**). We used the captive chimpanzee dataset and previously published data [16] to create an annotated map of genomic regions under natural selection. Second, we used a genome-wide panel of SNPs in 112 wild chimpanzees sampled across multiple habitats across Cameroon (**Fig. 1b**) to develop a high-resolution, spatially explicit map of allele frequencies to understand the link between habitat variation and loci under selection.

### Captive chimpanzee genome analysis and SNP discovery

#### Developing SNP datasets

We identified SNPs from 32 chimpanzee genomes across all four subspecies, which included eight newly sequenced genomes from the contact zone between the western and central/eastern chimpanzee lineages. After quality filtering, we retained 12,754,225 high-quality SNPs. Based on this initial whole-genome SNP set, two datasets were created. The first dataset was thinned for linkage disequilibrium (LD), retaining only SNPs with r^2^ ≤ 0.1, which resulted in 1,113,142 SNPs retained. The second dataset was thinned to include only SNPs that followed our neutrality criteria (**Methods**), resulting in 147,700 SNPs. **S1 Text** provides additional details on heterozygosity (**Fig. S2**) and population cluster analyses (**Figs. S3 and S4**).

#### Genome scans for signals of selection and defining genomic ‘outlier’ regions

To calculate a test statistic for cross-population extended haplotype homozygosity (XP-EHH) and integrated haplotype score (iHS), SNP-based results were summarized into windows following Pickrell et al. [43], but chromosomes were split into 100kb windows and SNPs were binned in 100 SNP increments. We merged windows indicating positive selection for each method and population. The analysis identified regions specific to the two lineages, and those shared among the Western and Central/Eastern lineages were analyzed separately. **Table 1** summarizes outlier regions for the XP-EHH and iHS and combined outlier tests. In the Western lineage, we found 335 outlier windows stretching 83.5 Mb with 695 candidate genes. The Central/Eastern lineage had 318 windows stretching 81 Mb with 682 genes. We found 25 windows over 13.6 Mb with 80 candidate genes shared between lineages. We plotted the distribution of the outlier regions on individual chromosomes (**Fig. 2a**).

**Fig 2.**
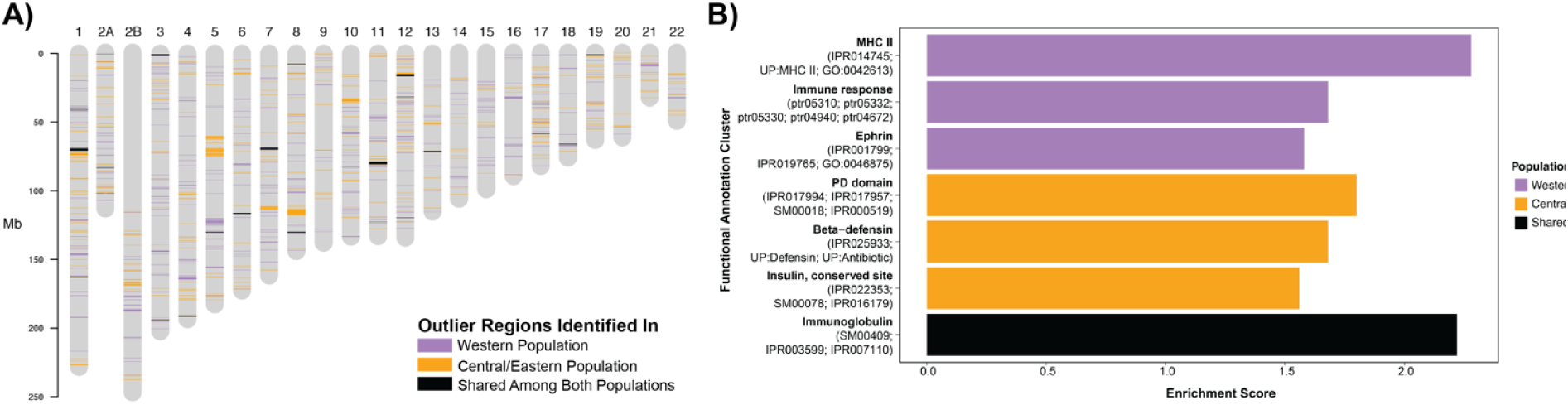
Natural selection in chimpanzees. (A) Regions under selection found using captive chimpanzee genomes plotted on individual chromosomes. (B) Functional enrichment clustering of genes and pathways under selection in chimpanzees. Enriched functional annotation clusters (based on genes in outlier regions) including their respective enrichment score. The name of one functional annotation of each cluster was taken to represent the complete cluster.

**Table 1.** Summary of captive chimpanzee whole genome “outlier” regions.

|  | XP-EHH |  | iHS |  | Combined |  |  |
| --- | --- | --- | --- | --- | --- | --- | --- |
|  | Western lineage | Central/Eastern lineage | Western lineage | Central/Eastern lineage | Western lineage | Central/Eastern lineage | Shared |
| Windows found | 257 | 270 | 118 | 95 | 335 | 318 | 25 |
| Base pairs in windows | 53,753,675 | 55,149,110 | 34,000,000 | 32,200,000 | 83,453,675 | 81,049,110 | 13,600,085 |
| Protein coding genes used in enrichment analysis | 563 | 563 | 152 | 144 | 695/610 | 682/593 | 80/70 |

While all chromosomes are affected by selective sweeps some chromosomes show more regions under selection in one lineage or the other. The most extreme example in chimpanzees is chromosome 20, showing 6 times as many genetic regions under selection in the Central/Eastern lineage than in the Western lineage. Less extreme examples are found on chromosomes 8, 9, 13, and 19 with 2-fold more genome space showing evidence of selective sweeps in the Central/Eastern than the Western lineage. In the Western lineage chromosomes 16, 15, 18, 3, and 11 show 5-, 4-, 3-, 2-, and 2-fold more genome space to be under selection than the Central/Eastern lineage, respectively.

While the number of regions under selection in the Western as well as the Central/Eastern lineage was equal on chromosomes 1, 2A, 2B, 4-8, 10-14, and 17, there were some differences in the remaining chromosomes. Chromosomes 20 and 21 in the Central/Eastern lineage had five and four times more regions affected by selective sweeps than the Western lineage. Chromosomes 9, 19, and 22 showed two times more regions. In the Western lineage chromosomes 16 and 15 exhibited four and three times more regions under selection than chromosomes in the Central/Eastern lineage. Chromosomes 3 and 18 showed two times more regions under selection in the Western lineage compared to the Central/Eastern lineage. There was no evidence for selection shared between both lineages on chromosomes 2B, 9, 10, 14-16, and 20, 21-22.

#### Functional annotation, enrichment, and cluster analysis of outlier regions under selection

We analyzed annotated outlier regions with complete or partially overlapping genes and other genetic content (*e.g.*, non-coding genes, pseudogenes). Both lineages had significantly more protein-coding and non-coding genes than randomly sampled genome regions (one sample t-test, *p*=0.0001 & *p*=0.0001) (**Table 2**). Additionally, the number of non-coding genes (ncRNA) was also significantly higher (one sample t-test, *p*=0.0001 & *p*=0.0016), while the number of pseudogenes showed no significant differences (one sample t-test, *p*=0.3023 & *p*=0.6518) (**Table 2**). Closer inspection of the most enriched regions (**Table S2**) revealed these contained mostly protein-coding genes and ncRNAs. The region with the highest significance value in the Western lineage carries exclusively ncRNAs and one window did not contain any annotated genetic features at all.

**Table 2.** Genetic content of 200 kb windows under selection and ten randomly sampled genome regions.

| POPULATION | Real Regions | Random Regions (n=10) |  | p-value<sup>a</sup> |
| --- | --- | --- | --- | --- |
| WEST |  | Average | StDev<sup>b</sup> |  |
| Protein coding genes | 500 | 400.8 | 31.84 | 0.0001 |
| ncRNAs | 178 | 141.1 | 18.60 | 0.0001 |
| Pseudogenes | 5 | 6.2 | 2.90 | 0.3023 |
| CENTRAL/EAST |  |  |  |  |
| Protein coding genes | 536 | 401.1 | 30.65 | 0.0001 |
| ncRNAs | 155 | 136.7 | 24.00 | 0.0016 |
| Pseudogenes | 5 | 6.4 | 2.72 | 0.6518 |

<sup>a</sup>p-values were obtained using the “one sample t-test”.
<sup>b</sup>StDev: Standard Deviation

We examined enriched gene ontology (GO) terms in the ‘Biological Processes’ category (**Table S3**) and enriched KEGG pathways (**Table S4**) for genes under selection in the Western lineage, the Central/Eastern lineage, or shared between the two populations. Genes significantly enriched in the Western lineage are involved in developmental processes (hair follicle development, embryonic development, pattern specification, melanocyte differentiation), cellular and metabolic processes, and protein localization and degradation. Enriched KEGG pathways in the Western lineage were mainly related to diseases caused by pathogens or internal dysfunctions, branched-chain amino acids (BCAAs) degradation, and neurological development. The Central/Eastern lineage genes are enriched for innate immune system response, cellular processes, and wound healing. Enriched KEGG pathways in the Central/Eastern lineage are involved in several diseases affecting the heart muscle and Amoebiasis. The shared dataset showed enrichment only in bone mineralization without any KEGG pathway.

To minimize annotation redundancy and clarify the biological functions in each lineage, we grouped genes into functional clusters based on similar biological meaning, not physical distance [44]. **Fig 2b** and **Table S5** show functional enrichment clusters of genes that were unique to the western group (purple), unique to the central/eastern group (orange) and shared between the western and central/eastern lineage.

We grouped 610 candidate genes from the Western lineage into three clusters. The cluster with the highest enrichment score (ES = 2.3) included four genes (PATR-DOB, PATR-DMB, MAMU-DMA, HLA-DOA) functionally associated with the Major Histocompatibility Complex (MHC) II (**Fig. 3a**). MHC II genes, located on chromosome 6, play a crucial role in the adaptive immune response by activating CD4 T cells to respond to extracellular pathogens [45]. The second cluster (ES = 1.8) contains the same four genes as the first cluster, plus gene HLA-DQA1. This cluster is defined by additional gene functions and displays enrichment in additional disease pathways active in diseases like Asthma, Graft-versus-host disease, Allograft rejection, type I diabetes mellitus, and the intestinal immune network for IgA production [45]. The third cluster (ES 1.58) contains three genes containing the Ephrin receptor-binding domain. These three genes (EFNA4, EFNA3, EFNA1) form a gene cluster on chromosome 1 from position 133,320,040 to 133,391,332. Depending on the context, Eph signaling pathways are key determinants of neurological development, cell morphogenesis, tissue patterning, angiogenesis, and neural plasticity [46, 47].

**Fig 3.**
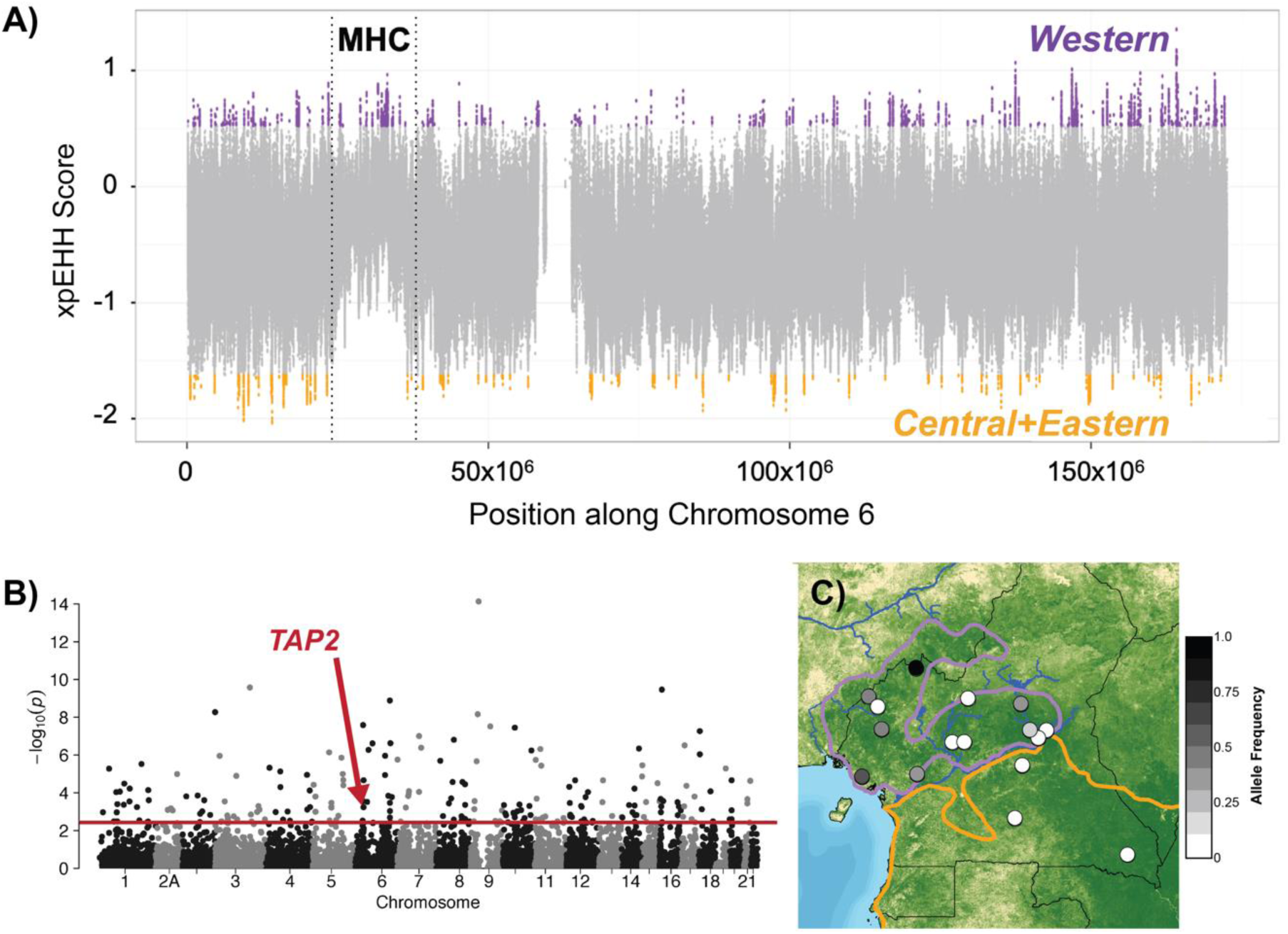
Genome-wide variation of immune response genes under selection in chimpanzees. (A) XP-EHH analysis of SNPs on chromosome 6 from whole genome sequences of captive chimpanzees. Colored points represent SNPs within the 1% tail of the XP-EHH scores across the genome. The entire MHC region is notated, showing SNPs in MHC genes under selection in the Western lineage (*P. t. verus* and *P. t. ellioti*). (B) Manhattan plot shows the genome-wide significance level (solid red line) for SNPs associated with Normalized Difference Vegetation Index (NDVI) - Brown with the *TAP2* SNP noted. (C) Map of allele frequencies for the *TAP2* SNP superimposed onto NDVI and chimpanzee subspecies ranges in Cameroon.

In the Central/Eastern lineage, 593 genes were analyzed, forming three functional clusters. The first (ES=1.8) showed enrichment of three genes (TFF3, TFF2, TFF1) with a PD (or trefoil) domain. These three genes form a cluster on chromosome 21, but their functions are not understood: The peptides coded for in these segments are in several tissues but are most abundant in the GI tract where they may stabilize the mucosa and promote healing [48]. The second cluster (ES=1.7) contained five genes belonging to the beta-defensin gene including DEFB126, DEFB127, DEFB129, and DEFB132 are located on chromosome 20, and DEFB125 on chromosome 8. As antimicrobial peptides they are important in the innate response, including resistance of epithelial surfaces to microbial colonization and encapsulating viruses [49]. The third cluster (ES=1.6) comprises genes INS, RLN3, and INSL6, all sharing an Insulin-like domain.

Functional enrichment analysis of genes shared between both lineages revealed only one cluster of six genes with an enrichment score of 2.2: IL1RL2 and IL18RAP form a gene cluster on chr2A, CNTN6 is located on chr3, and ROBO3, ROBO4, and HEPACAM form a gene cluster on chr11. These genes are all annotated with an Immunoglobulin-like domain.

### Wild chimpanzee SNP genotyping, population structure, and selection analysis

#### Sequence analysis, filtering, SNP calling, and on-target read assessment

We isolated DNA from fecal samples collected non-invasively from unhabituated natural communities of chimpanzees sampled across Cameroon. For 192 of these samples, we obtained 412,081,940 raw reads from single Illumina HiSeq PE125 lane – an average of ~2.15 million reads per sample. In total, 275,443,720 of these reads mapped to the chimpanzee reference genome; from these, we removed approximately 75 million reads and were left with 38,657,083 reads that mapped to our target sites (**Fig. S5a**) – an average of 201,339 on-target reads per sample. The 9,986 targeted sites had a mean read depth of 20x with one site showing as much as 166x coverage (**Fig. S6**). After removing samples for missing data and relatedness, we were left with two datasets (‘10k’ and ‘1k’). The ‘10k dataset’ samples had significantly more on-target reads per sample than the total dataset; an average of 328,863 on-target reads per sample, representing ~16% of the total reads from these samples (**Figs. S5b** and **S5d**). The ‘10k dataset’ filtering process resulted in 7,878 SNPs and 112 samples, and all samples from Boumba Bek (BB) and Campo Ma’an (CP) were removed. To retain more geographic representation of samples from at least one of these sites, we created another dataset (‘1k dataset’) by applying a more stringent site filter and the same individual missingness filter above which resulted in 994 SNPs and 142 samples (including two individuals from CP, but none from BB). The ‘1k dataset’ samples had an average of 268,773 on-target reads per sample, representing ~12% of the total reads from these samples (**Figs. S5b** and **S5f**).

#### Testing for isolation-by-distance and isolation-by-environment

We found that pairwise *F*_ST_ values between sampling locations from different habitats were significantly higher than pairwise *F*_ST_ between sampling locations within the same habitat for both the ‘10k’ (one-tailed *t*-test, *p*-value = 2.2e-16; **Figs. 1c** and **S7a**) and the ‘1k’ dataset (one-tailed *t*-test, *p*-value = 2.1e-16; **Fig. S7b** and **S9a**). Additionally, the geographic distance between sampling locations from different habitats was significantly higher than between locations within the same habitat for all 19 sampling locations included in this study (one-tailed *t*-test, *p*-value = 6.324e-11).

We also performed a permutation test to account for the fact that population structure across habitats can confound the detection of isolation-by-distance (IBD). This categorized population pairs by geographic distance and randomized their habitat origins, forming a null distribution of t-statistics. Using this distribution, we assessed if *F*_ST_ differed more between than within habitats/populations. For the ‘10k’ and ‘1k’ datasets, we found that *F*_ST_ was significantly higher between populations/habitats than within them (*p*-value < 0.0001; **Figs. S8b** and **S9b**). This suggests that IBD alone cannot fully explain the high *F*_ST_ values between populations/habitats. We also ran the permutation test for *P. t. ellioti* sampling locations alone and found that *F*_ST_ is significantly higher between *P. t. ellioti* (Rainforest) and *P. t. ellioti* (Ecotone) than within them compared to the null distribution (*p*-value = 0.0002; **Fig. S10**). Taken together, the results of the permutation tests suggest that habitat differences play a much stronger role than geographic distance alone, although the signal is slightly stronger within *P. t. ellioti* than in *P. t. troglodytes*. This may be attributed to the fact that *P. t. troglodytes* in Cameroon occupies more uniform Congo Basin forested habitat south and east of the Sanaga River. In contrast, *P. t. ellioti* occupies the comparatively diverse Gulf of Guinea forest comprising lowland forest, montane forest, and the forest-savanna gradient north of the Sanaga River.

#### Population structure, hybridization, and demographic history

We next investigated population structure, hybridization, and demographic history. Principal Components Analysis (PCA) (**Figs. S11, S12, S13, S14** and **Table S10**), population clustering analysis results (**Figs. S15** and **S16**), and Analysis of Molecular Variance (AMOVA) (**Table S11**) consistently distinguished between *P. t. ellioti* and *P. t. troglodytes*. The results from wild chimpanzee samples were consistent with results from the genome analysis of captive individuals (**Figs. S17, S18, S19, S20** and **S21**), indicating that our SNP discovery approach from captive individuals is likely capturing pockets of genetic differentiation present in wild individuals. In addition, certain individuals showed hybrid ancestry, notably an F1 hybrid in the *P. t. ellioti* population and a potential backcrossed hybrid in *P. t. troglodytes*. The demographic history model indicates that *P. t. ellioti* and *P. t. troglodytes* split from one another around 478,000 years ago, with continuous but rare gene flow between them since splitting, underlining a complex demographic history characterized by significant admixture and evolutionary divergence within the region. **S1 Text** provides more detailed results from these analyses. Based on these results, we concluded that neither the IBD model nor simple allopatric divergence along the banks of the Sanaga River fully explains the separation of *P. t. ellioti* from *P. t. troglodytes*.

#### Mapping wild chimpanzee genomic variation across habitats

These findings drew our attention to investigating how habitat variation corresponds with neutral and adaptive genetic differentiation among chimpanzees in Cameroon. Using a gradient forest model [50] and 31 environmental predictor variables sourced from publicly available databases (See **Methods**, **SI Text,** and **Table S7**), we quantified environmental associations with genomic loci, pinpointing key environmental drivers and projecting genomic diversity spatially. We identified 581 SNPs with significant environmental associations, representing 6% of all SNPs from wild chimpanzees genotyped in this study. From these, 346 unique candidate genes within 10kb windows of these SNPs, matched outliers from captive chimpanzee genome scans. When mapped to the study, these showed clear signals driven by a phylogenetic split between *P. t. ellioti* and *P. t. troglodytes* across the Sanaga River *<u>and</u>* habitat variation across Cameroon (**Fig. 4c**). Latitude (a proxy for geographic distance) had a pronounced effect along PC1 (**Fig. 4d**). Isothermality and surface moisture also contributed heavily to the model in differentiating between coastal and interior rainforest habitats, as well as rainforest versus ecotone habitats (**Fig. 4c** and **4d**).

**Fig 4.**
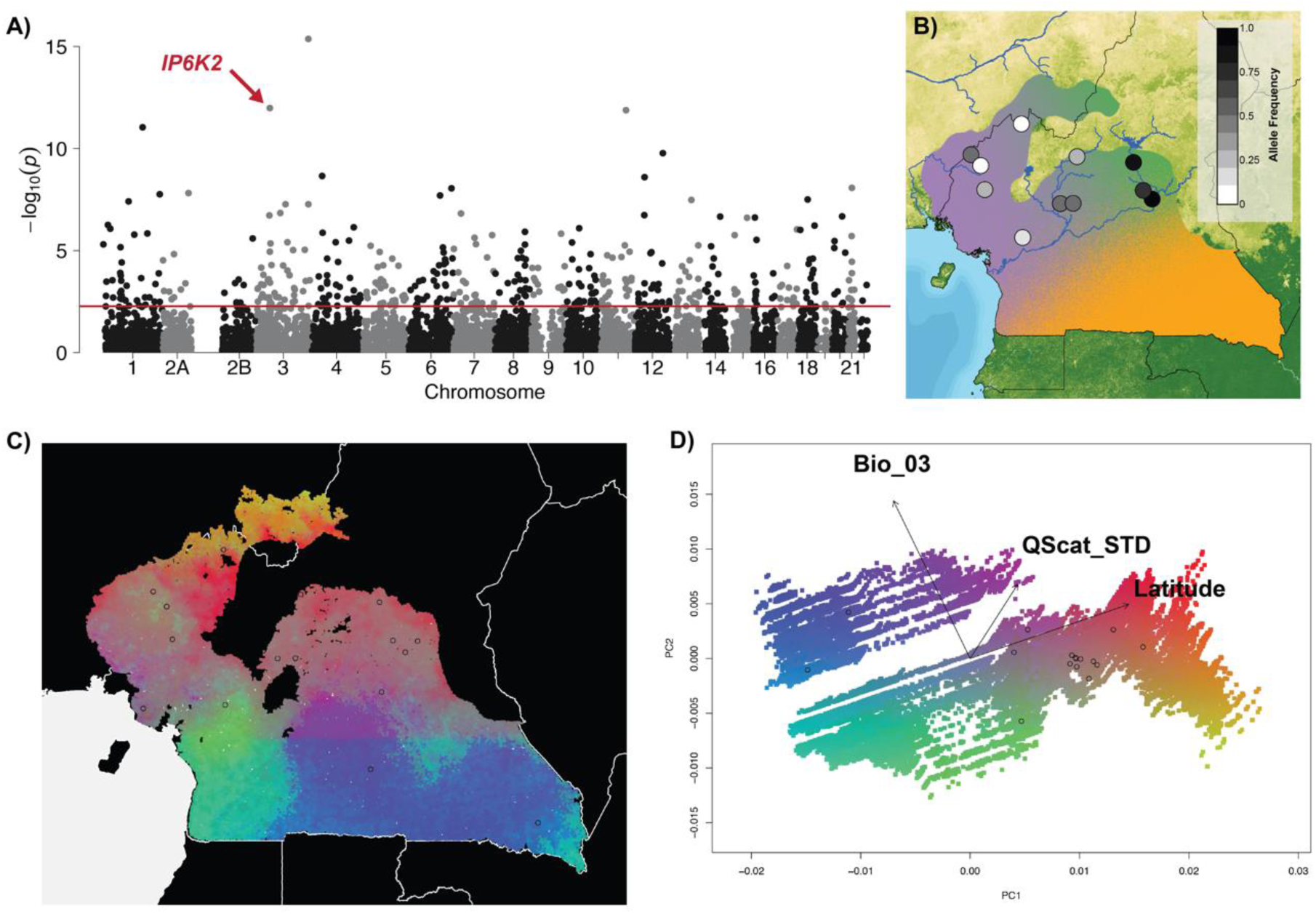
Dietary gene under selection and gene-environment relationships. (A) Manhattan plot shows the genome-wide significance level (solid red line) for SNPs associated with Annual Mean Temperature (BIO1) with the *IP6K2* SNP noted. (B) Spatialized allele frequencies for the *IP6K2* SNP showing differentiation between *P. t. ellioti* populations. (C) Gradient forest-transformed climate variables show climate adaptation across the study area. (D) Colors are based on a PCA of transformed climate variables.

Among all predictors tested in the model, latitude had the highest *R*^2^ weighted importance, likely reflecting the deep split between chimpanzee subspecies and/or bioclimate turnover across the rainforest-savanna gradient. Precipitation during the dry season and vegetation density were also important for predicting chimpanzee genomic diversity (**Fig. S24**). The second most important axis of variation in the gradient forest model primarily contributed to isothermality (bio3) and surface moisture (QScat_STD). Thus, the variables contributing the most align with a rainforest/savanna ecotone split (**Figs. 4c** and **4d**), consistent with previous studies of niche modeling [42].

#### Detecting environmentally associated loci under selection in wild chimpanzee populations

We used Latent Factor Mixed Models (LFMM) to test for signals of selection on individual SNPs in a manner that controls for confounding effects of population structure. We identified 1,690 SNPs significantly associated with one of 31 environmental predictors (**Table S7**) after accounting for population structure (K=3). We then identified 905 unique candidate genes within 10kb windows of the environmentally associated outlier SNPs, all of which were outliers in the captive chimpanzee genomes selection scan. Of the population groupings, we identified 695 associated with General Temperature variables, 388 associated with Temperature Range, 66 associated with Temperature Seasonality, 160 associated with Precipitation (Wet/Cold), 305 associated with Precipitation (Dry/Warm), 456 associated with Surface Moisture, 448 associated with Tree Cover, 355 associated with Vegetation Greenness, 325 associated with Vegetation Brownness, and 428 associated with Topography. A simple Mantel test revealed a significant correlation between pairs of environmental predictor variables and shared outlier SNPs returned by LFMM between variable pairs (Mantel *r* = 0.425, *p* = 1.00e-6) demonstrating that independent LFMM performed as expected.

#### Quantifying environmental relationships with candidate genes

We searched for functional enrichment signals in environment-associated genes in two complementary ways. First, we compared the functions of genes associated with environmental variation with genes that show no signs of positive selection. This comparison helped us determine whether genes influenced by environmental factors and potentially under selection differ functionally from those evolving under neutral conditions. We used genes outside outlier regions from the captive chimpanzee whole-genome analysis as a reference. We identified 47 biological processes enriched in 1,018 unique environmentally associated outlier genes from both the gradient forest and LFMM models (**Table S12**). There were several enrichment clusters, notably two processes functionally associated with immune response, one was related to the Major Histocompatibility Complex (MHC) II – an important part of the adaptive immune system - and eight processes associated with neurological development, including 60 unique genes. We also found 48 enriched KEGG pathways in this subset of outliers (**Table S13**). Key clusters included pathways in neurological development (56 genes), digestion and metabolism (40 genes), and immune response (40 genes).

Our second analysis examined if genes influenced by environmental variation showed functional enrichments compared to those under positive selection without a clear environmental impact. This test uses a much smaller set of background genes composed only of those assayed in wild chimpanzee SNP scan but were not environmentally associated outliers in the LFMM and gradient forest models. Unsurprisingly, the enrichment analysis using this more limited background set of genes resulted in one significantly enriched biological process and KEGG pathway each, both relating to neurological development, specifically axon guidance (**Table S14).**

Of the genes linked with immune response and MHC II, Transporter 2, ATP binding cassette subfamily B member (*TAP2*) stands out. It contains a SNP that significantly associated with Vegetation Brownness (NDVI BRN) (-log_10_ = 3.231762344, p < 0.001) (**Fig. 3b**) and is associated with *General Temperature* variables in the LFMM analysis of wild chimpanzees. The *TAP2* SNP in wild chimpanzees is nearly fixed in *P. t. troglodytes* and is variable across *P. t. ellioti* habitats (**Fig. 3c**). Additionally, *TAP2* was found to be under natural selection in the analysis of captive chimpanzee whole genomes, and it is part of the enriched KEGG pathways under selection in *P. t. verus* and *P. t. ellioti* (**Table S4**). *TAP2* is a component of the transporter associated with antigen processing (TAP) complex, which plays a role in ensuring that MHC class I (MHC-I) molecules are expressed on the cell surface [51]. TAP complex proteins, including TAP2, are essential for viral peptide transport from the cytoplasm onto MHC-I receptors within the endoplasmic reticulum [52]. In humans, several *TAP2* gene variants are linked to an increased HIV-1 infection risk [53].

We identified the gene, Leucine rich repeat and Ig domain containing 2 (*LINGO2*), as having one of the strongest associations with the environmental predictor variable, mean annual normalized vegetation index (NDVI), in the LFMM analysis (-log_10_ = 8.32330639, p = 0.00000000475) (**Fig. 5a**). A linear regression also revealed a significant association between the allele frequencies of the *LINGO2* SNP and mean annual NDVI (R^2^ = 0.4826, p = 0.003495) (**Fig. 5b**). Low allele frequencies were found in *P. t. ellioti* (Rainforest), with variable frequencies found in *P. t. ellioti* (Ecotone) (**Fig. 5c**). *LINGO2* is highly expressed in human brain tissue [54] and affects synapse development and function [55]. *LINGO2* is also under positive selection in Lidia cattle breed subpopulations and partially drives neurobehavioral phenotype variation among them [56].

**Fig 5.**
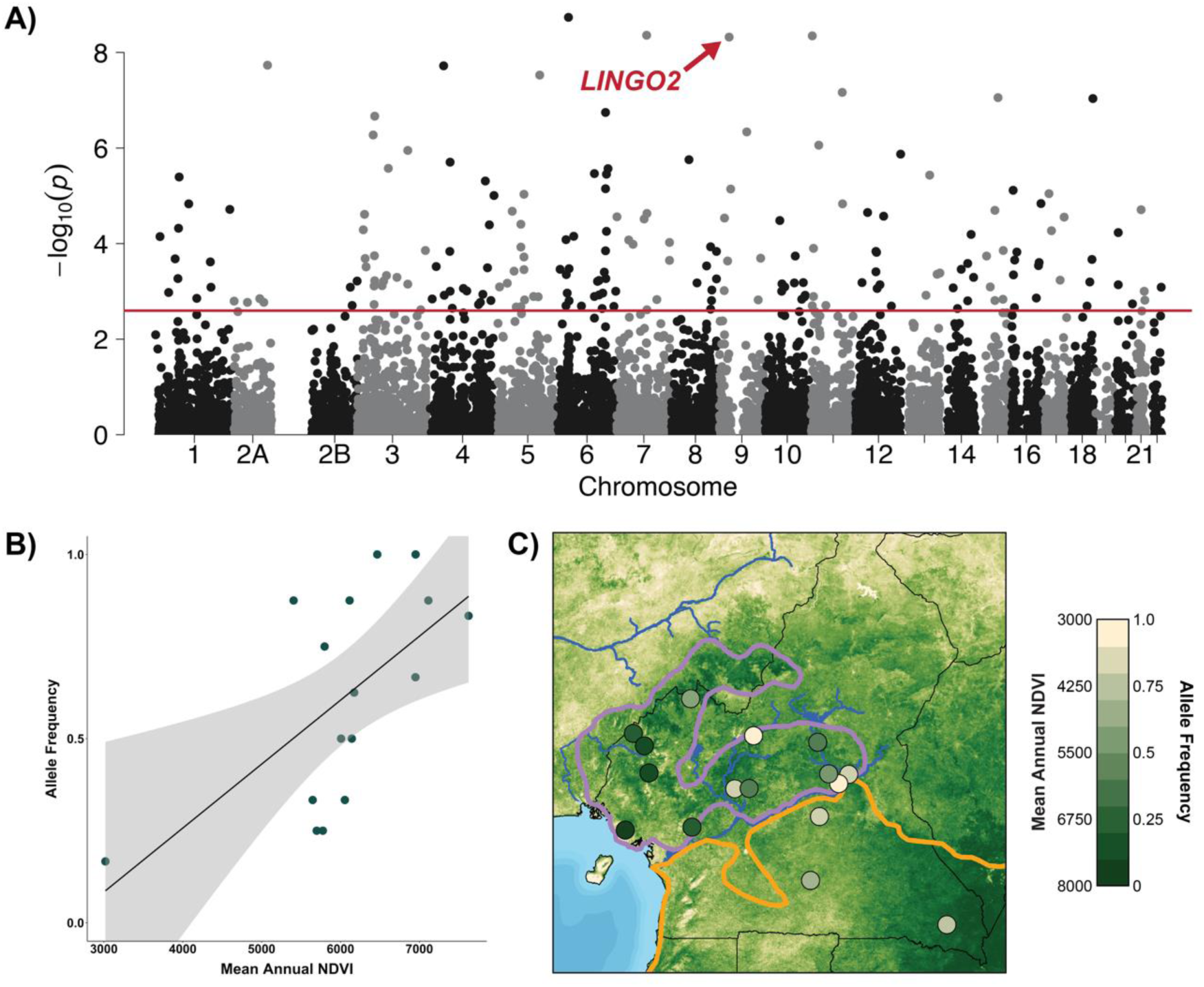
Genome-wide variation of neurological development genes under selection. (A) Manhattan plot shows the genome-wide significance level (solid red line) for SNPs association with mean annual normalized vegetation index (NDVI) with the *LINGO2* SNP noted. (B) Correlation between *LINGO2* allele frequency and mean annual NDVI. (C) Spatialization of allele frequencies for this SNP superimposed onto mean annual NDVI.

Of the 24 digestion and metabolism-related genes identified in the biological processes and KEGG pathways, we further narrowed down our search by using additional measures to quantify relationships of each of the genes with environmental variables and were able to identify two genes with associated SNPs exhibiting significant linear relationships directly with their associated environmental variables across space, suggesting a potential role for diversifying selection across the forest/savanna ecotone gradient. Acetyl-CoA acetyltransferase 2 (*ACAT2*) contains the SNP at position 161,530,902 on chromosome 6 (**Fig. S26a**), which had the strongest association of all SNPs with temperature seasonality according to results of the LFMM analysis (-log_10_ = 3.594887672, p = 0.000254163) (**Fig. S26b**). Linear regression revealed a strong and significant relationship between *ACAT2*’s outlier SNP and temperature seasonality (R^2^ = 0.5651, p = 0.0005) (**Fig. S26c**). When plotting allele frequencies of *ACAT2*’s outlier SNP, higher frequencies were observed in the ecotone’s northern sampling sites. Sampling sites within the range of the *P. t. troglodytes* population had lower frequencies of the allele (**Fig. S26d**). The product of the *ACAT2* gene is known to be involved in cholesterol and beta-oxidation lipid metabolism [57].

Another gene identified to have an environmentally associated outlier SNP was Phospholipase C like 2 (*PLCL2*). *PLCL2* contains the SNP at position 17,268,745 on chromosome 3 (**Fig. S27a**). We identified a strong relationship between this SNP and the environmental variable *precipitation of the wettest month* through LFMM analysis (-log_10_ = 3.36472742, p = 0.00043179) (**Fig. S27b**). Linear regression revealed a strong and significant relationship between *PLCL2*’s outlier SNP and the environmental variable *precipitation of the wettest month* (R^2^ = 0.3422, p = 0.0102) **(Fig. S27c**). We observed higher allele frequencies of the *PLCL2* SNP in the *P. t. ellioti* ecotone population, with the *P. t. ellioti* rainforest population having the lowest frequencies across Cameroon (**Fig. S27d**). *PLCL2* is associated with obesity in mouse models. Individuals lacking the allele were shown to have a leaner phenotype; were able to resist induced obesity due to increased protection from glucose metabolism disorders and insulin resistance; and exhibited higher energy expenditure [58].

Finally, a SNP in the Inositol hexakisphosphate kinase 2 (*IP6K2*) gene was identified as a significant outlier differentiating the *P. t. ellioti* ecotone and rainforest populations and associated with the environmental variable *Mean Annual Temperature* through LFMM analysis (-log_10_ = 11.98296666, p < 0.0001) (**Fig. 4a**). *IP6K2*’s SNP was significantly more divergent than neutral SNPs between the two *P. t. ellioti* populations (*F*_ST_ = 0.49, p < 0.0001) (**Fig. 4b**). In humans, the *IP6K2* gene is linked with inflammatory bowel disease and cellular response to flavonoids, plant metabolites found in fruits and vegetables [59]. The human KEGG pathway containing *IP6K2* is associated with VACTERL/VATER syndrome, often associated with congenital heart disease and chondrodysplasia [60, 61].

## Discussion

We presented genome-wide SNP genotyping from a representative sample of 112 wild chimpanzees from across Cameroon, along with eight newly sequenced genomes of captive chimpanzees to enhance SNP discovery. We supplemented these data by combining genetic analyses with environmental association scans to search for evidence of environmentally-mediated selection. While prior studies have largely concentrated on neutral evolution mechanisms across this contact zone between chimpanzee lineages, our findings support a role for diversifying selection in the divergence of chimpanzee subspecies across different environments. The proposed population history of chimpanzees across this contact zone is well supported in this study. *P. t. ellioti* and *P. t. troglodytes* last shared a common ancestor around 478,000 years ago, with occasional gene flow between them evidenced by an F1 hybrid in *P. t. ellioti* and a potential backcrossed hybrid in *P. t. troglodytes*. These findings support prior studies suggesting that this contact zone between subspecies best fits an isolation-with-migration population model in which allopatric divergence and positive selection contribute to the partitioning of genetic variation [62].

The evidence supporting a role for environmentally-mediated selection across this contact zone is also compelling. We found 1,690 unique SNPs were associated with at least one of 31 environmental predictors, indicating that prevailing environmental conditions contribute to local adaptation in *P. t. elloti* and *P. t. troglodytes*, and to a lesser extent, among populations within *P. t. ellioti*. These SNPs are distributed among 905 outlier genes enriched for 48 biological processes. Overall, the sets of genes with highly divergent allele frequencies that separate *P. t. ellioti* from *P. t. troglodytes* suggest a role for selection in pathways important in two main categories: immune response and life history traits (neurological development, behavior, and dietary function).

It is important to reiterate that all outliers identified in wild chimpanzees using LFMM-based approaches were also identified as outliers in the haplotype homozygosity selection scans of captive chimpanzee genomes. This two-tiered approach offers heightened reliability of selection scans in wild populations while mitigating the incidence of false positives in our final dataset. Moreover, the congruence of these identified genomic regions between the two methods and two complementary datasets suggests that these outliers are subject to positive selection and not merely an artifact of demographic history or neutral population genetic structure.

### Signatures of selection from variable pathogen histories

The lack of natural SIVcpz infection in *P. t. ellioti* sparks interest because it persists despite opportunities for transmission. SIVcpz*Ptt* virus infects *P. t. troglodytes*, crossed the species barrier on at least four occasions: from chimpanzees to humans in southern Cameroon, giving rise to the HIV-1 group M pandemic and to HIV-1 group N [63, 64]. HIV-1 group O and P also arose from transmission from chimpanzees to gorillas before subsequent transmission to humans [65, 66]. Thus, SIVcpz can cross genus boundaries which makes its absence in *P. t. ellioti* particularly striking since this subspecies still exchanges occasional migrants with *P. t. troglodytes*. Finally, the presence of prey primate species that harbor endemic SIV strains also creates multiple pathways for cross-species transmissions [34, 36, 67, 68]. Thus, we speculate that the absence of an SIVcpz in *P. t. ellioti* must be at least partially explained by adaptations that interrupt SIVcpz cell entry and/or boost immune response to clear SIVcpz infection.

Four processes functionally associated with the Major Histocompatibility Complex (MHC) II on chromosome 6 play a crucial role in the adaptive immune response. MHC II peptides stimulate CD4+ T cells that activate downstream immune responses to intracellular pathogens, including viruses. In particular, the Th1/Th2 cell differentiation pathway determines the type of helper cell a CD4+ T cell will become. Naïve CD4+ T cells recognize an MHC class II molecule, activate, and divide to produce clone effector CD4+ T cells specific for a particular antigen. CD4+ T cells can differentiate into T helper type-1 (Th1), T helper type-2 (Th2), or other T helper types, each with distinct cytokine-secretion phenotypes, production of distinct interferons, and different downstream immune responses. This finding corresponds well with a growing body of evidence that positive selection associated with pathogen defenses has contributed to the genetic and phenotypic differentiation of chimpanzee subspecies, especially *P. t. troglodytes* and *P. t schweinfurthii*, which is perhaps due to exposure to different viruses [32].

This finding naturally called our attention to the absence of SIVcpz in *P. t. ellioti* attributed to a lack of gene flow between *P. t. ellioti* and *P. t. troglodytes* [30]. Given that gene flow occurs between *P. t. ellioti* and *P. t. troglodytes*, and that SIVcpz*ptt P. t. troglodytes* is the source of multiple cross-species infections in both humans and gorillas, it is logical to assume that SIVcpz should naturally infect *P. t. ellioti*. We observed evidence of positive selection in *P. t. ellioti* due to highly differentiated SNPs enriched in genic sites. Among these, *TAP2* (**Fig. 3c**) variants increase the risk of HIV infection in humans [53], and may have a similar function in chimpanzees. Given the low level of gene flow, and the absence of sequence data upstream or downstream of the *TAP2* in our data, we cannot conclude whether this is evidence for recent adaptation to SIVcpz*ptt* or evidence of ancient selection resulting from exposure to SIV-like viruses. Evidence is mounting that chimpanzees have had a long and continuing relationship with SIV-like viruses such that differences in viral exposures and immune responses have likely been a central feature of the evolution of chimpanzees [32, 33, 69].

For instance, *P. t. troglodytes* and *P. t. schweinfurthii* also show evidence of recent positive selection in genes involved in SIV/HIV cell entry and immune response to SIV, biological pathways responsible for T-helper cell differentiation, including CD4 [33], and multiple genes that SIV/HIV use to infect and control host cells including CCR3, CCR9 and CXCR6 [32]. There is also compelling evidence of past selective sweeps leading to reduced diversity in the MHC II repertoire of *P. t. verus* that has been attributed to past infections with SIV or SIV-like viruses [70]. Although we cannot speculate further given the nature of the data in this study, our findings add to the mounting evidence that chimpanzees have experienced long-lasting host-virus relationships with SIV-like viruses and that these relationships have been a critical process underpinning their evolution. More detailed investigations are needed on whether the positive selection in *P. t. ellioti* is due to past, recent, or ongoing infection with SIVcpz and/or related viruses.

### Signatures of selection across variable habitats

Cameroon is also a uniquely positioned ‘natural laboratory’ to examine the relative contributions of neutral evolutionary forces versus natural selection in the evolution of many animals, including chimpanzees. In addition to being home to the Sanaga River, a well-known biogeographic boundary for many species, the country is exceptionally ecologically diverse. We speculate that the area is important for understanding how habitat variation and behavioral diversity may impact chimpanzee evolution. The Congo Basin Forest in the south, the Gulf of Guinea Forest in the west, and the Sahelian habitats in the north of Cameroon all converge and interdigitate to form a unique ecotone habitat composed of open woodland, savanna, and riparian forest [71]. This ecotone is a known engine of diversification for many species [72–78]. Differences between *P. t. ellioti* and *P. t. troglodytes* have been linked with habitat variation across Cameroon [62], which suggests a possible role of local adaptation in their genetic differentiation. Finally, there is a further genetic distinction within *P. t. ellioti* itself, with one gene pool associated with the mountainous rainforest in western Cameroon and the other with the ecotone in central Cameroon [24] (**Fig. 1b**). Each gene pool has a unique ecological niche [42, 79] with marked differences in key socioecological variables, including sex-specific differences in community structure [80] and dietary preferences [81].

We observed compelling evidence for positive selection that distinguished *P. t. troglodtyes* from *P. t. ellioti* across this contact zone. We also found evidence of diversifying selection that distinguished *P. t. ellioti* populations that occupy different niches [42], which adds strength to our previous findings that both allopatric divergence due to genetic drift and environmentally-mediated local adaptation contribute to sustaining the prolonged separation of these two subspecies across this narrow contact zone between them. In particular, we found 246 genes involved in cellular, metabolic, and developmental processes were associated with one or more of the 31 environmental predictor variables. Genes with the most divergent allele frequencies were associated with latitudinal variation and separate *P. t. ellioti* from *P. t. troglodytes*. We detected an additional more subtle signal of positive selection among *P. t. ellioti* chimpanzees located in western Cameroon’s mountainous, forested regions compared to the population inhabiting central Cameroon’s drier ecotone forests. Environmentally driven pressures between habitats shape adaptive variation, especially between rainforest and ecotone habitats. Chimpanzees in these two different habitat types were previously identified as distinctive ecological populations occupying unique niches [42, 79]. The two ecological populations also exhibit distinctive differences in diet and nesting preferences [81, 82], key elements of chimpanzee cultural diversity.

Thus, multiple phenotypic axes appear to be under environmentally mediated selection that can be linked to habitat variation and variation in chimpanzee socioecology. Genes under environmentally mediated selection associated with neurological development (e.g., *LINGO2*) could be shaped by selective pressures linked to the development of cultural traits in diverse habitats [37], while those associated with diet and metabolism are likely shaped by pressures related to fruit availability and seasonality [81]. One of the most compelling findings of our study is the identification of 24 genes related to digestion and metabolism with the strongest signals of environmentally-mediated selection, including *ACAT2*, *PLCL2*, and *IP6K2*, which present promising avenues for future research.

This study adds to the emerging evidence that neutral evolutionary forces alone cannot explain the prolonged persistence of the separation of *P. t. ellioti* from *P. t. troglodytes* across the narrow contact zone between them. Local adaptation to prevailing conditions has led to divergence in sets of genes important in immune response, neurological development, behavior, and dietary function. Together, these findings suggest that local adaptation, notably to varying pathogen pressure and different habitat types, has shaped chimpanzee subspecies differentiation in Cameroon, and likely across their broad range. Future studies exploring how these genetic differences map to phenotypic differences in wild populations are needed to better understand precisely which traits — particularly those associated with pathogen defense, diet, social organization, and other aspects of chimpanzee cultural diversity — provide for local adaptation and divergence among chimpanzee populations across this region.

## Materials and Methods

### Captive chimpanzee genomes

#### Sequencing and read mapping

Our captive chimpanzee genomic dataset includes 24 previously sequenced [16] and 8 new chimpanzee genomes from Cameroon, representing all four subspecies: 4 *P. t. verus*, 10 *P. t. ellioti*, 12 *P. t. troglodytes*, and 6 *P. t. schweinfurthii* (**Table S1**). These eight genomes were sequenced using established methods [16] and deposited in GenBank. Details on the samples, the estimated origins of the captive chimpanzees [83], and GenBank accession numbers are in **Fig S1** and **Table S1**. We mapped raw sequencing reads against the chimpanzee reference genome Pan_troglodytes-2.1.4 (*panTro4*; https://www.ncbi.nlm.nih.gov/assembly/GCF_000001515.6/) using BWA-MEM v0.7.12 [84] with default parameters. After removing PCR duplicates using PICARD v1.119 (https://broadinstitute.github.io/picard/index.html), we called variants using FREEBAYES v0.9.20 [85]. After filtering, 12,754,225 high-quality bi-allelic SNPs on the autosomes were retained.

#### Genome scans for signals of selection

We divided SNP datasets into a ‘Western lineage’ (*P. t. verus* & *P. t. ellioti*; *n=*15) and ‘Central/Eastern lineage’ (*P. t. troglodytes* & *P. t. schweinfurthii*; *n=*17). We applied two selection scan methods, cross-population extended haplotype homozygosity (XP-EHH) [86] and integrated haplotype score (iHS) [87] to detect sweeps using hapbin v1.2.0 [88]. Since both tests require haplotypes, we phased the whole-genome SNP datasets (12,754,225 SNPs) with SHAPEIT v2.r837 [89] following established methods [18]. Genetic maps for *panTro4* were provided by de Manuel & Kuhlwilm *et al.* [18] and Auton & Feldel-Alon *et al.* [90]. iHS calculations used SNPs with a minor allele frequency (MAF) over 5%. We determined the ancestral state of each allele using the 6-primate EPO alignment (ftp://ftp.ensembl.org/pub/release-80/fasta/ancestral_alleles/) [91, 92]. After phasing, ancestral allele assignment, and MAF filtering we used 4,577,055 SNPs in the Western lineage and 6,475,338 SNPs in the Central/Eastern lineage for iHS. XP-EHH scores compared both lineages using 12,450,633 SNPs, and the results were normalized across the genome.

#### Defining population-informative neutral SNPs

Using normalized XP-EHH and iHS values, we identified SNPs expected to follow neutral evolution that met the following criteria: (*i*) a *p*-value of > 0.05, (*ii*) absent from the top 1% genomic regions under selection (see *Defining genomic ‘outlier’ regions in captive chimpanzees*), (*iii*) be located >10kb from a gene, and (*iv*) be in linkage equilibrium. Using these parameters, we defined 147,700 neutral SNPs reflecting chimpanzee population structure. We annotated these using the Variant Effect Predictor (VEP) v82 and the UpDownDistance plugin [93].

#### Defining genomic ‘outlier’ regions

To understand the amount of selection on the genome, we considered numbers of base pairs under selection (magnitude) and the size of regions affected (genome space). We employed iHS [87] and XP-EHH [86] to detect signatures of local positive selection. Both assume that long-range haplotypes remain unaffected by recombination, signifying natural selection even with small datasets [43, 94]. They are also complementary: while iHS detects partial sweeps, XP-EHH identifies near-fixation events. Following Pickrell *et al.* [43], chromosomes were split into 100kb non-overlapping windows, and the fraction of SNPs with |iHS| > 2 and the maximum XP-EHH was used as a test statistic. We analyzed the fraction of SNPs with |iHS| > 2 and the maximum XP-EHH per window. We turned these into empirical *p*-values by binning windows by SNP count, with iHS dropping windows with < 100 SNPs. Each window’s statistic value was compared against others in its bin to determine an empirical p-value. All bins were then sorted by this *p*-value. The top 1% of each test statistic was noted. ‘Outlier regions’ were windows in this 1% (*p*-value < 0.01). Adjacent windows were merged, retaining the smallest *p*-value.

#### Characterizing genomic regions under selection

As XP-EHH and iHS are complementary, we analyzed the 1% tail of each test merging adjacent windows. Windows were extended by 50kb on either side and annotated for gene content using Ensembl’s BioMart [95], including protein-coding genes, pseudogenes, and RNA coding genes. Genes within outlier regions were considered candidate genes. We tested whether these genomic regions carry certain types of gene content more often than expected by chance by randomly selecting regions equivalent in length and annotating them as described above. We repeated this process 10 times for the Western and Central/Eastern populations, respectively. We counted the number of protein coding genes, non-coding RNAs, and pseudogenes in the real and randomized datasets. We then calculated mean and standard deviation for each and performed a one-sample t-test to determine significance.

#### Functional annotation and enrichment analysis of whole-genome datasets

We used DAVID Bioinformatics Resources v6.8 [44] to annotate candidate genes and perform an enrichment analysis with default functional annotations (GO terms, KEGG pathways, protein domains). We concentrated on the ‘functional annotation clustering’ function using the highest classification stringency and adjusted the enrichment thresholds for EASE to 0.05, reducing non-significant term inclusion. This clustering reduces redundancy by grouping similar annotations. Clusters received a Group Enrichment Score based on their *p*-value, ranking their biological importance. High scores likely mean lower *p*-values for annotation members [44]. We omitted windows found in both Western and Central/Eastern lineages, analyzing them separately. We set a background gene population as the entire chimpanzee genome, as recommended for genome-wide studies [44].

### Wild chimpanzee SNP genotyping, population history, and selection analysis

#### Fecal sample collection, DNA extraction, and quantification

We sampled wild, non-habituated chimpanzee populations using non-invasive methods during a series of field studies from 2003 to 2015 spanning remote forested regions of Cameroon (**Fig. 1b**). Sampling occurred in protected and unprotected areas, as detailed in **Table S6.** Chimpanzee fecal samples were collected and stored following established protocols [24]. All samples were transported from Cameroon to the United States in full compliance with the Convention of International Trade in Endangered Species of Wild Fauna and Flora (CITES), the Centers for Disease Control (CDC) export and import regulations, and with approval from the Government of Cameroon.

Following established protocols [24], DNA was extracted from fecal samples with the QIAamp DNA Stool Mini Kit (Qiagen). Due to the low proportion of endogenous DNA in fecal gDNA extracts [96, 97], samples were sometimes extracted up to six times to ensure enough chimpanzee DNA for sequencing processes. The concentration of endogenous DNA was measured via quantitative real-time PCR using the Quantifiler™ Human DNA Quantification Kit (Applied Biosystems) and methods from prior studies [24].

#### SNP ascertainment, library preparation, DNA capture enrichment, and sequencing

We genotyped 9,986 SNPs of wild chimpanzees from Cameroon, chosen from the larger set of 12,450,633 SNPs identified in the captive chimpanzee genome dataset. This selection comprised: (i) population informative neutral SNPs (n=3,492) randomly selected 147,700 neutral SNPs defined in the whole-genome dataset from above; (ii) ‘outlier’ SNPs (n=6,494) identified through iHS or xpEHH tests as being in the top 1% for selection signals and within or 10k bp up- or down-stream of a known gene; and, (iii) SNPs in genes involved in immune response, disease resistance, and dietary adaptation in humans (n=20) [98, 99]. For each targeted SNP, we designed two 80 nucleotide biotinylated RNA probes, overlapping by 20bp, to create 100bp windows around each SNP using the panTro4 chimpanzee reference genome. After rigorous filtration using the Arbor Biosciences BLAST pipeline, we finalized a bait-set of 19,974 probes, assigning one or two probes to each SNP based on the outcome of the stringent filtering process.

gDNA samples were prepared in clean facilities at Arbor Biosciences to prevent contamination. DNA was quantified, sonicated, and size-selected for around 300nt fragment lengths. Samples were converted to sequencing libraries via adapter ligation. They were index-amplified based on the DNA input amount. Up to 2μg of each library was then enriched using the myBaits system v3. Different enrichment and amplification protocols were applied depending on the DNA quantity in the starting extract. Libraries were combined for equal representation, sequenced on an Illumina HiSeq PE125 lane at HudsonAlpha, and protocols were consistent with studies on degraded or low endogenous DNA samples [96, 97].

#### SNP calling and on-target read assessment

We filtered sequence reads with the Illumina CASAVA-1.8 FASTQ Filter (http://cancan.cshl.edu/labmembers/gordon/fastq_illumina_filter/) and mapped them to the chimpanzee panTro4 genome using BWA-MEM. After removing PCR duplicates with PICARD, we called variants using FREEBAYES. We evaluated DNA capture enrichment and sequencing for the raw and filtered sequence reads using SAMtools v1.3.1 (101) and VCFtools v0.1.15 [100] following established methods (97) for all 192 individuals. Using VCFtools, we filtered variant calls based on quality and coverage. SNPs with <5x coverage or quality score <30 were recorded as missing data [97]. Positions with >60% missing data, a minor allele frequency below 5%, or individuals with >70% missing data were removed. This resulted in 7,878 SNPs from 112 samples, termed the ‘10k dataset’. Due to removing all Boumba Bek (BB) and Campo Ma’an (CP) samples, a second ‘1k dataset’ was made with stricter site filtering, yielding 994 SNPs and 142 samples, which included two from CP but none from BB. We also removed closely related and duplicate samples using the R package related [101] using the triadic likelihood method [102], resulting in 85 individuals in the ‘10k dataset’ and 108 individuals in the ‘1k dataset.’

#### Testing for isolation-by-environment and inferring population structure

**S1 Text** provides full details on methods to test IDB, IBE and to infer population structure, hybridization, and demographic history. In brief, we examined IDB versus IBE using pairwise F_ST_ values between sampling locations using Arlequin v3.5 [103], while geographic distances were determined with the geosphere package in R [104], focusing on areas with more than one individual. Population structure was inferred by PCA and ADMIXTURE analysis [105], DISTRUCT v1.1 [106], CLUMPP v1.1.2 [107], with geographic without pre-assigned population labels using the SNPrelate package in R, focusing on ‘neutral’ SNPs identified from captive chimpanzee genomes. We mapped genetic clusters using TESS [108] and Ad-Mixer v1.0 [109], accounting for IDB. We calculated observed and expected heterozygosity using the adegenet package in R [110], identified potential hybrids using NEWHYBRIDS v1.0 [111] as implemented in the R packages *hybriddetective* [112] and *parallelnewhybrid* [113], and investigated demographic history using δaδi [114] to model asymmetric migration patterns between *P. t. ellioti* and *P. t. troglodytes*.

Environmental data layers (**Table S7**) were compiled and analyzed to assess habitat suitability and IBE for chimpanzees in Cameroon and Nigeria. These layers, sourced from publicly available databases, included diverse variables such as topography, hydrography, climate, vegetation, moisture content, and tree cover. After standardizing these layers to a 30-arcsecond resolution and converting them to the WGS84 coordinate system, the dataset underwent cross-correlation analysis to pinpoint environmental factors significantly influencing chimpanzee distribution (**Table S8**).

#### Mapping genomic variation across habitats

Using the R package *gradientForest* [50], we calculated associations between allele frequencies and environmental variation across suitable habitat. This extended random forest model identifies links between response variables (e.g., SNP allele frequencies) and spatial environmental factors [115] by iteratively processing datasets, assessing outliers and predictor significance. Gradient forests further apply regression to multiple responses, revealing genomic variation from environmental shifts. This can pinpoint areas of high intraspecific variation, subspecies transitions, or barriers separating genomic variation related to the environment [116]. Following established methods, we refined the environmental dataset (**Table S7**) to reduce noise and applied the gradient forest model [76].

We ran gradient forests on 7,878 SNP allele frequencies used as a response dataset, and 17 environmental variables as the predictors in our final model, including measures of temperature, precipitation, vegetation, surface moisture, and geographic features at the sampling locations. We ran 100 trees in our model, noting SNPs significantly associated with any environmental variable (R^2^ > 0) and the average regression of all associated SNPs. To assess model performance, we randomized the environmental data and ran 200 permutations of the model, creating a distribution of R^2^ and significant SNP associations. We then ran 200 permutations of the actual model, comparing these distributions. (**Fig. S25**).

#### Detecting environmentally associated loci under selection

In order to understand the degree to which environmentally driven natural selection may cause chimpanzees to be locally adapted to different habitats, we used latent factor mixed models implemented in the program LFMM v1.5 [117]. LFMM quantifies statistical associations between allele frequencies and environmental variables, accounts for underlying population genetic structure, and detects loci with stronger environmental correlations than population structure. We ran five MCMC replicates for all environmental variable with 25,000 burn-in steps, 100,000 iterations, and a latent factor of *K*=3 from *a priori* knowledge of wild chimpanzee population structure in Cameroon (**Figs. S11, S12, S13, S14, S15, S16, S20** and **S21**). We calculated median z-scores across runs and used them to calculate the genomic inflation factor (λ) and adjusted *p*-values. To correct for multiple testing, we applied a conservative false discovery rate (FDR) of 0.1 using the Benjamini-Hochberg algorithm. We identified unique candidate SNPs linked to at least one environmental variable, presented via Manhattan plots using the *qqman* package [118].

Using outlier analysis with LFMM, we grouped highly correlated environmental variables together to create ‘environmental groupings’ (**Table S9**). These included: General Temperature (n=7), Temperature Range (n=2), Temperature Seasonality (n=2), Precipitation – Wet/Cold (n=4), Precipitation – Dry/Warm (n=4), Tree Cover (n=2), Vegetation Brownness (n=2), Vegetation Greenness (n=3), Surface Moisture Content (n=2), and Topography (n=3).To determine the degree to which types of environmental variation may drive selection of different genomic regions, we identified panels of unique candidates from each ‘environmental grouping.’ We also analyzed the impact of multicollinearity of our environmental predictors on the LFMM results by correlating the degree of association between pairs of environmental predictors and the number of shared outlier SNPs, using a Mantel test in R. (**Table S8**).

#### Enriched gene ontologies and KEGG pathways

We identified candidate genes near candidate SNPs positions with Ensembl’s BioMart tool [119, 120], including both complete and partial genes within these windows. We used the DAVID database [44] for annotation and enrichment analysis of candidate gene lists focusing on the ‘Biological Processes’ category of the ‘Gene Ontology’ database [121] and KEGG pathways [60] using a *p*-value threshold of 0.05 and two different background populations of genes to control for potential bias since the SNPs assayed in wild chimpanzees were selected from a subset of those identified using whole-genome data from captive individuals. The first background we used was composed of the population of genes found outside regions under selection identified in the whole-genome sequencing data (**Fig 2a**). This resulted in a broad view of environmentally mediated selection in wild chimpanzees by including only genes in putatively neutral regions of the genome. The second background population of genes consisted of all genes assayed in wild chimpanzees, excluding environmental outliers.

## Acknowledgments

We thank the government of Cameroon for permission to conduct this research. We thank the Congo Basin Institute, the International Institute for Tropical Agriculture, the Cameroon Biodiversity Association, the San Diego Zoo Wildlife Alliance, the Wildlife Conservation Society, and the World Wildlife Fund for their support in Cameroon. We thank Kevin Njabo, Beatrice Hahn, Martine Peeters, Louis Nkembi, and Amy Pokempner for their assistance in collecting fecal samples in Cameroon. We thank Felice Elefant for the use of her lab’s 7500 Real-Time PCR System. We thank Alison Devault and Jacob Enk of Arbor Biosciences for assistance in the experimental design and execution of the wild chimpanzee DNA capture enrichment and sequencing as part of Arbor Biosciences’ myReads services.

## Supporting Information

**S1 File. Supplemental Results.**

**S2. File Extended Methods and Materials.**

**S1 Fig. Origins of chimpanzees of Cameroon included in this study.** Sample locations and proportions of estimated ancestry were estimated in previous studies [83, 122].

**Table S1. Captive chimpanzee genomes included in this study**

**S2 Fig. Heterozygosity estimates of captive chimpanzee genomes.**

A. Individual heterozygosity.
B. Subspecies heterozygosity.

**S3 Fig. Population structure of captive chimpanzee genomes**.

A. PCA of LD pruned SNP data set consisting of 1,113,142 SNPs.
B. sNMF individual ancestry analysis of the LD pruned data set in a range of K values.
C. PCA of Neutral SNP data set consisting of 147,000 SNPs.
D. sNMF individual ancestry analysis of the neutral SNP data set in a range of K values.

**S4 Fig. Cross Entropy Results.**

Value of the cross-entropy criterion as a function of the number of ancestral populations in sNMF for (A) the LD pruned SNP panel and (B) the neutral SNP panel.

**Table S2. Top 10 regions under selection including their genetic content.**

**Table S3. Enriched GO terms in the “Biological Processes” domain.**

**Table S4. Enriched KEGG pathways.**

**Table S5. Functional enrichment clustering.**

**Table S6. Number of wild chimpanzee samples collected and used in this study.**

**Table S7. Environmental predictor variables used characterize chimpanzee habitats.**

**Table S8. Pearson correlation table of environmental variables.**

**Table S9. Environmental variable groupings.**

**S5 Fig. Illumina reads of wild chimpanzee samples evaluated into 4 categories.**

Raw reads – yellow, mapped reads (sequence reads that mapped to the panTro4 reference genome) – green, mapped and deduplicated reads (PCR duplicates were removed) – light blue, and on-target reads mapped to our sites – dark blue.

A. Proportion of read types for all 192 sequenced chimpanzee samples.
B. Number of read types for 85 samples included in the ‘10k’ dataset.
C. Number of read types for samples removed due to missingness (>30% missing sites).
D. Proportion of read types for 85 samples included in the ‘complete’ dataset.
E. Number of read types for duplicate samples removed following relatedness analysis.
F. Number of read types for 23 individuals included in the ‘1k’ dataset, but not the ‘10k’ dataset.

**S6 Fig. Frequency of mean sequencing coverage of wild chimpanzee samples for each site.**

Mean coverage across all sites was 20.2 reads/site. The red vertical line represents the minimum coverage needed to accurately call SNPs (5x coverage).

**S7 Fig. Pairwise *F*_ST_ between sites shows population structure. (A) ‘10k’ dataset. (B) ‘1k’ dataset.**

**S8 Fig. Isolation-by-distance for ‘10k’ dataset.**

A. Correlation between ‘linearized *F*_ST_’ and geographic distance (km) generated using the ‘10k’ dataset. Solid circles represent pairs of sampling locations from the same habitat. Dual-colored diamonds represent pairs of sampling locations from different habitats.
B. Null distribution of t-statistics from 10,000 permutations same- or different habitat/population pairs in four bins of geographic distance. The red dotted line shows the t-statistic value for actual data.

**S9 Fig. Isolation-by-distance for ‘1k’ dataset.**

A. Correlation between ‘linearized *F*_ST_’ and geographic distance (km) generated using the ‘1k’ dataset. Solid circles represent pairs of sampling locations from the same habitat. Dual-colored diamonds represent pairs of sampling locations from different habitats.
B. Null distribution of *t*-statistics from 10,000 permutations same- or different habitat/population pairs in four bins of geographic distance. The red dotted line shows the *t*-statistic value for actual data.

**S10 Fig. Isolation-by-distance for only *P. t. ellioti* populations.**

A. Correlation between ‘linearized *F*_ST_’ and geographic distance (km) generated using the ‘10k’ dataset. Solid circles represent pairs of sampling locations from the same habitat. Dual-colored diamonds represent pairs of sampling locations from different habitats.
B. Null distribution of *t*-statistics from 10,000 permutations same- or different habitat/population pairs in four bins of geographic distance. The red dotted line shows the *t*-statistic value for actual data.

**S11 Fig. PCA of all SNPs from the ‘10k’ dataset.**

The first two principal components recapitulate known population structure of chimpanzees in Cameroon. They show 3 clear populations, and one *P. t. ellioti* (Ecotone) individual (CMMD06) clustering with *P. t. troglodytes*, as well as multiple *P. t. ellioti* (Rainforest) clustering together with *P. t. ellioti* (Ecotone) and vice versa.

**S12 Fig. PCA of only neutral SNPs from the ‘10k’ dataset.**

**S13 Fig. PCA of all SNPs from the ‘1k’ dataset**

**S14 Fig. PCA of only neutral SNPs from the ‘1k’ dataset**

**Table S10. The results of the Tracy-Widom test for all SNPs from the ‘10k’ SNP dataset**.

**S15 Fig. ADMIXTURE bar plots for *K*=2-3.**

These three populations correspond to known population structure. However, at *K*=2, there is a signal of possible historic gene flow of *P. t. troglodytes* into *P. t. ellioti* (Ecotone). Moreover, the is one individual (CMMD06 – also identified in the PCA) as being a potential *ellioti/troglodytes* hybrid. At *K*=3, we see evidence of three additional individuals that may be Rainforest/Ecotone hybrids, as well as evidence of mixing between the populations.

**S16 Fig. The cross-validation error results of ADMIXTURE analysis of wild chimpanzees (‘10k’ dataset).**

**S17 Fig. PCA results of the merged captive and wild datasets.**

**S18 Fig. ADMIXTURE bar plots for *K*=2-5 for merged captive and wild datasets.**

**S19 Fig. The cross-validation error results of ADMIXTURE analysis of merged captive and wild chimpanzees (‘10k’ dataset).**

**S20 Fig. Cluster analysis and spatial interpolation of population structure.**

A. TESS bar plots showing individual proportions of ancestry of wild chimpanzees.
B. Spatial interpolation of the *Q* matrix for *K*=3 generated using TESS and Ad-Mixer.

**S21 Fig. Estimating *K*_MAX_ from TESS analysis.**

D*K* values estimated for *K*=1-5 across 10 replicate runs.

**Table S11. Analysis of Molecular Variance (AMOVA).**

**S22 Fig. Mean observed heterozygosity.**

A. Heterozygosity of all loci for all individuals grouped by population.
B. Heterozygosity for all individuals.

There were no significant differences between heterozygosity for each population.

**S23 Fig. Posterior plots of model performance.**

A. The observed Joint SFS for *P. t. troglodytes* and *P. t. ellioti*.
B. the simulated Joint SFS for *P. t. troglodytes* and *P. t. ellioti* under the most likely asymmetric migration scenario obtained from *δaδi*.
C. The residuals between the modeled and observed Joint SFS.
D. A 1D histogram of the residual values between the model and the observed data

**S24 Fig. *R*^2^ weighted importance of the environmental predictor variables to the Gradient Forest model of gene-environment relationships.**

**S25 Fig. Results of randomized gradient forest models (n=200), as compared to results from the observed data (n=200).**

An average 588 of 7,878 SNPs demonstrated a positive R^2^ with at least one environmental variable, with an average R^2^ = 0.155 in the Observed data distribution (n=200, represented by the red histograms in A) and B) above). A significantly different average was obtained when randomizing the associations between the genomic data (SNPs) and environmental predictors for both total SNPs with a positive R^2^ (average total = 504, t = 5.011(unequal variances df = 202.28), p < 0.0001), as well as for the average R^2^ (average = 0.152, t = 2.806 (unequal variance df = 261.97), p = 0.0054) of the randomized gradient forests runs (n=200).

**Table S12. Enriched GO terms in the ‘Biological Processes’ domain for environmentally associated outliers (LFMM and gradient forest) in wild chimpanzees**.

**Table S13. Enriched KEGG pathways for environmentally associated outliers (LFMM and gradient forest) in wild chimpanzees.**

**Table S14. Enriched GO terms in the ‘Biological Processes’ domain for environmentally associated outliers (LFMM and gradient forest) in wild chimpanzees.**

**S26 Fig. Evidence of selective pressures on acetyl-CoA acetyltransferase 2 (*ACAT2*).**

A. Map of the *ACAT2* gene on chromosome 6 with brown star representing the SNP identified through outlier analysis between *P. t. troglodytes* and *P. t. ellioti*.
B. Manhattan plot showing the significance (as the negative log_10_ p-value) of SNP associations with the environmental variable temperature seasonality. Grey colors distinguish different chromosomes. The red line represents the threshold for significant association (p = 0.05). The SNP contained in the ACAT2 gene is highlighted by the red arrow.
C. Correlation between the allele frequency of the SNP contained in the ACAT2 gene and temperature seasonality values at each corresponding sampling location (R^2^ = 0.5615, p = 0.0005).
D. Allele frequencies of the SNP contained in the ACAT2 gene across Cameroon. Sampling sites are represented by circles that are shaded according to the frequency of the allele within the population. SNP frequencies are plotted against temperature seasonality across the region.

**S27 Fig. Evidence of selective pressures on phospholipase C like 2 (*PLCL2*).**

A. Map of the PLCL2 gene on chromosome 3 with brown star representing the SNP identified through outlier analysis between P. t. troglodytes and P. t. ellioti.
B. Manhattan plot showing the significance (as the negative log10 p-value) of SNP associations with the environmental variable precipitation of wettest month. Grey colors distinguish different chromosomes. The red line represents the threshold for significant association (p = 0.05). The SNP contained in the PLCL2 gene is highlighted by the red arrow.
C. Correlation between the allele frequency of the SNP contained in the PLCL2 gene and precipitation of wettest month values at each corresponding sampling location (R2 = 0.3422, p = 0.0102).
D. (D). Allele frequencies of the SNP contained in the PLCL2 gene across Cameroon. Sampling sites are represented by circles that are shaded according to the frequency of the allele within the population. SNP frequencies are plotted against precipitation of wettest month across the region.

