## Supplemental Information for "Environmentally-mediated selection parallels population divergence across a chimpanzee subspecies contact zone"

#### Short title
Local adaptation in wild chimpanzees

#### Authors
Matthew W. Mitchell[1,2,*], Walker Alexander[3], Dana V. Mitchell[1], Adam H. Freedman[4], Janina Dordel[1], Ryan J. Harrigan[5], Ahmet Sacan[3], Fabrice Kentatchime[1,6], Bryan S. Featherstone[1], Ekwoge E. Abwe[1,7,8], Paul R. Sesink Clee[1], Abwe E. Abwe[8], Sabrina Locatelli[9], Bethan J. Morgan[7,8,10], Bernard Fosso[11], Roger Fotso[11], Sarah A. Tishkoff[12], Evan E. Eichler[13,14], Nicola M. Anthony[15], Thomas B. Smith[5,16], Mary Katherine Gonder[1,6,*]

**This file contains all supplemental text (results and methods), figures, and tables referenced in this manuscript.**

### Supplemental Results

#### Captive chimpanzee genome analysis and SNP discovery

##### Genetic diversity and population structure

We calculated heterozygosity estimates to assess genetic diversity within and between subspecies on an individual basis (**Fig. S2a**) and then summarized the estimates into subspecies values (**Fig. S2b**). The overall pattern of subspecies heterozygosity rates confirm previously reported estimates [1, 2] with *P. t. verus* showing the lowest heterozygosity (0.68), followed by *P. t. ellioti* and *P. t. schweinfurthii* with almost equal rates (1.29 and 1.35), and *P. t. troglodytes* with the highest estimate (1.71). These heterozygosity values were calculated using a different methodology, and as a result the values themselves are inconsistent with previously reported estimates [1, 2], although the pattern remains the same. Two individuals, Nicolene and Ewake, show unusual high heterozygosity rates (2.86 and 2.35, respectively), suggesting recent hybridization between *P. t. ellioti* and *P. t. troglodytes*.

Principal components analyses (PCAs) of the LD-pruned panel (**Fig. S3a**) and the neutral SNP panel (**Fig. S3c**) reveal continental population structure consistent with previous studies. The first principal component (PC) of both PCAs separates the Western lineage (*P. t. verus* and *P. t. ellioti*) from the Central/Eastern lineage (*P. t. troglodytes* and *P. t. schweinfurthii*) while the second PC separates *P. t. verus* from *P. t. ellioti*. Several individuals show some degree of genetic distinction compared to other individual of their subspecies. For *P. t. ellioti* these individuals are Tobi, Julie, Julie_LWC21, and Banyo confirming what has been shown in previous studies that these individuals are consistently distinctive from other chimpanzees of their subspecies suggesting the unsampled or under-sampled demes might exist in Cameroon [1-3]. The newly sequenced individual, Nicolene, however, stands out and is found right *between* *P. t. ellioti* and *P. t. troglodytes* clusters and is separated from *P. t. ellioti* by the third PC. Within *P. t. troglodytes* two samples show some degree of divergence: Kita and Doris. Kita is separated from the other *P. t. troglodytes* individuals by the second PC.

The results from the sNMF cluster analysis performed for *K* = 2-5 using the LD-pruned (**Fig. S3b**) and the neutral SNP panels (**Fig. S3d**), respectively are largely consistent with the pattern observed in the PCAs. *K* = 2 separated the two main lineages: the Western lineage (*P. t. verus* and *P. t. ellioti*) and the Central/Eastern lineage (*P. t. troglodytes* and *P. t. schweinfurthii*). At *K* = 3, *P. t. verus* from *P. t. ellioti* separate from one another, and the cross-entropy results indicate that *K* = 3 was the best fit for the data (**Fig. S4**). At *K* = 4, *P. t. troglodytes* from *P. t. schweinfurthii* separate from one another. *K* = 5 revealed known population substructure among *P. t. ellioti* chimpanzees from central Cameroon, north of the Sanaga River [4], but given the cross-entropy results, the population structure revealed at *K* = 4 and 5 may be indicative of oversplitting. These unique individuals, along with other *P. t. ellioti* individuals, showed high amounts of *P. t. troglodytes* ancestry (in contrast to *P. t. verus*) chimpanzees, which reflects the known gene flow between the Nigeria-Cameroon chimpanzees with the central/eastern chimpanzee lineage.

As described by Prado-Martinez et al. [1] and de Manuel et al. [2], Julie, Banyo, and Tobi, as well as the newly sequences Nicolene and Kita show evidence of admixture (**Figs. S1, S3b, and S3d**). Nicolene likely originated from central Cameroon north of the Sanaga River (**Fig. S1**) and shows evidence of admixture between *P. t.*

*ellioti* and *P. t. troglodytes*. Kita likely originated south of the Sanaga River with primary ancestry shared with *P. t. troglodytes* individuals (**Fig. S1**) but with a low level of admixture with *P. t. ellioti*. However, at K = 5, these individuals cluster together as a fifth population (**Figs. S3b & S3d**).

### Wild chimpanzee SNP genotyping, population structure, and selection analysis

#### Population structure

Principal Component sAnalysis (PCA) of the ‘10k’ dataset (all SNPs – **Fig. S11** and neutral only – **Fig. S12**) and the ‘1k’ dataset (all SNPs – **Fig. S13** and neutral only – **Fig. S14**) were consistent with one another as well as the patterns of known population structure of chimpanzees in Cameroon. A Tracy-Widom test classified the first seven eigenvectors as significant ($p < 0.05$) for both the ‘10k’ dataset (**Table S10**), although only the first two are shown in **Figs. S11-14**, as they capture the majority of the variation. The first principal component (PC) for all PCAs generally separates *P. t. ellioti* from *P. t. troglodytes*, with one notable exception. One individual originating from the ecotone region in central Cameroon north of the Sanaga River (CMMD06) is an outlier and clusters closely to *P. t. troglodytes*, indicating that it might have recent hybrid ancestry. The second PC separates *P. t. ellioti* (Rainforest) from *P. t. ellioti* (Ecotone) clusters from one another, although there are several individuals who cluster with a different population (CMEB31, CMEB46, and CMYB03), indicating admixture between the two *P. t. ellioti* sub-populations. These ‘outlier’ individuals are consistent across all PCAs. Within the *P. t. ellioti* (Ecotone) cluster there are several individuals from

Batanga (KM) and Wouchaba (WC) with some degree of divergence from the rest of the cluster, although other individuals from these sampling locations cluster together with the other *P. t. ellioti* (Ecotone) individuals. Batanga and Wouchaba are located in the remote south-east of the *P. t. ellioti* (Ecotone) range (**Fig. 1b**).

**Fig S15** shows the results from the ADMIXTURE cluster analysis for $K$ = 2 and $K$ = 3 using the ‘10k’ neutral SNP panel. These results are consistent with the PCAs and known population structure of wild chimpanzees in Cameroon. The cluster analysis for $K$ = 2 separated *P. t. ellioti* and *P. t. troglodytes*, with all individuals originating from the *P. t. ellioti* (Ecotone) range showing at least some degree of *P. t. troglodytes* ancestry, while one individual (CMMD06) shows ~70% *P. t. troglodytes* ancestry. At $K$ = 3, *P. t. ellioti* is sub-divided into Rainforest and Ecotone populations, with three individuals showing ancestry into different populations (CMEB31, CMEB46, and CMYB03), consistent with the PCAs. The results from the cross-validation error values for all runs indicate that $K_{MAX}$ = 3 and is the best fit for the data (**Fig. S16**). These results indicate population structure into known regions with both historic admixture between the subspecies, as well as recent hybridization between *P. t. ellioti* and *P. t. troglodytes,* as well as *P. t. ellioti* (Rainforest) and *P. t. ellioti* (Ecotone).

The results of the cluster analyses on the merged captive and wild chimpanzee dataset are consistent with known population history and structure of chimpanzees across Africa. The PCA of the merged dataset show that the first PC separates the Western lineage (*P. t. verus* and *P. t. ellioti*) from the Central/Eastern lineage (*P. t. troglodytes* and *P. t. schweinfurthii*) and the second PC separates *P. t. verus* from *P. t. ellioti* (**Fig. S17a**). There is also clear separation of the two *P. t. ellioti* subpopulations along PC1. The third PC separates *P. t. ellioti* (Ecotone) from the rest of the populations while the fourth PC separates *P. t. schweinfurthii*, indicating that the *P. t. ellioti* sub-populations are potentially more divergent from one another than are the *P. t. troglodytes* and *P. t. schweinfurthii* subspecies (**Fig. S17b & S17c**). There are two captive *P. t. ellioti* individuals that cluster with the *P. t. ellioti* (Ecotone) sub-population, Tobi and Julie_LWC21, which indicate that they originate from the ecotone region in central Cameroon. Additionally, the individual Nicolene which stood out in the PCA of only captive chimpanzees again falls between the *P. t. ellioti* and *P. t. troglodytes* clusters along with the wild individual CMMD06.

The ADMIXTURE analysis of the merged dataset is consistent with the sNMF cluster analysis performed using only captive chimpanzees. At each value of *K*, the same populations are separated from one another (**Fig. S18**). The results from the cross-validation error test for all 100 replicate runs indicate that $K_{MAX} = 4$, and is the best fit for the data (**Fig. S19**). The same wild individuals that show ancestry into different populations from the ADMIXTURE analysis of only wild chimpanzees are also found in this analysis of merged individuals (**Fig. S3**). The captive individual Nicolene has mixed ancestry between *P. t. ellioti* and *P. t. troglodytes*, and Julie_LWC21, Tobi, and Banyo show varying levels of ancestry into the *P. t. ellioti* (Ecotone) subpopulation shows evidence that these captive individuals likely originate from the ecotone region in central Cameroon.

The results from the spatially explicit cluster analysis using TESS recapitulate the population structure recovered from the PCA and ADMIXTURE. At $K = 2$, *P. t. ellioti* separates from *P. t. troglodytes*, and at $K = 3$ *P. t. ellioti* is subdivided into the Rainforest and Ecotone populations (**Fig. S20**). The TESS analysis also identified the same mixed ancestry individuals in the two *P. t. ellioti* populations. Interestingly, the *P. t. ellioti* (Ecotone) population shows a higher level of *P. t. ellioti* (Rainforest) ancestry across almost all individuals than recovered in the PCA and ADMIXTURE analysis (**Fig. S15**). We calculated the *ad hoc* statistic D*K* for all runs *K* = 2-5 and indicated that $K_{MAX}$ = 3, also consistent with the other cluster analyses (**Fig. S21**). The spatial prediction of the *Q* matrix for *K* = 3 show the distribution of the two subspecies across the general range of the Sanaga River and the *P. t. ellioti* subpopulations less clearly defined from on another across the Mbam River, indicating higher levels of admixture between them than between *P. t. ellioti* and *P. t. troglodytes* (**Fig. 1b and S20b**).

We grouped results of the Analysis of Molecular Variance (AMOVA) (**Table S11**) for the '10k' SNP dataset according to three hierarchical sets of variation: (*i*) within sampling locations ($\Phi_{ST}$); (*ii*) among sampling locations in populations ($\Phi_{SC}$); and (*iii*) among the three populations found here and in a previous study [4] – *P. t. ellioti* (Rainforest), *P. t. ellioti* (Ecotone), and *P. t. troglodytes* ($\Phi_{CT}$). The majority of genetic variation (77.86%) found was accounted for variation within sampling locations. Differences between population of origin accounted for the second most variation (16.99%). And the least amount of variation was accounted for among sampling locations within populations (5.15%).

#### Detecting recent hybridization and gene flow in Cameroon

We tested the efficacy of the most informative SNPs using three independently simulated, multigenerational hybrid data sets replicated three times each. The top 100

SNPs allowed correct assignment of an individual as pure *P. t. ellioti*, pure *P. t. troglodytes*, or hybrid 98-100% of the time at all posterior probability thresholds. The mean assignment success for pure *P. t. ellioti* and pure *P. t. troglodytes* was 99% and 100%, respectively, at probability thresholds of 0.99. The mean assignment success for F1 and F2 hybrids was 99% and 87%, respectively, at probability thresholds of 0.99. And finally, the mean assignment success for BC1 and BC2 backcrossed hybrids was 92% and 74%, respectively, at probability thresholds of 0.99, but for BC2 hybrids had a mean assignment success of 100% at a lower probability threshold (0.95). In summary, all genotype classes could be assigned with 99-100% success at probability thresholds of 0.9.

Both *P. t. ellioti* and *P. t. troglodytes* populations consisted of almost entirely of pure individuals, with only one individual of hybrid class in each population. We found strong evidence of an F1 hybrid in the *P. t. ellioti* population (CMMD06) (**Fig. 1b**). This individual had 100% probability of belong to the F1 class in all simulation and replications of the data, and was the same individual identified as a potential hybrid in the PCA (**Figs. S11-14 and S17**), ADMIXTURE analyses (**Figs. S15 & S18**), and TESS analysis (**Fig. S20**). We also found evidence of a backcrossed hybrid in the *P. t. troglodytes* population. CMLB08 was identified as BC1 in *P. t. troglodytes* 33% of the time (with a probability of 0.75). and BC2 66% of the time (with probability of 0.45-0.9). The support for CMLB08 as a likely backcrossed hybrid is not as strong as the support for CMMD06 as an F1 hybrid because (*i*) our statistical power for detecting backcrossed hybrids is not as strong as it is for F1 or F2 hybrids and (*ii*) CMLB08 is only found as a backcrossed hybrid in a small number of runs and with a lower probability.

##### Demographic history model

The asymmetric migration model with the maximum log-likelihood inferred a subspecies split time of approximately 478 kya, with an ancestral effective population size of 24,965 individuals, a *P. t. troglodytes* effective population size of 9,639 individuals and *P. t. ellioti* effective population size 3,030 individuals. The model with the optimized parameters that we chose to capture known aspects of chimpanzee demographic history [1, 2, 4] also showed 0.434 individuals per generation migrating from *P. t. troglodytes to P. t. ellioti*, and 0.517 individuals per generation from *P. t. ellioti* to *P. t. troglodytes*. The residuals of the joint SFS simulated by the model had an approximately normal distribution and were uncorrelated (**Fig. S23**).

### Supplemental Methods

#### Captive chimpanzee genomes

##### Linkage disequilibrium, genetic diversity, and population structure

We assessed genetic diversity within and across chimpanzee subspecies by analyzing a SNP panel, thinned for linkage disequilibrium (LD) using PLINK v1.9 with parameters following previously established methods [1]. The LD-pruned dataset contained 1,113,142 SNPs. Heterozygosity was determined for each sample using vcfhetcount, which is part of the software package vcflib (https://github.com/vcflib/vcflib#vcflib) , calculating median heterozygosity per subspecies.

Population structure was reconstructed through PCA and admixture analysis, utilizing a neutral SNP panel of 147,700 SNPs and an LD-thinned panel of 1,114,534 SNPs. PCA was performed using the package SNPrelate [5] in R v3.4.3 [6]. Admixture analyzed using the 'sNMF' function [7] implemented in the R package LEA v1.2 [8] by estimating an admixture coefficient for each sample that closely approximates commonly used Bayesian clustering programs (e.g., STRUCTURE, ADMIXTURE, etc.) [7, 9, 10]. To identify the best explanation for the results, we evaluated an entropy criterion and used cross-validation to determine the most likely number of ancestral populations (KMAX), comparing it to ∆K [7, 10, 11]. Admixture coefficients were computed for K=2 to K=6 across 1000 iterations and 25 repetitions, choosing the best run by cross-entropy and plotting the results in LEA.

#### Wild chimpanzee population history, and selection analysis

##### Testing for Isolation-by-distance and Isolation-by-environment

We calculated pairwise $F_{ST}$ between sampling locations using Arlequin v3.5 [12] and calculated geographic distance between sampling locations using *geosphere* package in R [13] with n > 1 individuals. We investigated IBD, considering potential confounding by varied habitats [14] because *P. t. ellioti* populations occupy two different niches, one in western forested areas of Cameroon and adjacent parts of Nigeria and another in inhabits the savanna-woodland-rainforest ecotone; *P. t. troglodytes* inhabits the Congolian Rainforest) [4, 15]. We verified this using t-tests for pairwise $F_{ST}$ values and geographic distances. We used a permutation test to discern if geographic distance influenced $F_{ST}$ values across locations more than expected [14]. This test categorized location pairs based on their separating distance and randomized $F_{ST}$ values based on habitat/population. After 10,000 permutations, we derived a null distribution of t-statistics. The empirical *p*-value represented the proportion of null t-statistics exceeding the actual data's t-statistic. This was conducted for both the ‘10k’ and ‘1k’ SNP datasets. In addition to the permutation analysis, we created a binary matrix indicating if sampling location pairs were from the same or different habitat/population. We assessed the correlation between habitat/population classifications and genetic differentiation, adjusting for geographic distance, through a partial Mantel test with 999,999 permutations in R. We used both $F_{ST}$ and ‘linearized $F_{ST}$’ and regular and log-transformed geographic distances. This test was applied to our ‘10k’ and ‘1k’ datasets, specifically to *P. t. ellioti* populations using the ‘10k’ SNP dataset.

#### Population structure of wild chimpanzees in Cameroon

The PCA was performed using the SNPrelate package in R [5], without *a priori* population labels. To confirm the informativeness of the putatively ‘neutral’ SNPs defined using whole genome SNP data from captive chimpanzees (n=2,686 SNPs). Subsequent PCA evaluations were conducted on ‘10k’ and ‘1k’ SNP datasets from wild chimpanzees, and included calculating eigenvalues and the variance of each principal component. The significance of each principal component was assessed using Tracy-Widom statistics, revealing that significant components distinctly grouped samples and sequentially separating populations from most to least differentiated.[16].

ADMIXTURE v1.22 [10] was used to detect population structure and evaluate individual ancestry using our ‘neutral’ SNP dataset (n=2,686 SNPs). ADMIXTURE estimates each individual’s ancestry proportion (Q) into user-defined K clusters. We ran

100 replicates for K=1-10 (1,000 replicates), with a random seed generated for each run. Q matrices were analyzed with DISTRUCT v1.1 [17]. ADMIXTURE's cross-validation error, determined through 5-fold cross-validation for each iteration, helped identify the optimal number of clusters (KMAX) [10]. This was based on the lowest average cross-validation error across 100 iterations for each K. Our analysis also included merging wild chimpanzee samples with 32 captive samples from all four subspecies, ensuring consistent population history and structure. We refined the captive chimpanzee SNP dataset to match the 2,686 neutral SNPs from the wild '10k' dataset. After combining the captive and wild samples into one dataset, we conducted PCA and ADMIXTURE analysis as previously described.

We created spatial interpolations of ancestry proportions (Q matrix) to map the genetic population clusters of wild chimpanzees, using TESS v2.3 [18] and Ad-Mixer v1.0 [19]. Because TESS estimates the shared population history of individuals by incorporating both their genotype and geographic origin, it controls for spatial autocorrelation and improves assessments of population genetic structure, particularly in the presence of IBD [20-22]. We used the ‘1k’ dataset (which includes more sample sites) to maximize geographic sample coverage. We performed 10 replicate runs for K = 2-5 using the CAR model [23], which allows for admixture, with correlated allele frequencies, 10,000 burn-in steps, and 50,000 MCMC iterations per replicate run. We processed TESS outputs with CLUMPP v1.1.2 [24] and DISTRUCT [17]. We determined KMAX by calculating the *ad hoc* statistic ΔK [11] for K = 2-5. We used ASCII posterior predictive maps of admixture proportions generated by TESS as inputs for the program Ad-Mixer to generate spatial interpolations of the Q matrix.

We used Arlequin to perform an AMOVA, testing population differentiation against 10,000 random permutations of the data. We grouped individuals into three populations inferred from ADMIXTURE and PCA anlyses: *P. t. ellioti* (Rainforest), *P. t. ellioti* (Ecotone), and *P. t. troglodytes*. We excluded sampling locations VM and MK due to insufficient sample sizes. We also used Arlequin to calculate pairwise $F_{ST}$ values between *P. t. ellioti* (Rainforest), *P. t. ellioti* (Ecotone), and *P. t. troglodytes*.

#### Genetic diversity in wild chimpanzee populations

We calculated ‘per site’ observed (HO) and expected heterozygosity (HE) using the adegenet package in R [25]. We used per site HO and HE across all individuals to complete a Bartlett test of homogeneity of variances [26] to test the null hypothesis that HO = HE. We also calculated mean HO for all three populations and all sample sites (excluding VM and MK). We calculated individual heterozygosity using VCFtools and plotted it using the ggplot2 package in R [27].

#### Detecting recent hybridization and gene flow

We identified potential hybrids and hybrid classes using the Bayesian clustering program NEWHYBRIDS v1.0 [28] as implemented in the R packages *hybriddetective* [29] and *parallelnewhybrid* [30]. We identified pure individuals from *P. t. troglodytes* and *P. t. ellioti* populations using their membership coefficients (*Q* values) calculated from the ADMIXTURE analysis. Using individuals with Q values >0.80 or <0.10, we created a data set of pure *P. t. ellioti* and *P. t. troglodytes* individuals. The function *hybriddetective* was used on a subset of the 100 most informative SNP loci based on Weir and Cockerham’s $F_{ST}$ to simulate datasets of each of the following six genotype classes: pure parent (for each population), first- or second-generation hybrid (F1 or F2), or F1 backcross with a pure parent (BC1 or BC2). These loci were selected using PLINK. NEWHYBRIDS was then run using *parallelnewhybrid* on three simulated datasets of three replicates each to estimate posterior probabilities of each individual's membership to the six genotype classes. We used *hybriddetective* to determine the efficacy of the program in assigning the correct hybrid class to simulated individuals.

We then filtered our experimental individuals as identified by previously specified *Q* values to our 100 high $F_{ST}$ diagnostic SNP panel using the R package *genepop* [31]. Experimental individual datasets were merged with pure and simulated pure datasets using the function *nh_analysis_generateR*, which allows simulated datasets to be included based on user specified hybrid classes. We then ran the new dataset through NEWHYBRIDS using the Jeffreys prior probabilities and default genotype proportions with the Markov chain directed to run for 300,000 sweeps following a burn-in of 50,000 iterations.

#### Demographic history model

We used *δaδi* (Diffusion Approximations for Demographic Inference) [32] to fit a two-population demographic history for *P. t. ellioti* and *P. t. troglodytes*. *δaδi* approaches demographic history as a diffusion process on the Site Frequency Spectrum (SFS). We performed the analysis using a population informative neutral SNP panel of 2,686 SNPs. We computed the Joint SFS for *P. t. ellioti and P. t. troglodytes* using *δaδi* and projected down from the original allele size to 26 and 78 for *P. t. troglodytes* and *P. t. ellioti*, respectively. We chose this down-projection as it produced the most segregating SNPs through brute force search. Additionally, we used an unfolded SFS using the ancestral state of each variant calculated from the 6-primate EPO alignment [33, 34].

We fit an existing asymmetric migration model to the observed SFS using a modified version of a publicly available optimization framework for *δaδi* [35]. 100 model fits with random starting points were generated without constraints on the parameter space using a Nelder-Mead optimization routine provided by *δaδi* [32, 36]. Model replicates were compared using the negative log-likelihood score and percent missingness in the resulting SFS. We converted *δaδi* parameter estimates to demographic units assuming a constant mutation rate of $1.2 \times 10^{-8}$ per generation per site [2] and a generation time of 25 years [37].

#### Generating environmental data

We obtained environmental data layers from several sources, converted all data layers to 30-arcsecond (~1km$^2$) resolution, transformed them to the WGS84 coordinate projection, and clipped them to chimpanzee-suitable habitats in Cameroon and Nigeria [15, 38]. These layers and their abbreviations are described in **Table S7** and include: (*i*) topographic variables from the NASA Shuttle Radar Topography Mission [39], (*ii*) hydrography of the Sanaga River from HydroSHEDS [40], (*iii*) 19 BioClim variables downloaded from WorldClim [41], (*iv*) Normalized difference vegetation indices [42], surface moisture content from QuikSCAT [43], and (*v*) tree cover from the Global Land Cover Facility [44]. We tested for cross-correlation these layers using pairwise Pearson correlation tests in ENMTools v1.4.1 [45], identifying highly correlated pairs or groups ($-0.85 > p > 0.85$) (**Table S8**).

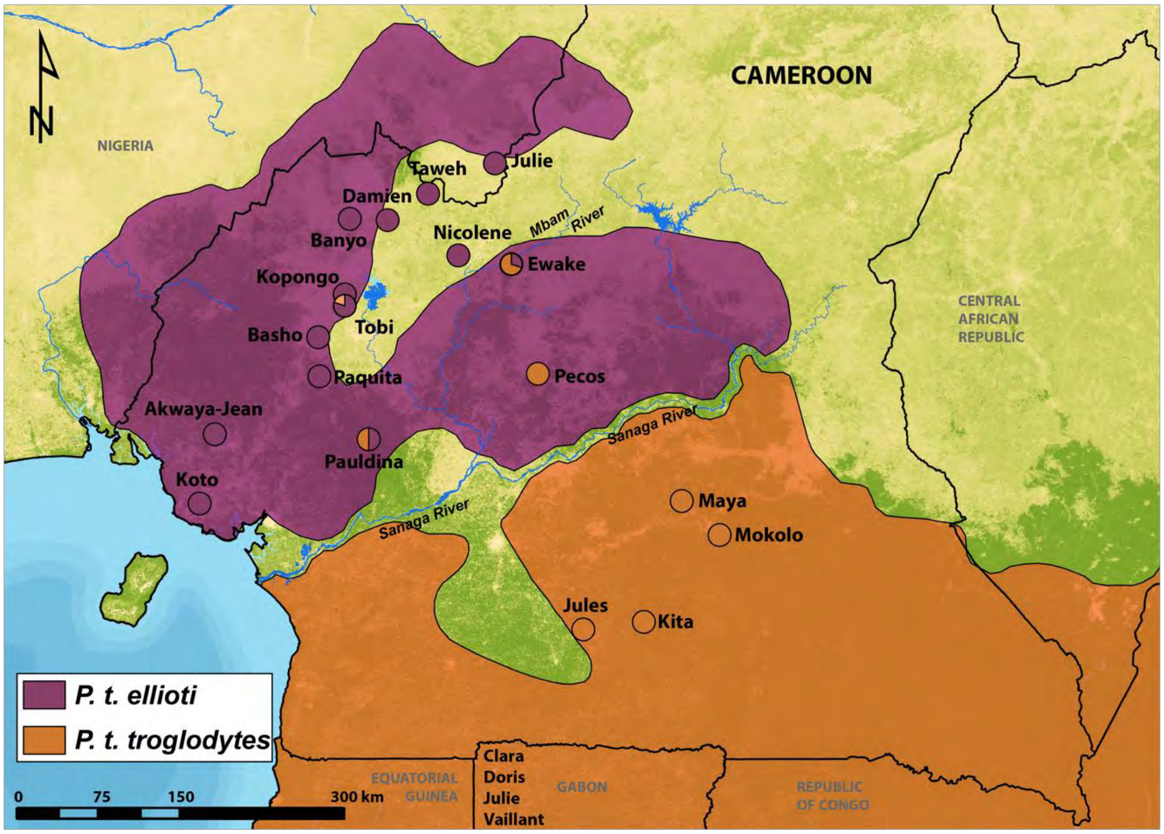


**S1 Fig. Origins of chimpanzees of Cameroon included in this study.**
Sample locations and proportions of estimated ancestry were estimated in previous studies [3, 46].

**Table S1. Captive chimpanzee genomes included in this study.**

| Species | Name | Sample ID | Sex | Source | Geographic origin (Confiscation Point) | Sequencing Center | PE Read Length | Accession number |
|---|---|---|---|---|---|---|---|---|
| *P. t. verus* | Bosco | 9668 10-0004 | M | Liver | | UW | 100 | SRX243448/SRX243449 |
| *P. t. verus* | Clint | C0471 | M | | Captive born | UW | 100 | SRX243527 |
| *P. t. verus* | Jimmie | A956 | F | Blood | Western Africa | CNAG | 100 | SRX243487/SRX243488 |
| *P. t. verus* | Koby | X00100 | M | | | CNAG | 100 | SRX243499 |
| *P. t. ellioti* | Akwaya-Jean | LWC2 | M | Blood | Cameroon (Akwaya, Adamoua) | UW | 100 | SRX243510 |
| *P. t. ellioti* | Banyo | LWC7 | F | Blood | Cameroon (Bankim, Adamoua) | UW | 51 | SRX243511 |
| *P. t. ellioti* | Basho | LWC8 | M | Blood | Cameroon (Takamanda, NW) | UW | 51 | SRX243512 |
| *P. t. ellioti* | Damian | LWC12 | M | Blood | Cameroon (Tinto, NW) | UW | 100 | SRX360476 |
| *P. t. ellioti* | Julie | LWC21 | F | Blood | Cameroon (Douala, Littoral) | UW | 100 | SRX243514 |
| *P. t. ellioti* | Kopongo | LWC23 | F | Blood | Cameroon (Kopongo-Edea Littoral) | UW | 51 | SRX243515 |
| *P. t. ellioti* | Koto | LWC24 | M | Blood | Cameroon (Edea, Littoral) | UW | 100 | SRX243516 |
| *P. t. ellioti* | Paquita | LWC038 | F | Blood | Cameroon (Bertoua, Eastern) | UW | 51 | SRX243517 |
| *P. t. ellioti* | Taweh | LWC43 | M* | Blood | Cameroon (Mamfe, SW) | UW | 100 | SRX243518 |
| *P. t. ellioti* | Tobi | LWC046 | F | Blood | Cameroon (Yaounde, Center) | UW | 51 | SRX243519 |
| *P. t. ellioti* | Nicolene | LWC034 | F | Blood | Cameroon (Tombel, NW) | UW | 101 | SRR28472748 |
| *P. t. troglodytes* | Ewake | LWC015 | F | Blood | Cameroon (Mamfe, NW) | UW | 101 | SRR28472753 |
| *P. t. troglodytes* | Pauldina | LWC039 | F | Blood | Cameroon (Bertoua, East) | UW | 101 | SRR28472747 |
| *P. t. troglodytes* | Pecos | LWC040 | M | Blood | Cameroon (Douala, Littoral) | UW | 101 | SRR28472746 |
| *P. t. troglodytes* | Maya | LWC029 | F | Blood | Cameroon (Yaounde, Center) | UW | 101 | SRR28472750 |
| *P. t. troglodytes* | Kita | LWC022 | F | Blood | Cameroon (Yaounde, Center) | UW | 101 | SRR28472751 |
| *P. t. troglodytes* | Mokolo | LWC031 | M | Blood | Cameroon (Douala, Littoral) | UW | 101 | SRR28472749 |
| *P. t. troglodytes* | Jules | LWC020 | M | Blood | Cameroon (Bertoua, Eastern) | UW | 101 | SRR28472752 |
| *P. t. troglodytes* | Clara | A960 | F | Blood | Gabon | CNAG | 100 | SRX243495/SRX243496 |
| *P. t. troglodytes* | Doris | A958 | F | Blood | Gabon (Ogooue Maritime) | CNAG | 100 | SRX243491/SRX243492 |
| *P. t. troglodytes* | Julie | A959 | F | Blood | Gabon (Haut Ogooue) | CNAG | 100 | SRX243493/SRX243494 |
| *P. t. troglodytes* | Vaillant | A957 | M | Blood | Gabon (Haut Ogooue) | CNAG | 100 | SRX243489/SRX243490 |

| Species | Name | Sample ID | Sex | Source | Geographic origin (Confiscation Point) | Sequencing Center | PE Read Length | Accession number |
|---|---|---|---|---|---|---|---|---|
| *P. t. schweinfurthii* | Andromeda | 100040 | F | Spleen | Tanzania, Gombe National Park | CNAG | 100 | SRX237492 |
| *P. t. schweinfurthii* | Bwambale | A910 | M | Blood | Uganda (Bundibujo District) | CNAG | 100 | SRX237524/SRX237526 |
| *P. t. schweinfurthii* | Harriet | 9729 10-0005 | F | Spleen | Uganda (Bundongo Forest) | UW | 100 | SRX243450/SRX243451 |
| *P. t. schweinfurthii* | Kidongo | A911 | F | Blood | Democratic Republic of Congo | CNAG | 100 | SRX237527/SRX237539 |
| *P. t. schweinfurthii* | Nakuu | A912 | F | Blood | Democratic Republic of Congo | CNAG | 100 | SRX237541/SRX237583 |
| *P. t. schweinfurthii* | Vincent | 100037 | M | Spleen | Tanzania (Gombe National Park) | CNAG | 100 | SRX237455 |

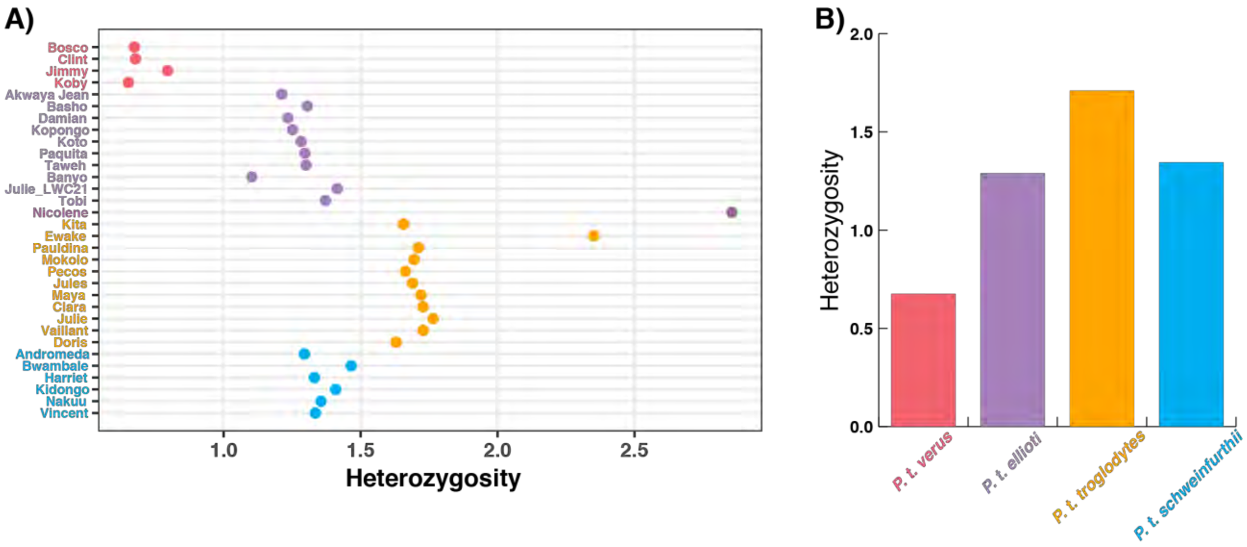


**S2 Fig. Heterozygosity estimates of captive chimpanzee genomes.**
(A) Individual heterozygosity.
(B) Subspecies heterozygosity.

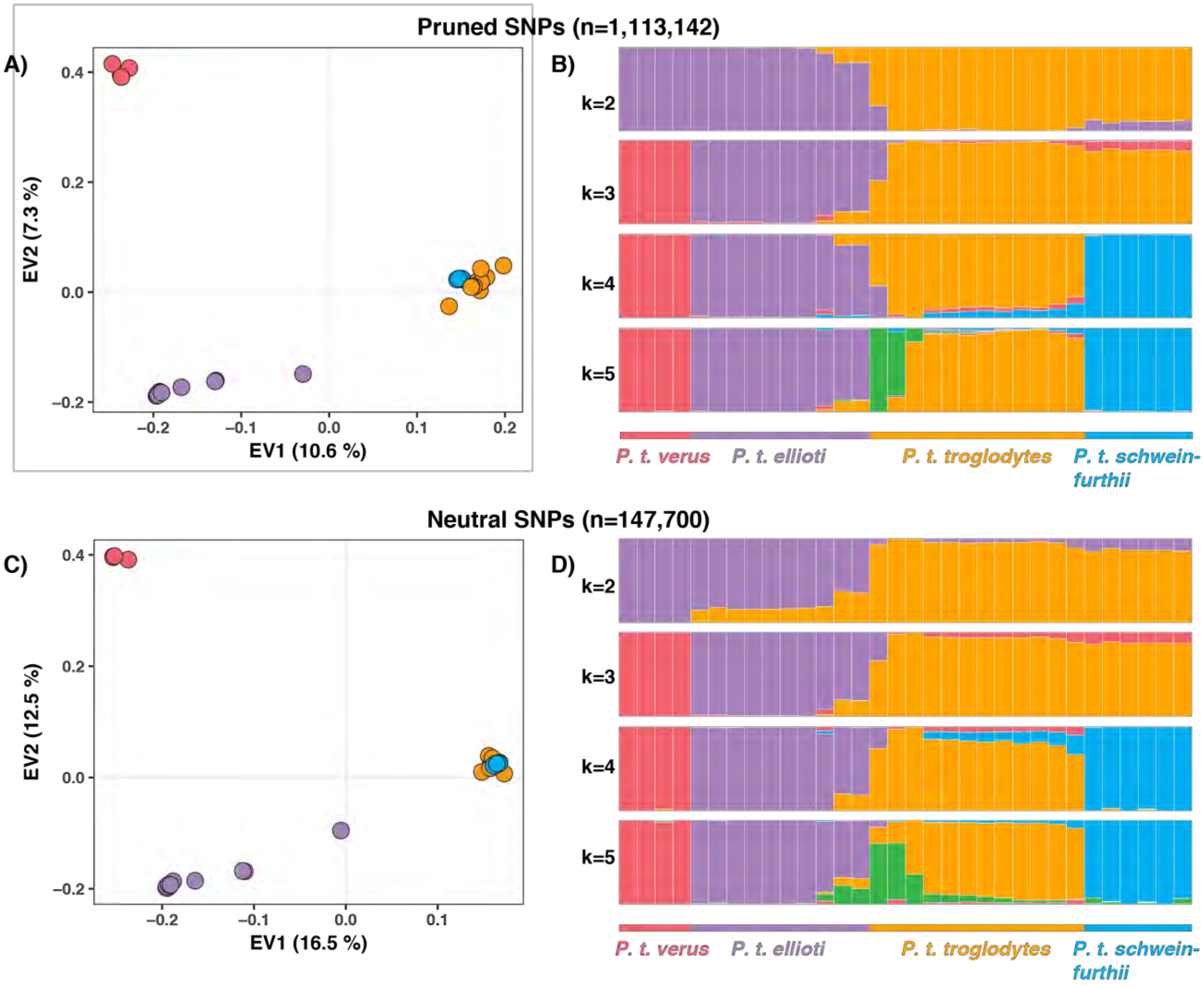


**S3 Fig. Population structure of captive chimpanzee genomes**.
(A) PCA of LD pruned SNP data set consisting of 1,113,142 SNPs.
(B) sNMF individual ancestry analysis of the LD pruned data set in a range of K values.
(C) PCA of Neutral SNP data set consisting of 147,000 SNPs.
(D) sNMF individual ancestry analysis of the neutral SNP data set in a range of K values.

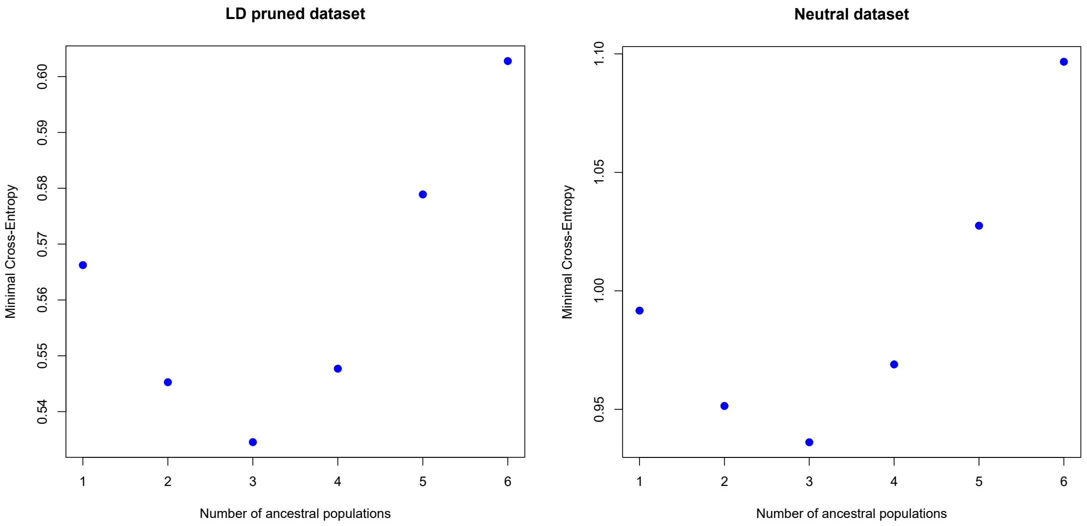


**S4 Fig. Cross Entropy Results.** Value of the cross-entropy criterion as a function of the number of ancestral populations in sNMF for (A) the LD pruned SNP panel and (B) the neutral SNP panel.

**Table S2. Top 10 regions under selection including their genetic content.**

| ID | Region | Content | Name | *p*-value |
|---|---|---|---|---|
| | | | **WESTERN POPULATION** | |
| 1 | 21:8350001:9950000 | ncRNA | snRNA [ENSPTRG00000026954]<br>miRNA [ENSPTRG00000033509]<br>snRNA [ENSPTRG00000036505]<br>misc_RNA [ENSPTRG00000031503] | 7.08165E-05 |
| 2 | 11:46050001:46350000 | ncRNA,<br>protein_coding | miRNA [ENSPTRG00000041850]<br>PHF21A [ENSPTRG00000003542] | 1.34626E-04 |
| 3 | 16:32850001:33450000 | protein_coding | PTHR11254_SF308 [ENSPTRG00000039745] | 1.40292E-04 |
| 4 | 4:164150001:164550000 | protein_coding | FSTL5 [ENSPTRG00000016559] | 1.41633E-04 |
| 5 | 17:82550001:82680442 | protein_coding | B3GNTL1 [ENSPTRG00000009813] | 1.47145E-04 |
| 6 | 3:195250001:195450000 | protein_coding | OSTN [ENSPTRG00000015742]<br>UTS2B [ENSPTRG00000015744]<br>CCDC50 [ENSPTRG00000015745] | 1.98531E-04 |
| 7 | 1:192150001:192350000 | protein_coding | RPS6KC1 [ENSPTRG00000001972] | 2.69251E-04 |
| 8 | 18:67250001:67850000 | intergenic | - | 2.72851E-04 |
| 9 | 1:176450001:176650000 | protein_coding | C1H1orf53 [ENSPTRG00000030241]<br>LHX9 [ENSPTRG00000001814]<br>DENND1B [ENSPTRG00000034501] | 2.80584E-04 |
| 10 | 9:22650001:22950000 | protein_coding | DMRTA1 [ENSPTRG00000020829]<br>MTRNR2L14 [ENSPTRG00000042327] | 2.83266E-04 |
| | | | **CENTRAL/EASTERN POPULATION** | |
| 1 | 1:123550001:123750000 | protein_coding | LRIG2 [ENSPTRG00000001120] | 1.32363E-04 |
| 2 | 16:89250001:89450000 | ncRNA,<br>protein_coding | snRNA [ENSPTRG00000037252]<br>snoRNA [ENSPTRG00000026910]<br>0 [ENSPTRG00000008482]<br>SPATA33 [ENSPTRG00000041260]<br>SPATA2L [ENSPTRG00000008486]<br>VPS9D1 [ENSPTRG00000031104]<br>0 [ENSPTRG00000039136]<br>RPL13 [ENSPTRG00000008480]<br>CDK10 [ENSPTRG00000008485]<br>CHMP1A [ENSPTRG00000008483] | 1.40528E-04 |
| 3 | 8:115650001:116950000 | ncRNA,<br>protein_coding | misc_RNA [ENSPTRG00000039762]<br>snoRNA [ENSPTRG00000037293]<br>snoRNA [ENSPTRG00000033076]<br>SLC30A8 [ENSPTRG00000020523] | 1.43205E-04 |

| | | | | |
|---|---|---|---|---|
| | | | MED30 [ENSPTRG00000020525]<br>0 [ENSPTRG00000040261]<br>EXT1 [ENSPTRG00000020526] | |
| 4 | 13:24350001:24550000 | protein_coding | CENPJ [ENSPTRG00000005715]<br>RNF17 [ENSPTRG00000005713] | 1.45518E-04 |
| 5 | 5:69050001:69750000 | protein_coding | HCN1 [ENSPTRG00000016855] | 1.79372E-04 |
| 6 | 5:62250001:62750000 | protein_coding | MOCS2 [ENSPTRG00000016862]<br>PELO [ENSPTRG00000034409]<br>ITGA2 [ENSPTRG00000016860]<br>ITGA1 [ENSPTRG00000016859] | 1.86150E-04 |
| 7 | 19:47750001:47950000 | protein_coding | PSG8 [ENSPTRG00000039356]<br>PSG3 [ENSPTRG00000039032] | 2.64725E-04 |
| 8 | 2B:235250001:235450000 | ncRNA, protein_coding | ENSPTRG00000027523 [misc_RNA]<br>FBXO36 [ENSPTRG00000013006]<br>SLC16A14 [ENSPTRG00000013007] | 2.81057E-04 |
| 9 | 11:33250001:33450000 | ncRNA, protein_coding | miRNA [ENSPTRG00000035945]<br>HIPK3 [ENSPTRG00000003486] | 2.91036E-04 |
| 10 | 1:222350001:222550000 | protein_coding | CEP170 [ENSPTRG00000002172] | 3.97088E-04 |
| | | | **SHARED** | |
| 1 | 12:15150001:16650000 | ncRNA, protein_coding | MGP [ENSPTRG00000004727]<br>ERP27 [ENSPTRG00000004728]<br>ARHGDIB [ENSPTRG00000004729]<br>PDE6H [ENSPTRG00000004730]<br>RERG [ENSPTRG00000004731]<br>PTPRO [ENSPTRG00000004732]<br>EPS8 [ENSPTRG00000004733]<br>snRNA [ENSPTRG00000025058]<br>misc_RNA [ENSPTRG00000035576]<br>STRAP [ENSPTRG00000004734]<br>DERA [ENSPTRG00000004735]<br>SLC15A5 [ENSPTRG00000004736] | 8.96861E-05 |
| 2 | 17:58750001:59050000 | ncRNA, protein_coding | miRNA [ENSPTRG00000039523]<br>miRNA [ENSPTRG00000041307]<br>miRNA [ENSPTRG00000041661]<br>HEATR6 [ENSPTRG00000009481]<br>CA4 [ENSPTRG00000009482]<br>USP32 [ENSPTRG00000009483]<br>snoRNA [ENSPTRG00000036999] | 2.04082E-04 |
| 3 | 17:37750001:37950000 | ncRNA, protein_coding | 0 [ENSPTRG00000008843]<br>snoRNA [ENSPTRG00000036988]<br>miRNA [ENSPTRG00000041827]<br>DRG2 [ENSPTRG00000008842]<br>GID4 [ENSPTRG00000008841] | 2.94291E-04 |

| | | | | |
|---|---|---|---|---|
| | | | ATPAF2 [ENSPTRG00000008840]<br>LRRC48 [ENSPTRG00000008839]<br>misc_RNA [ENSPTRG00000026228] | |
| 4 | 7:68650001:69950000 | ncRNA, protein_coding | AUTS2 [ENSPTRG00000019260]<br>WBSCR17 [ENSPTRG00000019262]<br>miRNA [ENSPTRG00000041240] | 3.58744E-04 |
| 5 | 1:70250001:70850000 | ncRNA, protein_coding | LRRC7 [ENSPTRG00000000848]<br>misc_RNA [ENSPTRG00000040785]<br>LRRC40 [ENSPTRG00000000850]<br>misc_RNA [ENSPTRG00000039960]<br>SRSF11 [ENSPTRG00000000852] | 7.17489E-04 |
| 6 | 18:66450001:66750000 | protein_coding | SOCS6 [ENSPTRG00000010105] | 1.02041E-03 |
| 7 | 1:69050001:70150000 | protein_coding | RPE65 [ENSPTRG00000000846]<br>DEPDC1 [ENSPTRG00000000847] | 1.03002E-03 |
| 8 | 2A:102550001:102750000 | pseudogene, protein_coding | SNTG2 [ENSPTRG00000011609]<br>IL1RL2 [ENSPTRG00000012300]<br>IL18R1 [ENSPTRG00000012302]<br>pseudogene [ENSPTRG00000000128]<br>IL18RAP [ENSPTRG00000012303] | 1.05890E-03 |
| 9 | 4:191150001:191450000 | protein_coding | ZFP42 [ENSPTRG00000031054]<br>TRIML2 [ENSPTRG00000016668] | 1.12423E-03 |
| 10 | 5:130150001:130550000 | protein_coding | ADAMTS19 [ENSPTRG00000017203] | 1.13306E-03 |

**Table S3. Enriched GO terms in the "Biological Processes" domain.**

| GO ID | Term Description | Genes | *p*-value | Count |
|---|---|---|---|---|
| **WESTERN POPULATION** | | | | |
| GO:0034141 | positive regulation of toll-like receptor 3 signaling pathway | CAV1, F2RL1, PELI1 | 0.015 | 3 of 6 |
| GO:0001942 | hair follicle development | ACVR1B, HOXC13, LDB1, RELA, TNFRSF19 | 0.016 | 5 of 30 |
| GO:0030308 | negative regulation of cell growth | ACVR1B, ACVRL1, ADAM15, BCL2, BMPR2, CCDC85B, FRZB, SERTAD2 | 0.017 | 8 of 82 |
| GO:0006826 | iron ion transport | FTMT, SFXN1 | 0.020 | 3 of 7 |
| GO:0030318 | melanocyte differentiation | BCL2, C10H10orf11, HPS6, OCA2 | 0.026 | 4 of 20 |
| GO:0001701 | in utero embryonic development | ACVR1B, ACVRL1, B9D1, FOSL1, FOXP1, MAPK8IP3, PCGF2, SMAD2, SYF2, UBE2B, WDTC1 | 0.029 | 11 of 154 |
| GO:0051603 | proteolysis involved in cellular protein catabolic process | CTSW, PSMB8, PSMB9 | 0.032 | 5 of 37 |
| GO:0006366 | transcription from RNA polymerase II promoter | BMPR2, FAM170A, HAND1, HIVEP3, LDB1, OVOL1, ZNF536 | 0.040 | 7 of 77 |
| GO:0008104 | protein localization | AKAIN1, CAV1, HAP1, MAPK8IP3 | 0.044 | 5 of 41 |
| GO:0009952 | anterior/posterior pattern specification | ALX1, BMPR2, HOXC10, HOXC11, HOXC13, PCGF2, SMAD2 | 0.046 | 7 of 80 |
| GO:0042445 | hormone metabolic process | CRHBP, LEP, SAFB | 0.048 | 3 of 11 |
| **CENTRAL/EASTERN POPULATION** | | | | |
| GO:0035455 | response to interferon-alpha | BST2, EIF2AK2, IFITM1, IFITM2, IFITM3 | 2.4e-4 | 5 of 11 |
| GO:0035456 | response to interferon-beta | BST2, IFITM1, IFITM2, IFITM3 | 9.1e-4 | 4 of 7 |
| GO:0045071 | negative regulation of viral genome replication | BST2, EIF2AK2, IFITM1, IFITM2, IFITM3, RSAD2 | 0.002 | 6 of 29 |
| GO:0034341 | response to interferon-gamma | BST2, IFITM1, IFITM2, IFITM3 | 0.014 | 4 of 17 |
| GO:0042060 | wound healing | FGF10, INS, TFF1, TFF2, TFF3 | 0.017 | 5 of 33 |
| GO:0042981 | regulation of apoptotic process | BMP1, BMP7, EIF2AK2, IKZF3, MITF, PARK2, RBM25, SARM1, TGFB3, TNFRSF11B | 0.025 | 10 of 137 |
| GO:0030866 | cortical actin cytoskeleton organization | EHD2, LLGL2, ROCK1, ROCK2 | 0.028 | 4 of 22 |
| GO:0019835 | cytolysis | C6, LYZ, MMD | 0.029 | 3 of 9 |
| GO:0007249 | I-kappaB kinase/NF-kappaB signaling | AZI2, ROCK1, ROCK2, TANK | 0.044 | 4 of 26 |
| GO:0006955 | immune response | C6, C7, CCL28, CSF3, CTSS, IL12B, RFX1, TNFAIP1, TNFRSF11B, TNFSF10, VTN | 0.044 | 11 of 176 |
| **SHARED** | | | | |
| GO:0030500 | regulation of bone mineralization | CYP27B1, MGP | 0.044 | 2 of 14 |

**Table S4. Enriched KEGG pathways.**

| KEGG Pathway[a] | Genes | *p*-value | Count |
|---|---|---|---|
| **WESTERN POPULATION** | | | |
| Inflammatory bowel disease (IBD), Asthma, Antigen processing and presentation, Herpes simplex infection, Graft-versus-host disease, Allograft rejection, Type I diabetes mellitus, Viral myocarditis, Leishmaniasis, Toxoplasmosis, Intestinal immune network for IgA production | BCL2, CAV1, CREBBP, HLA-DMA, HLA-DOA, HLA-DQA2, LAMC1, NXF1, PATR-DMB, PATR-DOB, RELA, SMAD2, TAF4, TAF6L, TAP1, TAP2 | 0.078 to 0.041 | 5 to 12 of 31 to 195 |
| Non-alcoholic fatty liver disease (NAFLD) | BAX, CASP7, COX8A, CYP2E1, LEP, MAP3K11, MLXIP, NDUFB7, RELA | 0.024 | 10 of 158 |
| Valine, leucine and isoleucine degradation | ACSF3, ALDH3A2, BCAT2, BCKDHB, ECHS1 | 0.035 | 5 of 47 |
| Axon guidance | CFL1, EFNA1, EFNA3, EFNA4, MET, NFATC3, NTN4, PLXNA4 | 0.05 | 8 of 126 |
| **CENTRAL/EASTERN POPULATION** | | | |
| Dilated cardiomyopathy, Hypertrophic cardiomyopathy (HCM), Arrhythmogenic right ventricular cardiomyopathy (ARVC) | CACNA2D4, CACNG1, CACNG4, ITGA1, ITGA2, ITGB1, PRKACA, SGCA, TGFB3 | 0.029 | 11 of 154 |
| Amoebiasis | COL1A1, COL5A3, IL12B, IL1R2, PIK3R2, PRKACA, PRKCA, RAB5A, TGFB3 | 0.013 | 9 of 107 |

[a]KEGG pathways that share more than half of the significant genes were merged for brevity.

#### Table S5. Functional enrichment clustering.

| Enrichment Score | Term Description | Genes | *p*-value |
|---|---|---|---|
| | **WESTERN POPULATION** | | |
| 2.28 | IPR014745:MHC class II, alpha/beta chain, N-terminal; UP:MHC II; GO:0042613:MHC class II protein complex | PATR-DOB, PATR-DMB, HLA-DMA, HLA-DOA | 0.0047 to 0.0063 |
| 1.68 | ptr05310:Asthma; ptr05332:Graft-versus-host disease; ptr05330:Allograft rejection; ptr04940:Type I diabetes mellitus; ptr04672:Intestinal immune network for IgA production | HLA-DQA2, PATR-DOB, PATR-DMB, HLA-DMA, HLA-DOA | 0.009 to 0.04 |
| 1.58 | IPR001799:Ephrin; IPR019765:Ephrin, conserved site; GO:0046875~ephrin receptor binding | EFNA4, EFNA3, EFNA1 | 0.026 |
| | **CENTRAL/EASTERN POPULATION** | | |
| 1.8 | IPR017994:P-type trefoil, chordata; IPR017957:P-type trefoil, conserved site; SM00018:PD; IPR000519:P-type trefoil | TFF3, TFF2, TFF1 | 0.0029 to 0.0308 |
| 1.68 | IPR025933:Beta-defensin; UP:Defensin; UP:Antibiotic | DEFB125, DEFB126, DEFB127, DEFB129, DEFB132 | 0.0096 to 0.0335 |
| 1.56 | IPR022353:Insulin, conserved site; SM00078:IlGF; IPR016179:Insulin-like | INS, RLN3, INSL6 | 0.0245 to 0.0308 |
| | **SHARED** | | |
| 2.22 | SM00409:IG; IPR003599:Immunoglobulin subtype; IPR007110:Immunoglobulin-like domain | ROBO3, ROBO4, HEPACAM, IL1RL2, IL18RAP, CNTN6 | 0.0028 to 0.0139 |

**Table S6. Number of wild chimpanzee samples collected and used in this study.**

| Sampling Location | Location Code | Latitude | Longitude | Samples sequenced | Samples removed for missingness | Samples removed for relatedness | Samples in '10k' dataset | Samples in '1k' dataset |
|---|---|---|---|---|---|---|---|---|
| Mount Cameroon | MC | 4.28782 | 9.10535 | 3 | 0 | 0 | 3 | 3 |
| Ebo Forest | EB | 4.34543 | 10.41474 | 29 | 8 | 2 | 19 | 23 |
| Liabelem Highlands | ED | 5.399501 | 9.571436 | 3 | 1 | 0 | 2 | 2 |
| Mone | YW | 5.927165 | 9.480388 | 4 | 2 | 0 | 2 | 3 |
| Takamanda | TK | 6.16834 | 9.26445 | 9 | 0 | 6 | 3 | 3 |
| Sabongida | SG | 6.84265 | 10.38777 | 12 | 3 | 5 | 4 | 5 |
| Bankim | BK | 6.123408 | 11.620949 | 2 | 0 | 0 | 2 | 2 |
| Mount Golep | MG | 5.091681 | 11.251401 | 11 | 6 | 0 | 5 | 7 |
| Yagba | YB | 5.097671 | 11.533904 | 13 | 8 | 0 | 5 | 7 |
| Vome | VM | 5.8104 | 12.1967 | 6 | 5 | 0 | 1 | 4 |
| Mbam Djerem | MD | 5.99759 | 12.87796 | 15 | 7 | 1 | 7 | 10 |
| Makombe | MK | 5.3895 | 12.7441 | 3 | 2 | 0 | 1 | 1 |
| Wouchaba | WC | 5.381611 | 13.092472 | 6 | 1 | 0 | 5 | 5 |
| Batanga | KM | 5.191496 | 13.290114 | 9 | 4 | 0 | 5 | 8 |
| Campo Ma'an | CP | 2.34623 | 10.20436 | 6 | 6 | 0 | 0 | 2 |
| Deng Deng | DD | 5.37184 | 13.48651 | 23 | 6 | 13 | 4 | 4 |
| Minta | MT | 4.55219 | 12.912 | 17 | 7 | 1 | 9 | 10 |
| Dja | DJ | 3.31061 | 12.73694 | 7 | 3 | 1 | 3 | 4 |
| Boumba Bek | BB | 2.46819 | 15.23472 | 2 | 2 | 0 | 0 | 0 |
| Lobeke | LB | 2.44953 | 15.40937 | 12 | 2 | 5 | 5 | 5 |
| **Total** | | | | **192** | **73** | **34** | **85** | **108** |

**Table S7. Environmental predictor variables used characterize chimpanzee habitats.**

| Variable Type | Variable Name[a] | Source |
|---|---|---|
| **Topographic Factors** | ***SRTM_HGT – Elevation*** | NASA SRTM; (Farr *et al.* 2007) |
| | ***SRTM_STD – Slope (Ruggedness)*** | |
| | ***Slope_Deg – Slope (Degrees)*** | |
| | ***Slope_Perc – Slope (Percent)*** | |
| | ***Sanaga River*** | HydroSHEDS; (Lehner *et al.* 2008) |
| **Climate Variables** | Bio 1 – Annual Mean Temperature | WorldClim; (Hijmans *et al.* 2005) |
| | Bio 2 – Mean Diurnal Range | |
| | ***Bio 3 – Isothermality*** | |
| | Bio 4 – Temperature Seasonality | |
| | ***Bio 5 – Max. Temp. of the Warmest Month*** | |
| | Bio 6 – Min. Temp. of the Warmest Month | |
| | ***Bio 7 – Temperature Annual Range*** | |
| | Bio 8 – Mean Temp. of the Wettest Quarter | |
| | Bio 9 – Mean Temp. of the Driest Quarter | |
| | Bio 10 – Mean Temp. of the Warmest Quarter | |
| | Bio 11 – Mean Temp. of the Coldest Quarter | |
| | Bio 12 – Annual Precipitation | |
| | Bio 13 – Precipitation of the Wettest Month | |
| | Bio 14 – Precipitation of the Driest Month | |
| | Bio 15 – Precipitation Seasonality | |
| | Bio 16 – Precipitation of the Wettest Quarter | |
| | ***Bio 17 – Precipitation of the Driest Quarter*** | |
| | ***Bio 18 – Precipitation of the Warmest Quarter*** | |
| | ***Bio 19 – Precipitation of the Coldest Quarter*** | |
| **Vegetation Indices** | ***NDVIMAX – Max. Annual NDVI*** | MODIS; (Carroll *et al.* 2004) |
| | ***NDVIMEAN – Mean Annual NDVI*** | |
| | ***NDVIGR – Max. NDVI of Greening Season*** | |
| | ***NDVIBR – Max. NDVI of Least Green Season*** | |
| | NDVIGRBR – NDVI Seasonality | |
| | ***QMEAN – QSCAT (Surface Moisture Content) Annual Mean*** | Quick Scatterometer; (Long *et al.* 2001) |
| | ***QSTD – QSCAT Standard Deviation*** | |
| | ***mdllafrica – MODIS Land Cover*** | MODIS; (Dimiceli *et al.* 2011) |
| | Tree – Percent Tree Cover | |

[a]Data layers used in Gradient Forest analysis are in ***bold italics***.

#### Table S8. Pearson correlation table of environmental variables.

| | bio_01 | bio_02 | bio_03 | bio_04 | bio_05 | bio_06 | bio_07 | bio_08 | bio_09 | bio_10 | bio_11 | bio_12 | bio_13 | bio_14 | bio_15 | bio_16 | bio_17 | bio_18 | bio_19 | mdllafrica | ndvi_brn | ndvi_grn | ndvi_grnbrn | ndvi_max | ndvi_mean | qscat_mean | qscat_std | sanaga | slope_deg | slope_perc | srtm_hgt | srtm_std |
|---|---|---|---|---|---|---|---|---|---|---|---|---|---|---|---|---|---|---|---|---|---|---|---|---|---|---|---|---|---|---|---|---|
| bio_01 | 0 | 0 | 0 | 0 | 0 | 0 | 0 | 0 | 0 | 0 | 0 | 0 | 0 | 0 | 0 | 0 | 0 | 0 | 0 | 0 | 0 | 0 | 0 | 0 | 0 | 0 | 0 | 0 | 0 | 0 | 0 | 0 |
| bio_02 | -0.297 | 0 | 0 | 0 | 0 | 0 | 0 | 0 | 0 | 0 | 0 | 0 | 0 | 0 | 0 | 0 | 0 | 0 | 0 | 0 | 0 | 0 | 0 | 0 | 0 | 0 | 0 | 0 | 0 | 0 | 0 | 0 |
| bio_03 | -0.314 | -0.091 | 0 | 0 | 0 | 0 | 0 | 0 | 0 | 0 | 0 | 0 | 0 | 0 | 0 | 0 | 0 | 0 | 0 | 0 | 0 | 0 | 0 | 0 | 0 | 0 | 0 | 0 | 0 | 0 | 0 | 0 |
| bio_04 | 0.280 | -0.102 | -0.890 | 0 | 0 | 0 | 0 | 0 | 0 | 0 | 0 | 0 | 0 | 0 | 0 | 0 | 0 | 0 | 0 | 0 | 0 | 0 | 0 | 0 | 0 | 0 | 0 | 0 | 0 | 0 | 0 | 0 |
| bio_05 | 0.831 | 0.193 | -0.596 | 0.497 | 0 | 0 | 0 | 0 | 0 | 0 | 0 | 0 | 0 | 0 | 0 | 0 | 0 | 0 | 0 | 0 | 0 | 0 | 0 | 0 | 0 | 0 | 0 | 0 | 0 | 0 | 0 | 0 |
| bio_06 | 0.840 | -0.701 | 0.019 | 0.106 | 0.449 | 0 | 0 | 0 | 0 | 0 | 0 | 0 | 0 | 0 | 0 | 0 | 0 | 0 | 0 | 0 | 0 | 0 | 0 | 0 | 0 | 0 | 0 | 0 | 0 | 0 | 0 | 0 |
| bio_07 | -0.089 | 0.874 | -0.557 | 0.342 | 0.453 | -0.593 | 0 | 0 | 0 | 0 | 0 | 0 | 0 | 0 | 0 | 0 | 0 | 0 | 0 | 0 | 0 | 0 | 0 | 0 | 0 | 0 | 0 | 0 | 0 | 0 | 0 | 0 |
| bio_08 | 0.956 | -0.244 | -0.106 | 0.034 | 0.737 | 0.809 | -0.143 | 0 | 0 | 0 | 0 | 0 | 0 | 0 | 0 | 0 | 0 | 0 | 0 | 0 | 0 | 0 | 0 | 0 | 0 | 0 | 0 | 0 | 0 | 0 | 0 | 0 |
| bio_09 | 0.968 | -0.436 | -0.352 | 0.401 | 0.774 | 0.890 | -0.191 | 0.885 | 0 | 0 | 0 | 0 | 0 | 0 | 0 | 0 | 0 | 0 | 0 | 0 | 0 | 0 | 0 | 0 | 0 | 0 | 0 | 0 | 0 | 0 | 0 | 0 |
| bio_10 | 0.987 | -0.234 | -0.444 | 0.408 | 0.891 | 0.778 | 0.027 | 0.911 | 0.961 | 0 | 0 | 0 | 0 | 0 | 0 | 0 | 0 | 0 | 0 | 0 | 0 | 0 | 0 | 0 | 0 | 0 | 0 | 0 | 0 | 0 | 0 | 0 |
| bio_11 | 0.982 | -0.222 | -0.199 | 0.119 | 0.805 | 0.807 | -0.079 | 0.983 | 0.913 | 0.954 | 0 | 0 | 0 | 0 | 0 | 0 | 0 | 0 | 0 | 0 | 0 | 0 | 0 | 0 | 0 | 0 | 0 | 0 | 0 | 0 | 0 | 0 |
| bio_12 | 0.313 | -0.672 | -0.208 | 0.303 | 0.053 | 0.515 | -0.466 | 0.165 | 0.395 | 0.304 | 0.222 | 0 | 0 | 0 | 0 | 0 | 0 | 0 | 0 | 0 | 0 | 0 | 0 | 0 | 0 | 0 | 0 | 0 | 0 | 0 | 0 | 0 |
| bio_13 | 0.256 | -0.722 | -0.305 | 0.439 | 0.025 | 0.480 | -0.456 | 0.082 | 0.378 | 0.265 | 0.139 | 0.940 | 0 | 0 | 0 | 0 | 0 | 0 | 0 | 0 | 0 | 0 | 0 | 0 | 0 | 0 | 0 | 0 | 0 | 0 | 0 | 0 |
| bio_14 | 0.014 | -0.395 | 0.564 | -0.588 | -0.400 | 0.235 | -0.596 | 0.187 | -0.035 | -0.105 | 0.086 | 0.229 | 0.091 | 0 | 0 | 0 | 0 | 0 | 0 | 0 | 0 | 0 | 0 | 0 | 0 | 0 | 0 | 0 | 0 | 0 | 0 | 0 |
| bio_15 | 0.107 | 0.103 | -0.761 | 0.809 | 0.429 | -0.077 | 0.463 | -0.112 | 0.194 | 0.236 | -0.012 | 0.095 | 0.298 | -0.818 | 0 | 0 | 0 | 0 | 0 | 0 | 0 | 0 | 0 | 0 | 0 | 0 | 0 | 0 | 0 | 0 | 0 | 0 |
| bio_16 | 0.284 | -0.559 | -0.421 | 0.530 | 0.159 | 0.414 | -0.269 | 0.078 | 0.390 | 0.316 | 0.161 | 0.946 | 0.959 | -0.047 | 0.384 | 0 | 0 | 0 | 0 | 0 | 0 | 0 | 0 | 0 | 0 | 0 | 0 | 0 | 0 | 0 | 0 | 0 |
| bio_17 | 0.006 | -0.423 | 0.585 | -0.599 | -0.423 | 0.249 | -0.630 | 0.168 | -0.036 | -0.115 | 0.076 | 0.283 | 0.125 | 0.977 | -0.831 | -0.002 | 0 | 0 | 0 | 0 | 0 | 0 | 0 | 0 | 0 | 0 | 0 | 0 | 0 | 0 | 0 | 0 |
| bio_18 | -0.125 | -0.206 | 0.616 | -0.663 | -0.431 | 0.073 | -0.462 | 0.043 | -0.217 | -0.231 | -0.031 | 0.198 | 0.051 | 0.761 | -0.730 | -0.045 | 0.790 | 0 | 0 | 0 | 0 | 0 | 0 | 0 | 0 | 0 | 0 | 0 | 0 | 0 | 0 | 0 |
| bio_19 | 0.270 | -0.307 | -0.523 | 0.594 | 0.300 | 0.287 | -0.015 | 0.035 | 0.353 | 0.332 | 0.152 | 0.825 | 0.806 | -0.276 | 0.477 | 0.923 | -0.225 | -0.207 | 0 | 0 | 0 | 0 | 0 | 0 | 0 | 0 | 0 | 0 | 0 | 0 | 0 | 0 |
| mdllafrica | 0.033 | -0.151 | 0.412 | -0.434 | -0.189 | 0.159 | -0.329 | 0.152 | -0.013 | -0.046 | 0.094 | 0.004 | -0.070 | 0.409 | -0.481 | -0.136 | 0.420 | 0.389 | -0.222 | 0 | 0 | 0 | 0 | 0 | 0 | 0 | 0 | 0 | 0 | 0 | 0 | 0 |
| ndvi_brn | 0.028 | 0.484 | -0.109 | -0.023 | 0.242 | -0.228 | 0.446 | 0.055 | -0.041 | 0.051 | 0.064 | -0.247 | -0.360 | -0.029 | -0.128 | -0.238 | -0.029 | -0.068 | -0.080 | -0.055 | 0 | 0 | 0 | 0 | 0 | 0 | 0 | 0 | 0 | 0 | 0 | 0 |
| ndvi_grn | 0.063 | 0.310 | 0.031 | -0.111 | 0.177 | -0.067 | 0.226 | 0.092 | 0.001 | 0.064 | 0.102 | -0.207 | -0.281 | -0.139 | -0.101 | -0.199 | -0.110 | -0.060 | -0.070 | 0.227 | 0.302 | 0 | 0 | 0 | 0 | 0 | 0 | 0 | 0 | 0 | 0 | 0 |
| ndvi_grn_brn | 0.001 | -0.354 | 0.128 | -0.029 | -0.167 | 0.204 | -0.354 | -0.014 | 0.043 | -0.023 | -0.018 | 0.157 | 0.239 | -0.036 | 0.084 | 0.152 | -0.022 | 0.042 | 0.050 | 0.164 | -0.892 | 0.162 | 0 | 0 | 0 | 0 | 0 | 0 | 0 | 0 | 0 | 0 |
| ndvi_max | 0.171 | -0.076 | 0.099 | -0.130 | 0.068 | 0.173 | -0.112 | 0.217 | 0.142 | 0.138 | 0.194 | 0.056 | -0.003 | 0.187 | -0.195 | -0.015 | 0.194 | 0.153 | -0.049 | 0.270 | 0.134 | 0.385 | 0.044 | 0 | 0 | 0 | 0 | 0 | 0 | 0 | 0 | 0 |
| ndvi_mean | 0.066 | 0.494 | -0.006 | -0.114 | 0.261 | -0.161 | 0.396 | 0.110 | -0.014 | 0.077 | 0.119 | -0.324 | -0.436 | -0.131 | -0.161 | -0.313 | -0.109 | -0.096 | -0.132 | 0.155 | 0.736 | 0.741 | -0.410 | 0.335 | 0 | 0 | 0 | 0 | 0 | 0 | 0 | 0 |
| qscat_mean | 0.134 | -0.406 | 0.511 | -0.491 | -0.246 | 0.361 | -0.582 | 0.255 | 0.088 | 0.029 | 0.192 | 0.261 | 0.151 | 0.590 | -0.592 | 0.052 | 0.617 | 0.584 | -0.097 | 0.568 | -0.238 | 0.079 | 0.283 | 0.270 | -0.059 | 0 | 0 | 0 | 0 | 0 | 0 | 0 |
| qscat_std | -0.147 | 0.420 | -0.560 | 0.532 | 0.267 | -0.376 | 0.615 | -0.307 | -0.110 | -0.030 | -0.215 | -0.115 | -0.027 | -0.661 | 0.660 | 0.104 | -0.679 | -0.582 | 0.300 | -0.582 | 0.183 | -0.084 | -0.229 | -0.292 | 0.022 | -0.826 | 0 | 0 | 0 | 0 | 0 | 0 |
| sanaga | 0.169 | 0.107 | -0.644 | 0.717 | 0.467 | 0.029 | 0.392 | -0.063 | 0.245 | 0.282 | 0.056 | 0.357 | 0.395 | -0.671 | 0.754 | 0.572 | -0.637 | -0.516 | 0.737 | -0.409 | 0.052 | 0.024 | -0.043 | -0.140 | 0.020 | -0.440 | 0.592 | 0 | 0 | 0 | 0 | 0 |
| slope_deg | -0.262 | 0.131 | -0.320 | 0.323 | -0.073 | -0.332 | 0.265 | -0.369 | -0.231 | -0.193 | -0.319 | 0.177 | 0.207 | -0.209 | 0.298 | 0.269 | -0.197 | -0.148 | 0.326 | -0.157 | 0.043 | -0.044 | -0.066 | -0.066 | -0.040 | -0.140 | 0.238 | 0.330 | 0 | 0 | 0 | 0 |
| slope_perc | -0.262 | 0.131 | -0.318 | 0.321 | -0.074 | -0.331 | 0.264 | -0.369 | -0.232 | -0.193 | -0.319 | 0.176 | 0.206 | -0.208 | 0.297 | 0.268 | -0.197 | -0.147 | 0.324 | -0.157 | 0.042 | -0.045 | -0.065 | -0.067 | -0.041 | -0.139 | 0.237 | 0.329 | 1.000 | 0 | 0 | 0 |
| srtm_hgt | -0.899 | 0.464 | 0.099 | -0.061 | -0.582 | -0.858 | 0.332 | -0.905 | -0.856 | -0.847 | -0.903 | -0.439 | -0.345 | -0.344 | 0.168 | -0.318 | -0.341 | -0.166 | -0.218 | -0.184 | 0.047 | -0.001 | -0.050 | -0.220 | 0.031 | -0.383 | 0.388 | 0.051 | 0.288 | 0.287 | 0 | 0 |
| srtm_std | -0.257 | 0.125 | -0.274 | 0.278 | -0.089 | -0.318 | 0.238 | -0.347 | -0.229 | -0.196 | -0.308 | 0.159 | 0.189 | -0.172 | 0.257 | 0.239 | -0.159 | -0.106 | 0.278 | -0.115 | 0.022 | -0.032 | -0.038 | -0.035 | -0.044 | -0.089 | 0.177 | 0.273 | 0.714 | 0.712 | 0.311 | 0 |

**Table S9. Environmental variable groupings[a].**

| **General Temperature** | **Temperature Range** | **Temperature Seasonality** |
|---|---|---|
| Bio_1 | Bio_2 | Bio_3 |
| Bio_5 | Bio_7 | Bio_4 |
| Bio_6 | **Precipitation (Wet/Cold)** | **Precipitation (Dry/Warm)** |
| Bio_8 | | |
| Bio_9 | Bio_12 | Bio_14 |
| Bio_10 | Bio_13 | Bio_15 |
| Bio_11 | Bio_16 | Bio_17 |
| **Topography** | Bio_19 | Bio_18 |
| SRTM_HGT | **Surface Moisture** | **Tree Cover** |
| SRTM_STD | QMEAN | mdllafrica |
| Slope_Deg | QSTD | Tree |
| Slope_Perc | **Vegetation Greenness** | |
| **Vegetation Brownness** | NDVIGR | |
| NDVIBR | NDVIMAX | |
| NDVIGRBR | NDVIMEAN | |

[a]Based on Pearson correlation values.

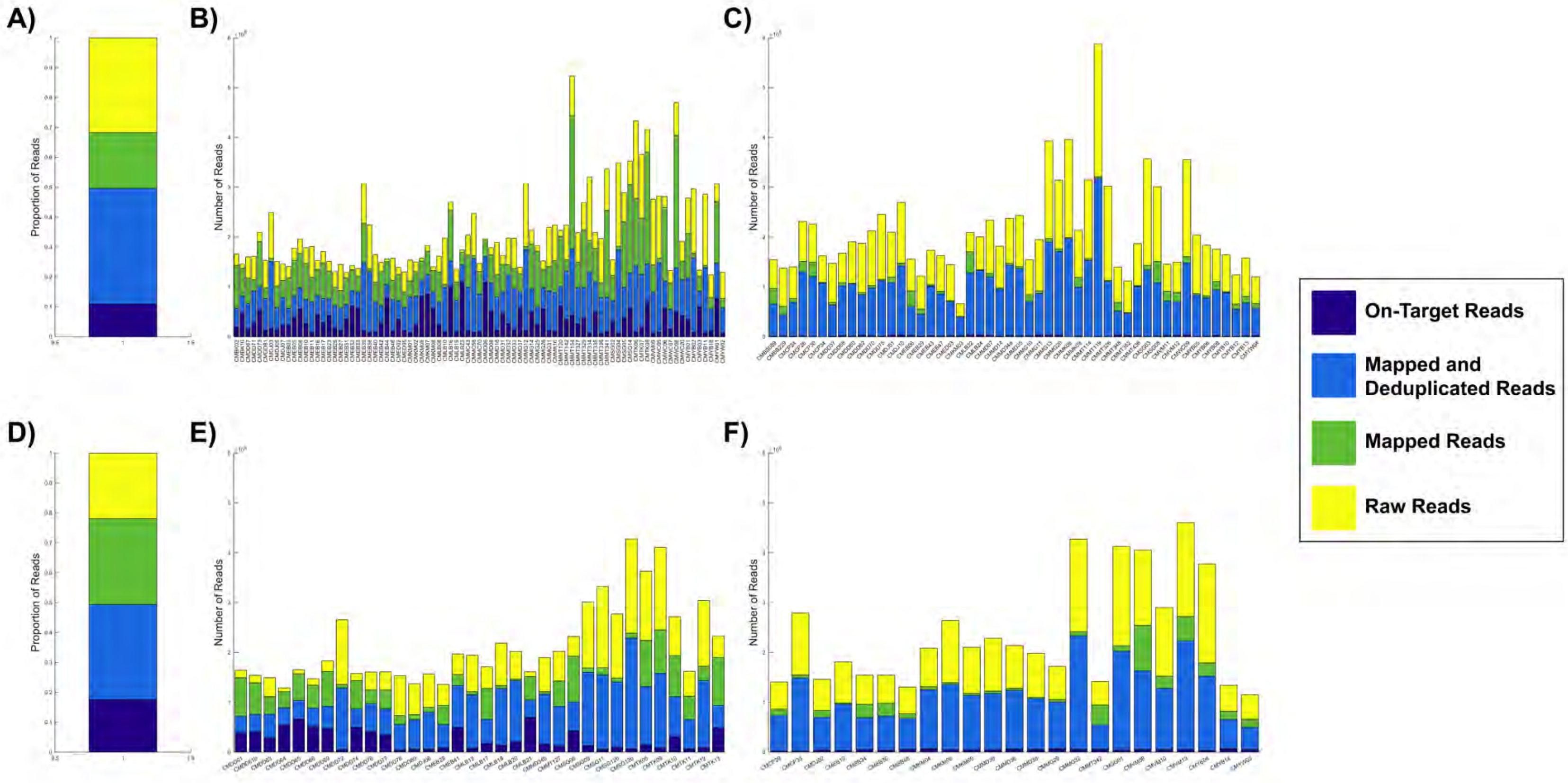


**S5 Fig. Illumina reads of wild chimpanzee samples evaluated into 4 categories.**
Raw reads – yellow, mapped reads (sequence reads that mapped to the panTro4 reference genome) – green, mapped and deduplicated reads (PCR duplicates were removed) – light blue, and on-target reads mapped to our sites – dark blue.
(A) Proportion of read types for all 192 sequenced chimpanzee samples.
(B) Number of read types for 85 samples included in the ‘10k’ dataset.
(C) Number of read types for samples removed due to missingness (>30% missing sites).
(D) Proportion of read types for 85 samples included in the ‘complete’ dataset.
(E) Number of read types for duplicate samples removed following relatedness analysis.
(F) Number of read types for 23 individuals included in the ‘1k’ dataset, but not the ‘10k’ dataset.

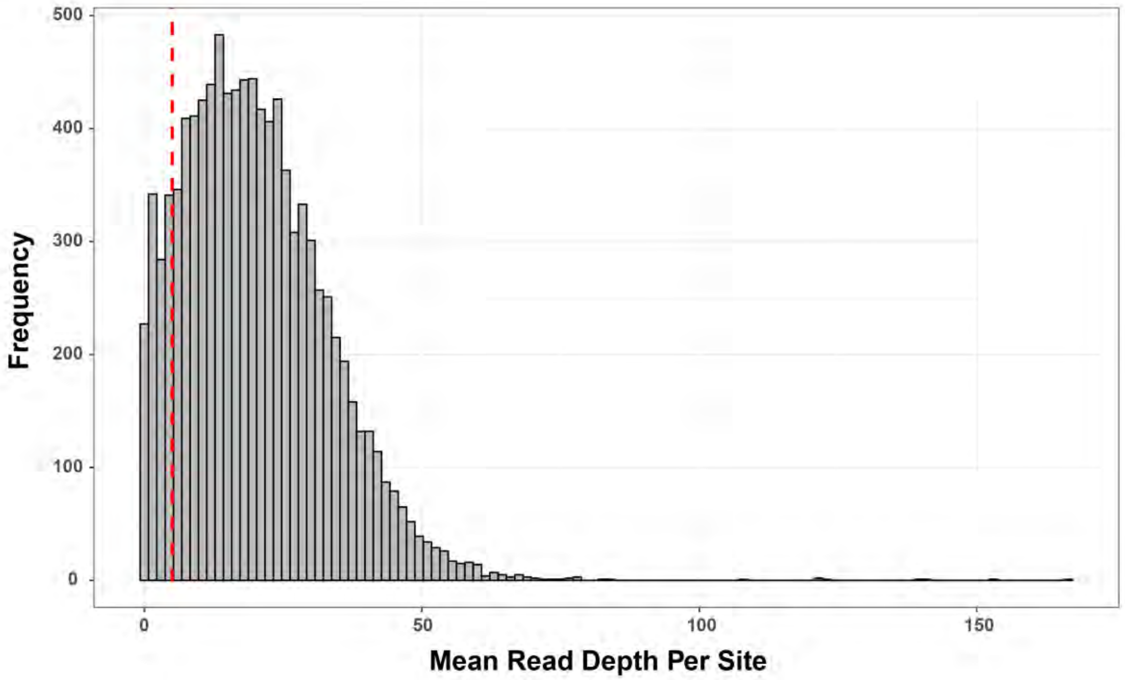


**S6 Fig. Frequency of mean sequencing coverage of wild chimpanzee samples for each site.**
Mean coverage across all sites was 20.2 reads/site. The red vertical line represents the minimum coverage needed to accurately call SNPs (5x coverage).

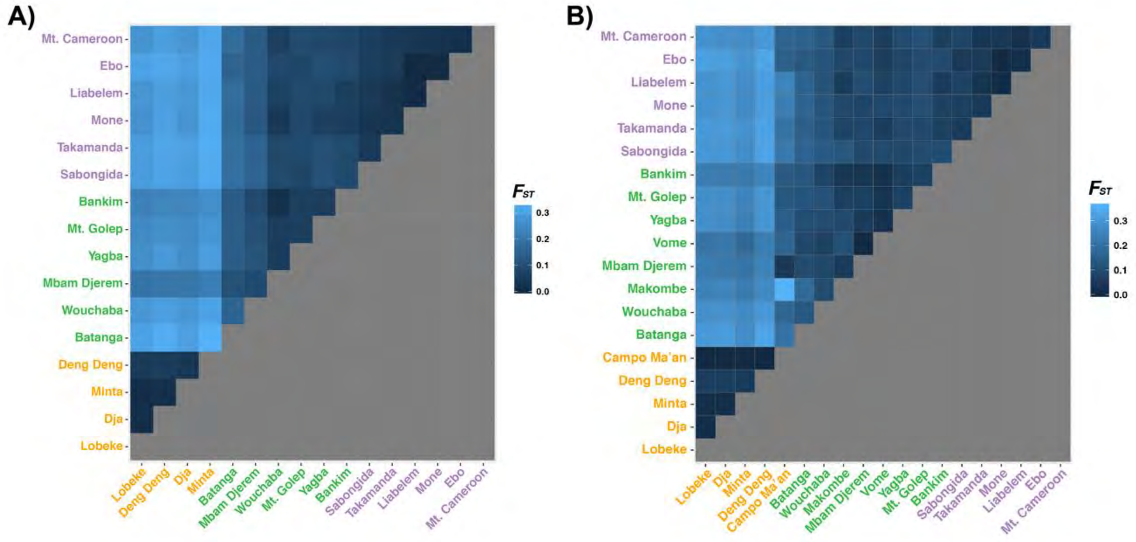


**S7 Fig. Pairwise $F_{ST}$ between sites shows population structure. (A) '10k' dataset. (B) '1k' dataset.**

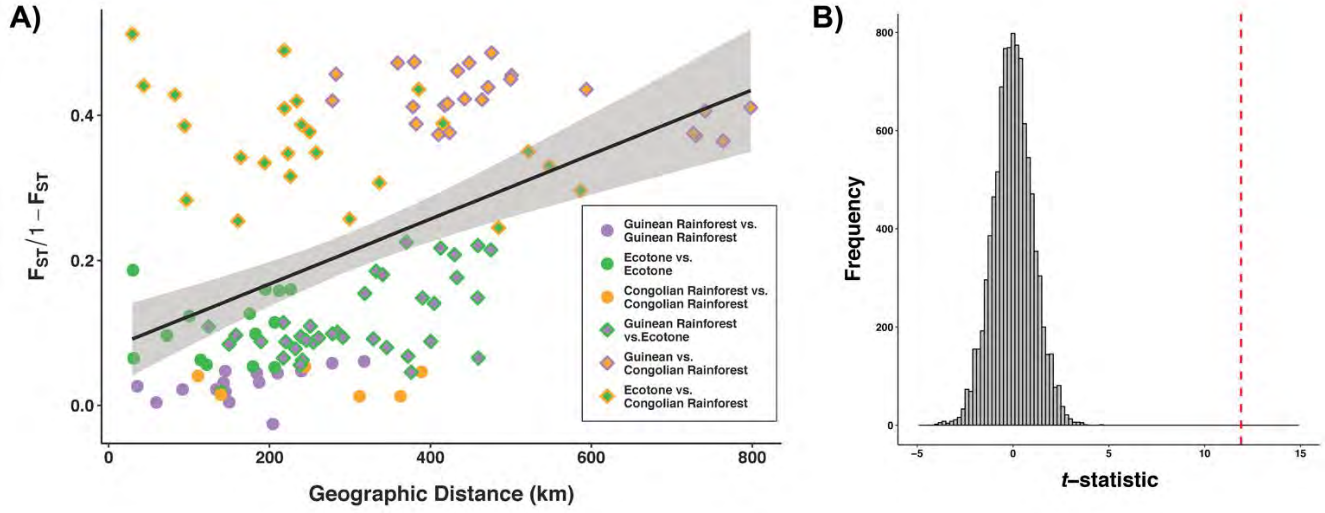


**S8 Fig. Isolation-by-distance for '10k' dataset.**

(A) Correlation between 'linearized $F_{ST}$' and geographic distance (km) generated using the '10k' dataset. Solid circles represent pairs of sampling locations from the same habitat. Dual-colored diamonds represent pairs of sampling locations from different habitats.

(B) Null distribution of t-statistics from 10,000 permutations same- or different habitat/population pairs in four bins of geographic distance. Red dotted line shows the t-statistic value for actual data.

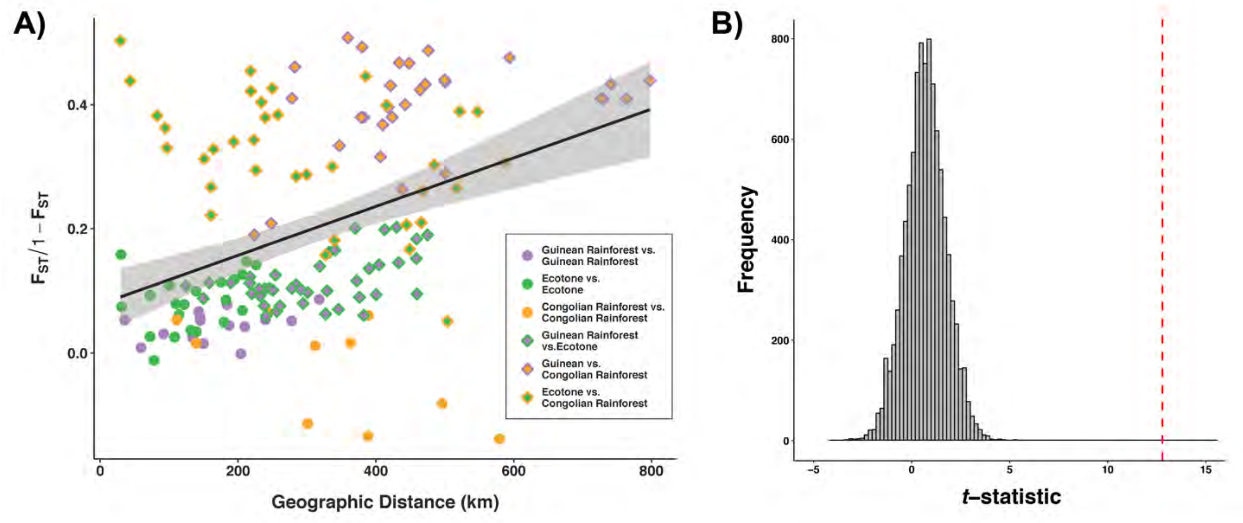


**S9 Fig. Isolation-by-distance for ‘1k’ dataset.**

(A) Correlation between ‘linearized $F_{ST}$’ and geographic distance (km) generated using the ‘1k’ dataset. Solid circles represent pairs of sampling locations from the same habitat. Dual-colored diamonds represent pairs of sampling locations from different habitats.

(B) Null distribution of *t*-statistics from 10,000 permutations same- or different habitat/population pairs in four bins of geographic distance. Red dotted line shows the *t*-statistic value for actual data.

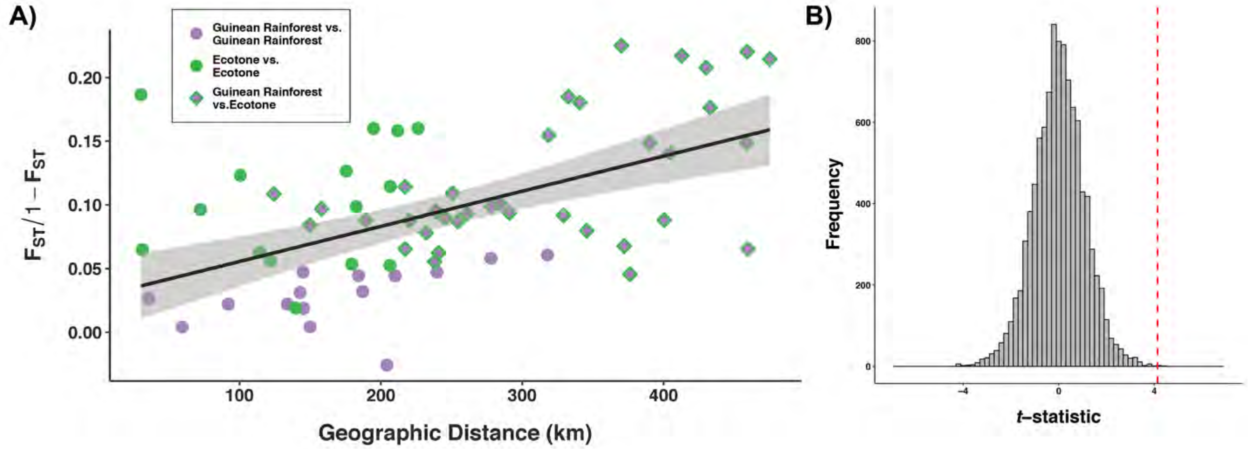


**S10 Fig. Isolation-by-distance for only *P. t. ellioti* populations.**
(A) Correlation between 'linearized $F_{ST}$' and geographic distance (km) generated using the '10k' dataset. Solid circles represent pairs of sampling locations from the same habitat. Dual-colored diamonds represent pairs of sampling locations from different habitats.
(B) Null distribution of *t*-statistics from 10,000 permutations same- or different habitat/population pairs in four bins of geographic distance. Red dotted line shows the *t*-statistic value for actual data.

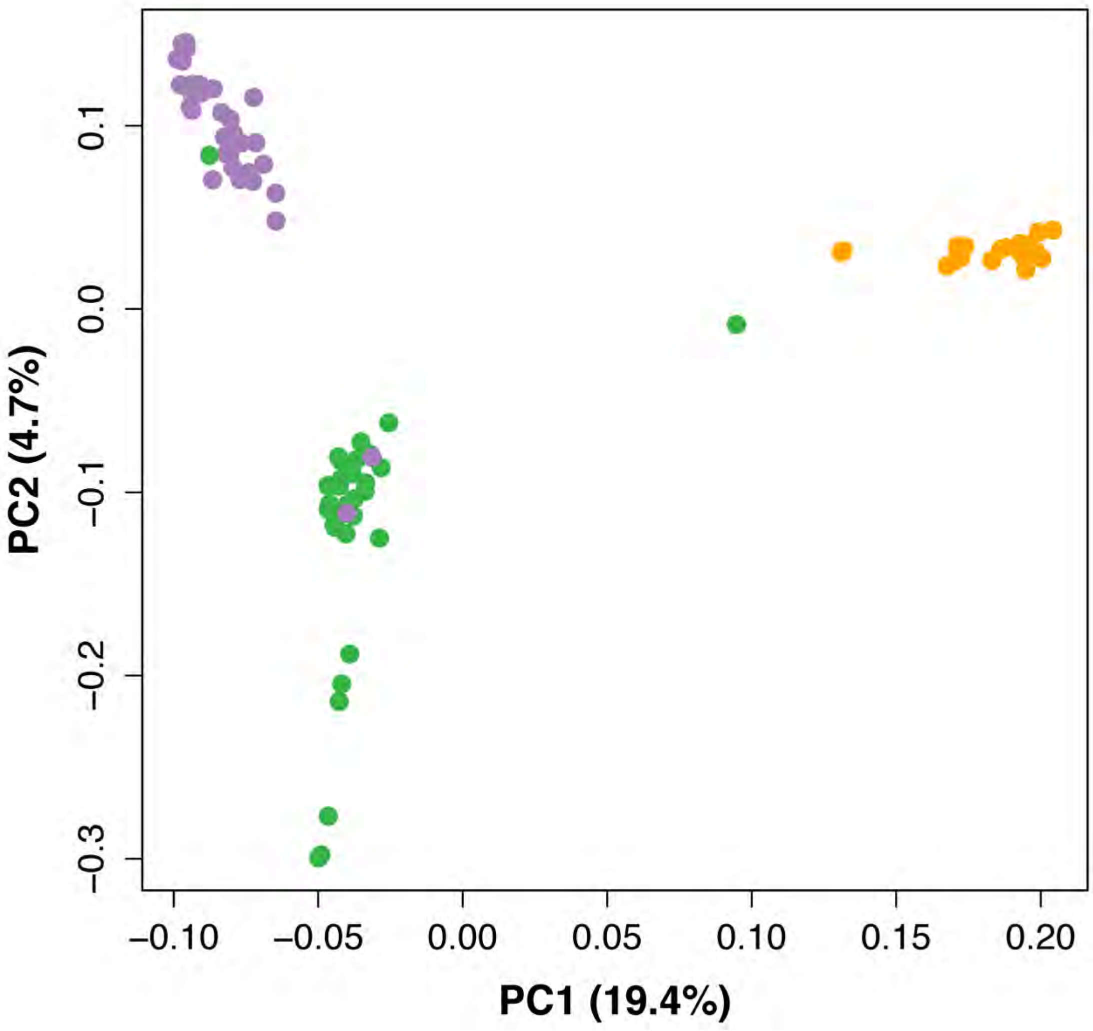


**S11 Fig. PCA of all SNPs from the '10k' dataset.**
The first two principal components recapitulate known population structure of chimpanzees in Cameroon. They show 3 clear populations, and one *P. t. ellioti* (Ecotone) individual (CMMD06) clustering with *P. t. troglodytes*, as well as multiple *P. t. ellioti* (Rainforest) clustering together with *P. t. ellioti* (Ecotone) and vice versa.

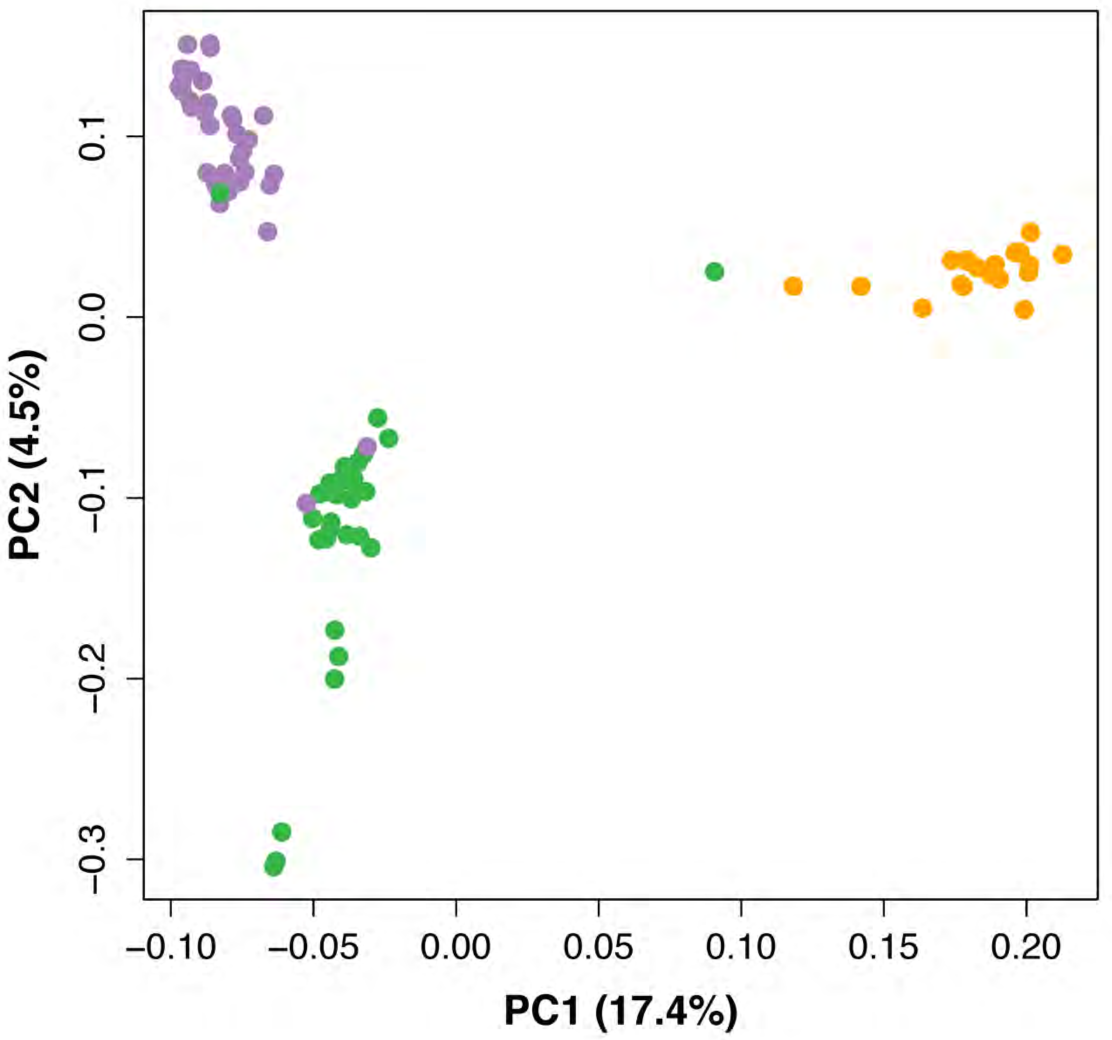


**S12 Fig. PCA of only neutral SNPs from the '10k' dataset.**

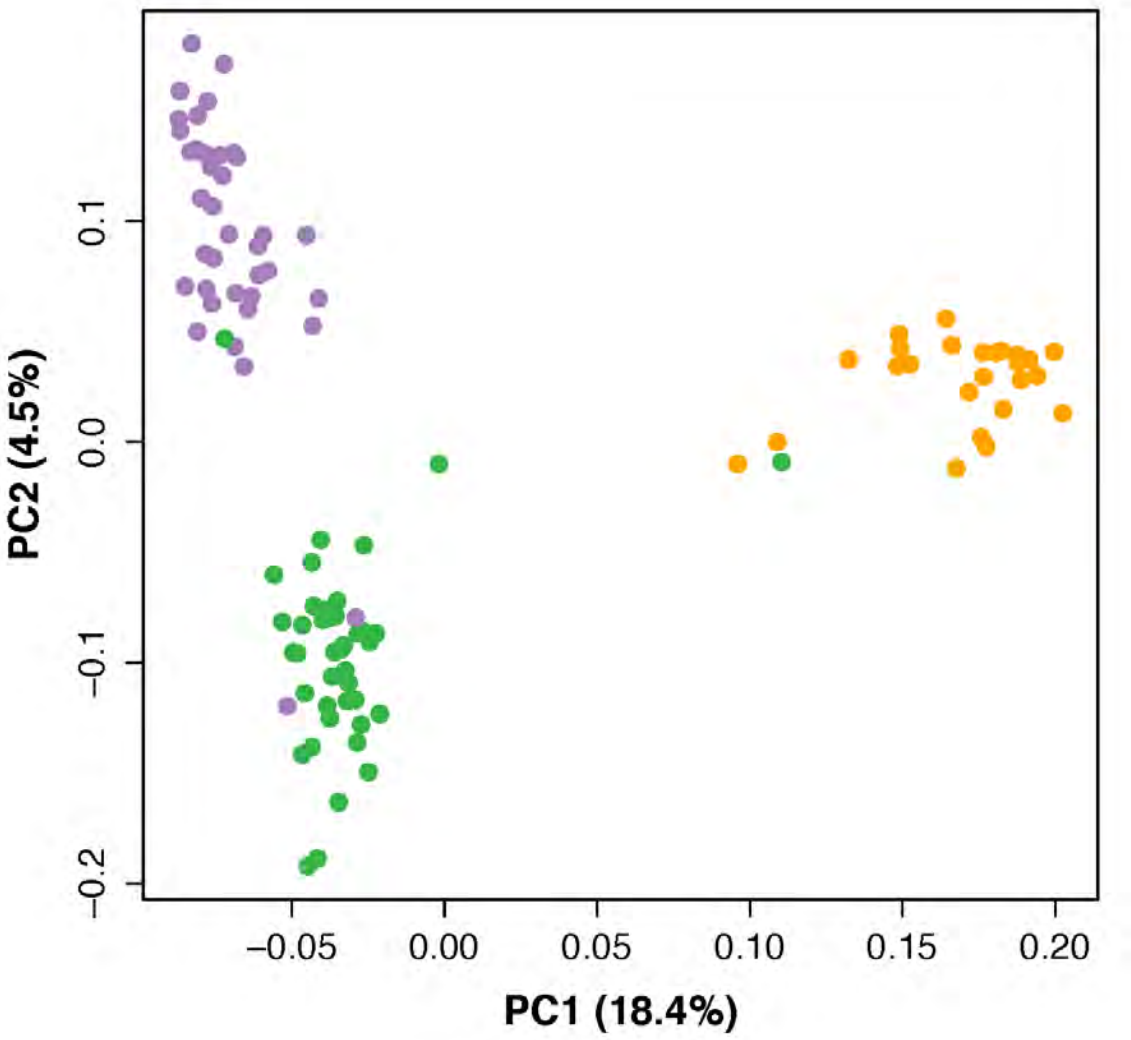


**S13 Fig. PCA of all SNPs from the '1k' dataset.**

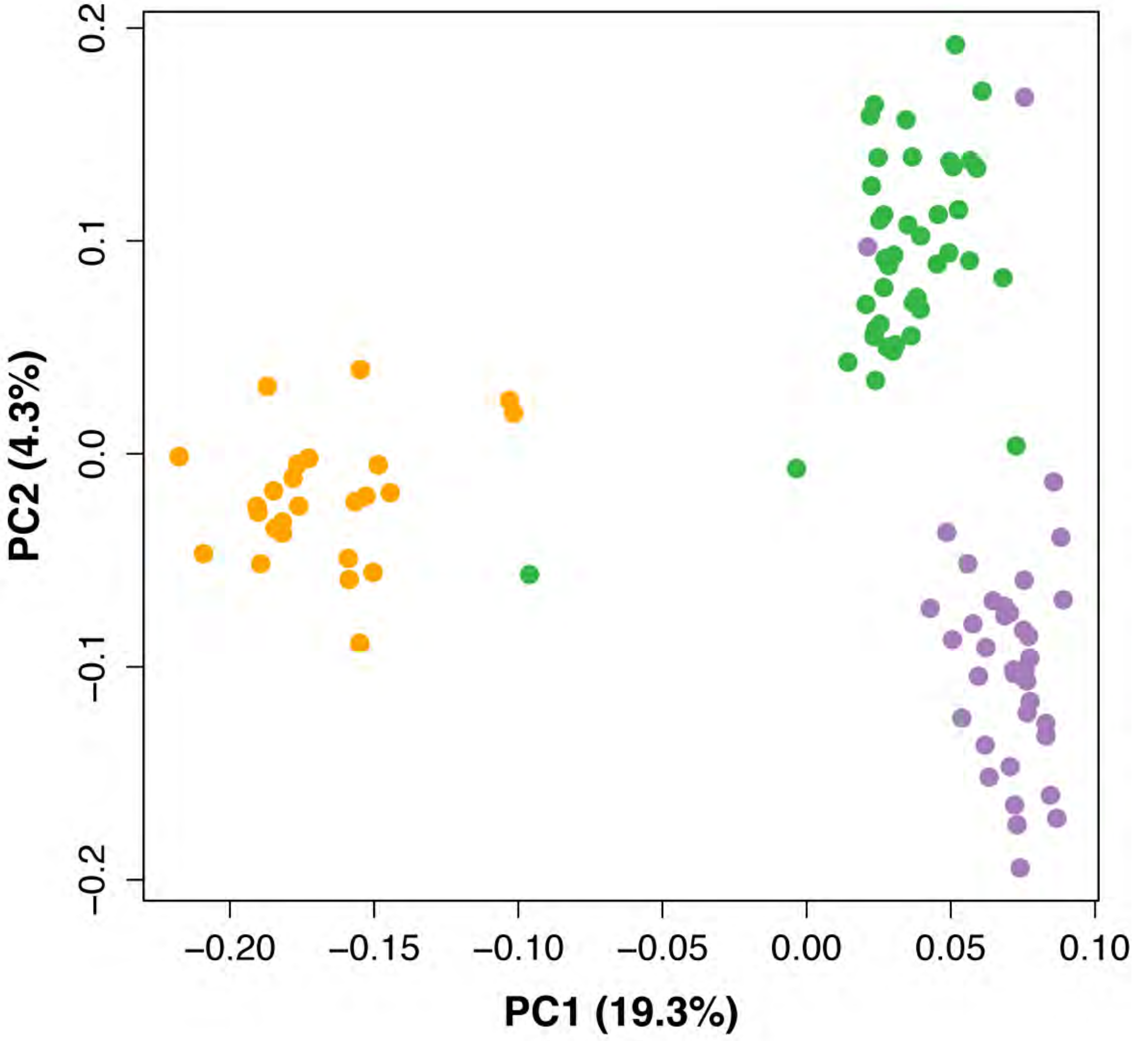


**S14 Fig. PCA of only neutral SNPs from the ‘1k’ dataset.**

**Table S10. The results of the Tracy-Widom test for all SNPs from the '10k' SNP dataset.**

| Principal Component | Eigen Values | Tracy-Widom Statistics | *p*-values | Percentage of Variation[a] | Percentage of Variation of Significant Eigenvalues |
|---|---|---|---|---|---|
| 1 | 39470 | 25.6 | 8.00e-09 | 19.4 | 54.8 |
| 2 | 9527 | 30.9 | 8.00e-09 | 4.7 | 13.2 |
| 3 | 6111 | 17.8 | 8.00e-09 | 3.0 | 8.5 |
| 4 | 5111 | 12.7 | 8.00e-09 | 2.5 | 7.1 |
| 5 | 4030 | 2.7 | 0.003086 | 2.0 | 5.6 |
| 6 | 4013 | 4.2 | 0.0001327 | 2.0 | 5.6 |
| 7 | 3828 | 3.6 | 0.0005331 | 1.9 | 5.3 |

[a]The first 7 principal components were significant, however, PCs 1 and 2 account for the majority of the statistically significant variation in the dataset.

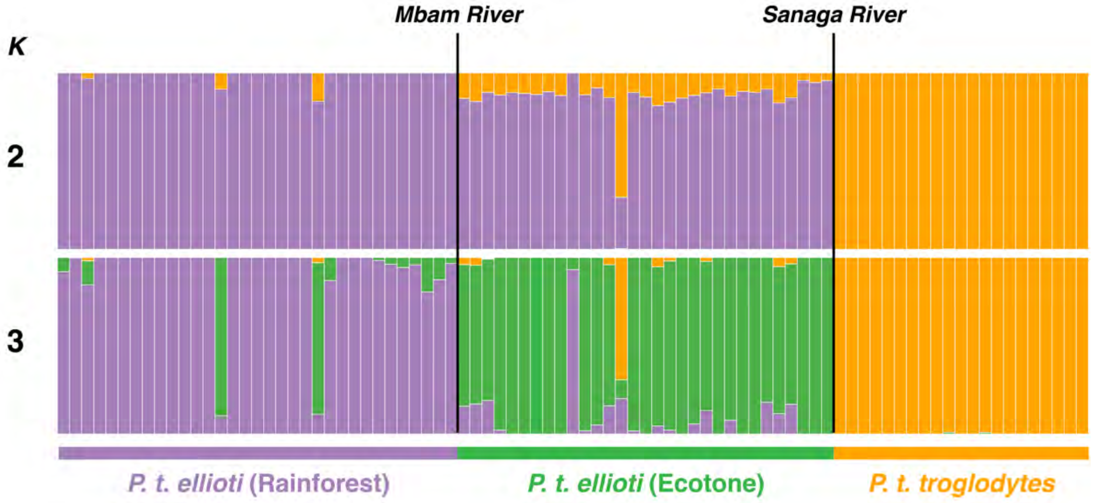


**S15 Fig. ADMIXTURE bar plots for *K*=2-3.**
These three populations correspond to known population structure. However, at *K*=2, there is a signal of possible historic gene flow of *P. t. troglodytes* into *P. t. elliot*i (Ecotone). Moreover, the is one individual (CMMD06 – also identified in the PCA) as being a potential *ellioti/troglodytes* hybrid. At *K*=3, we see evidence of three additional individuals that may be Rainforest/Ecotone hybrids, as well as evidence of mixing between the populations.

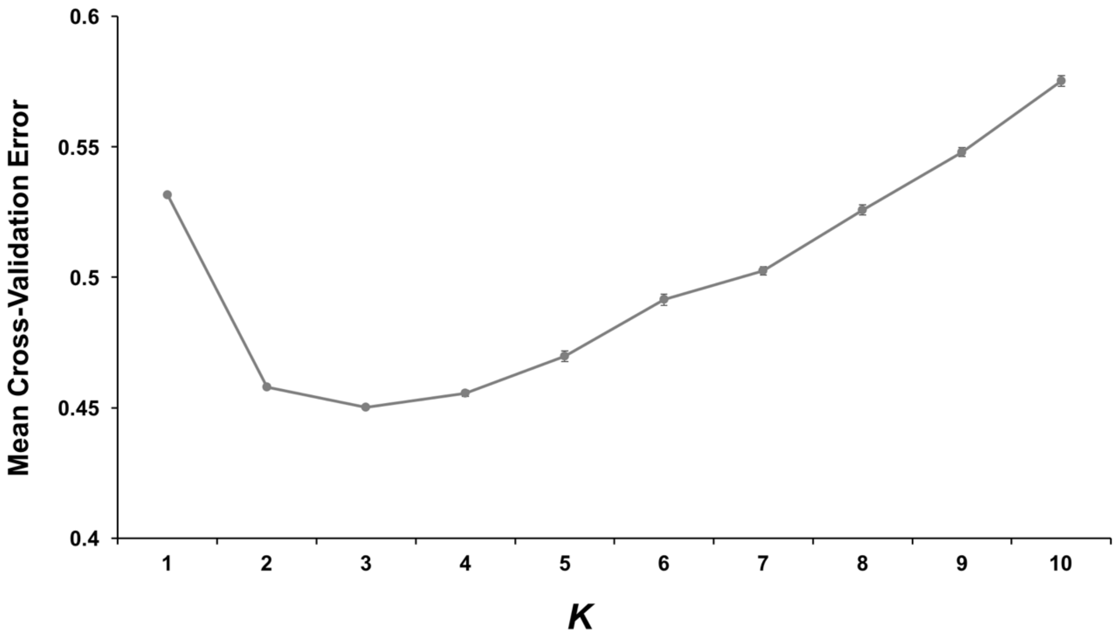


**S16 Fig. The cross-validation error results of ADMIXTURE analysis of wild chimpanzees (‘10k’ dataset).**

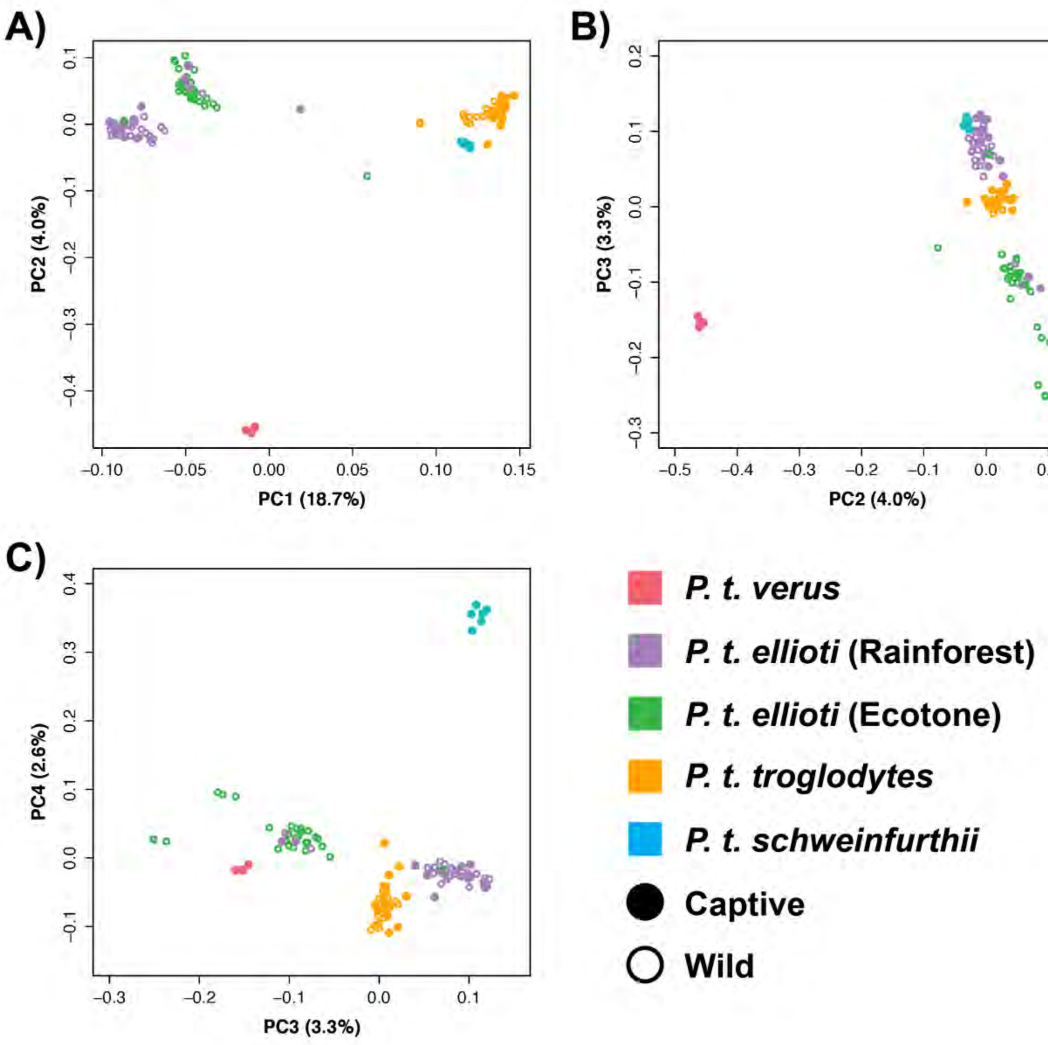


**S17 Fig. PCA results of the merged captive and wild datasets.**

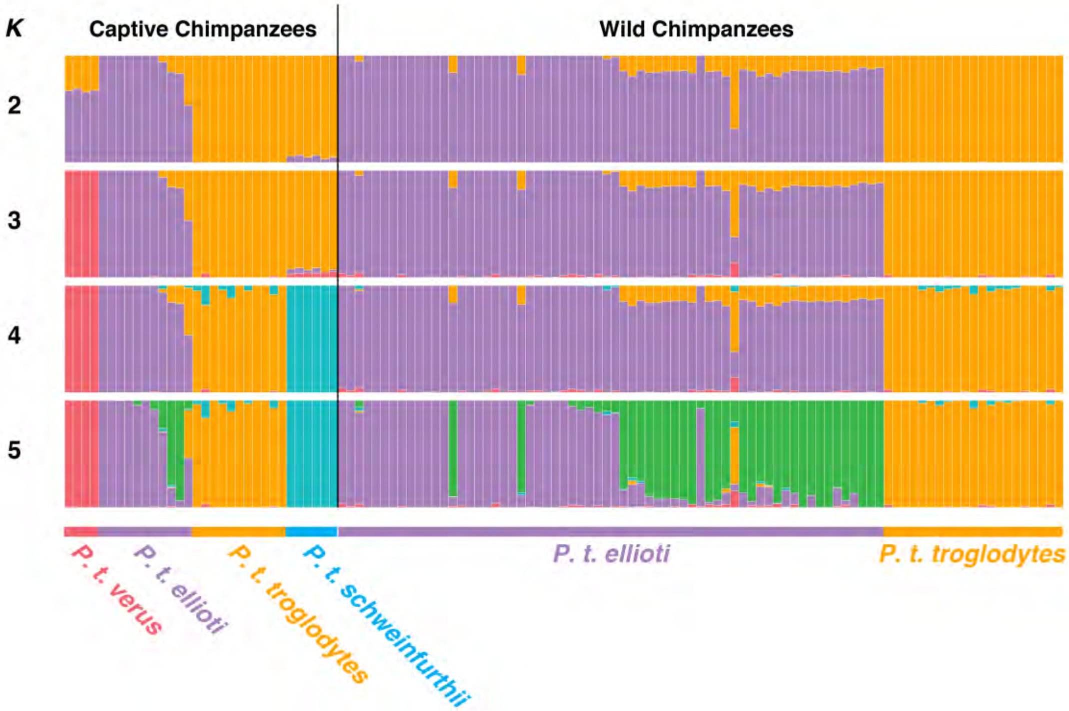


**S18 Fig. ADMIXTURE bar plots for *K*=2-5 for merged captive and wild datasets.**

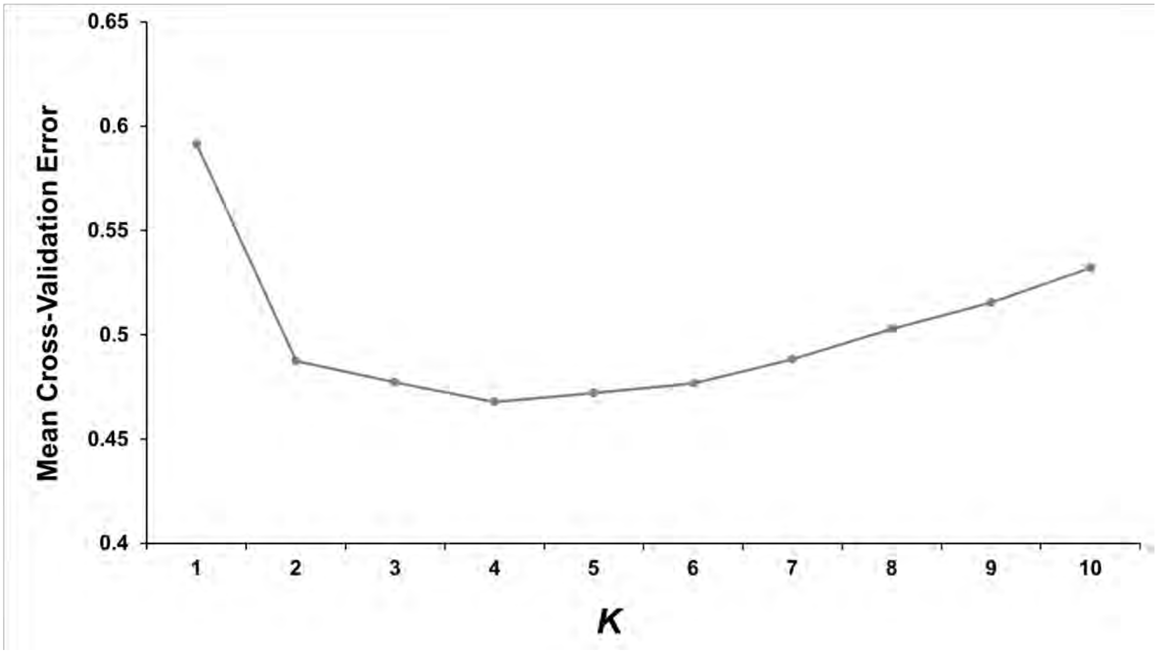


**S19 Fig. The cross-validation error results of ADMIXTURE analysis of merged captive and wild chimpanzees ('10k' dataset).**

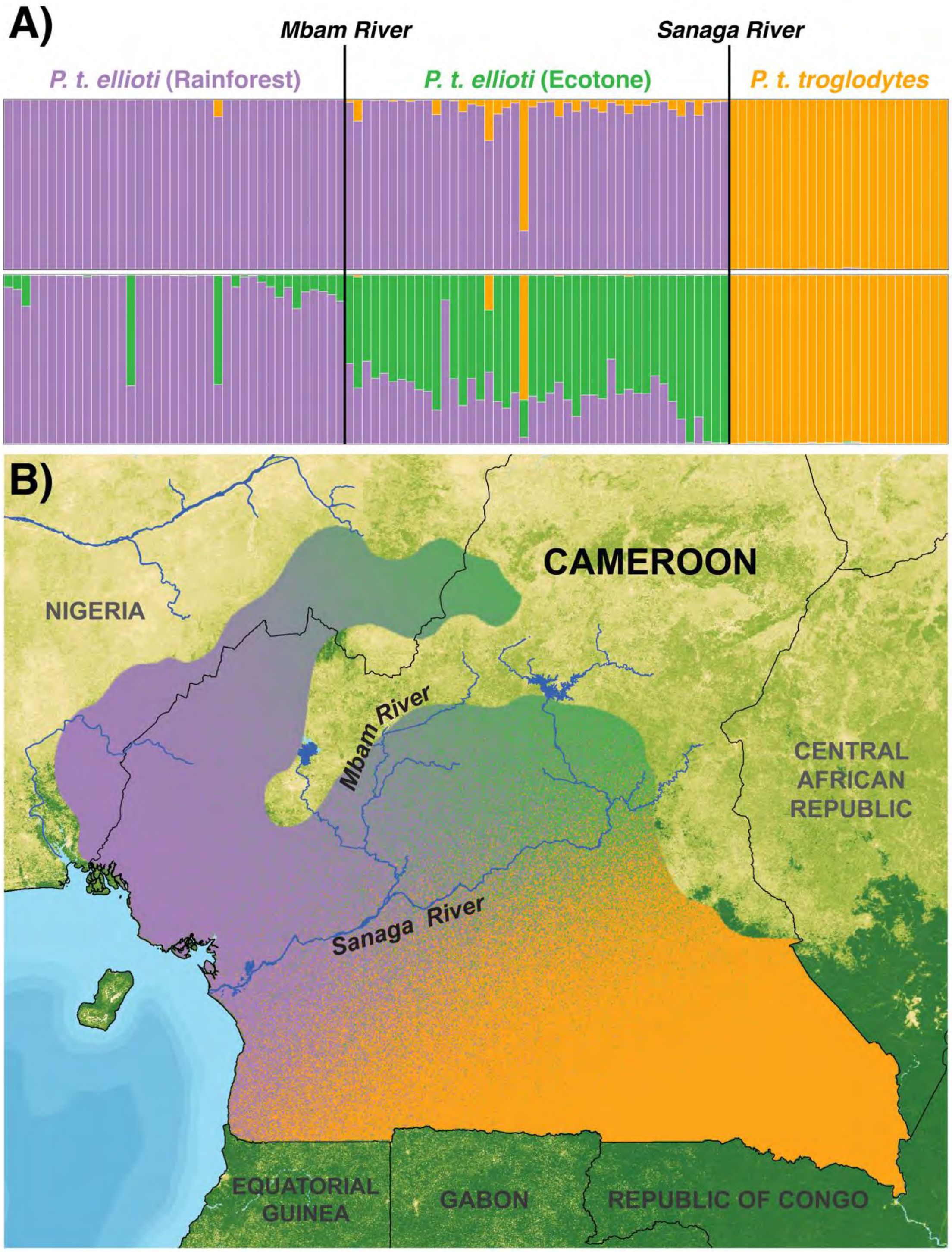


**S20 Fig. Cluster analysis and spatial interpolation of population structure.**
(A) TESS bar plots showing individual proportions of ancestry of wild chimpanzees.
(B) Spatial interpolation of the *Q* matrix for *K*=3 generated using TESS and Ad-Mixer.

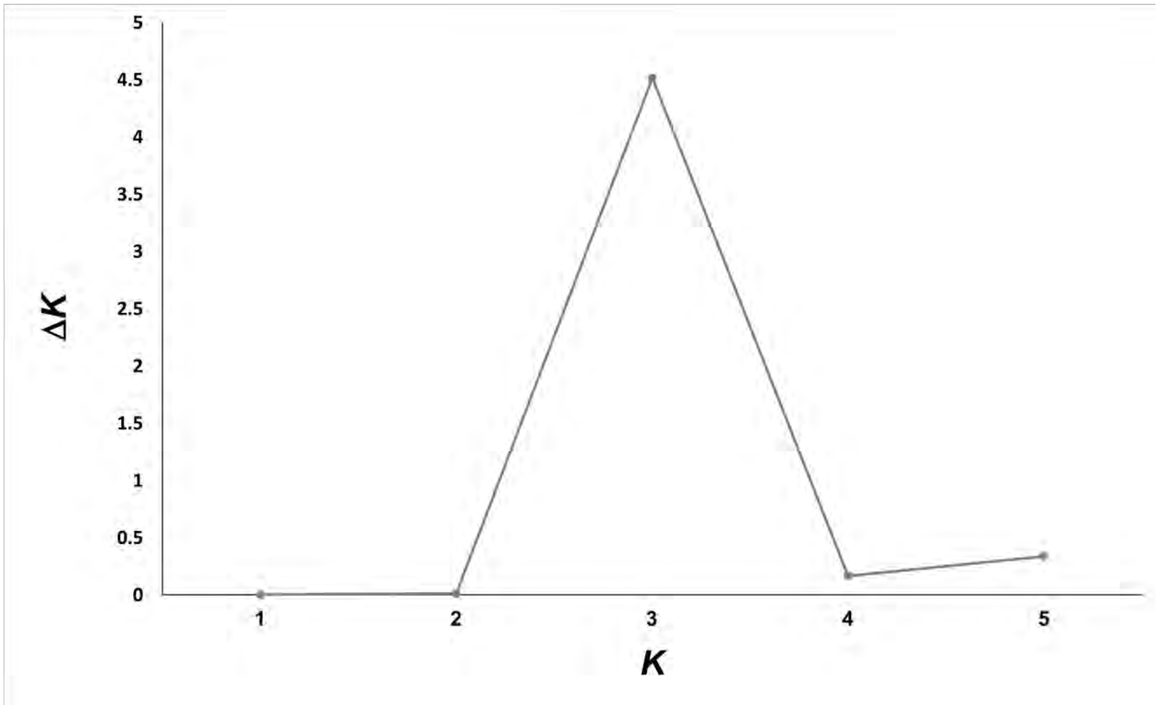


**S21 Fig. Estimating $K_{MAX}$ from TESS analysis.**
$\Delta K$ values estimated for $K$=1-5 across 10 replicate runs.

#### Table S11. Analysis of Molecular Variance (AMOVA).

| Partition | Fixation indices | Variance Components | Percentage of variation | *p*-value |
|---|---|---|---|---|
| **Among populations ($\Phi_{CT}$)** | 0.17 | 199.88 | 16.99 | $p < 0.001$ |
| **Among sample sites in populations ($\Phi_{SC}$)** | 0.06 | 60.53 | 5.15 | $p < 0.001$ |
| **Within sample sites ($\Phi_{IS}$)** | 0.22 | 915.84 | 77.86 | $p < 0.001$ |

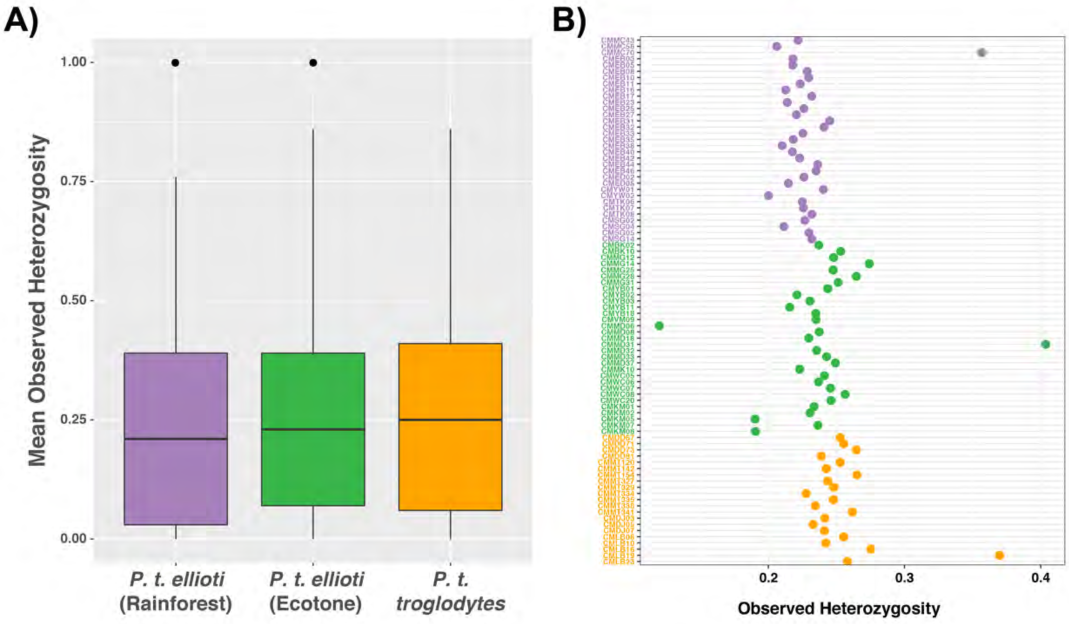


**S22 Fig. Mean observed heterozygosity.**

(A) Heterozygosity of all loci for all individuals grouped by population.

(B) Heterozygosity for all individuals.

There were no significant differences between heterozygosity for each population.

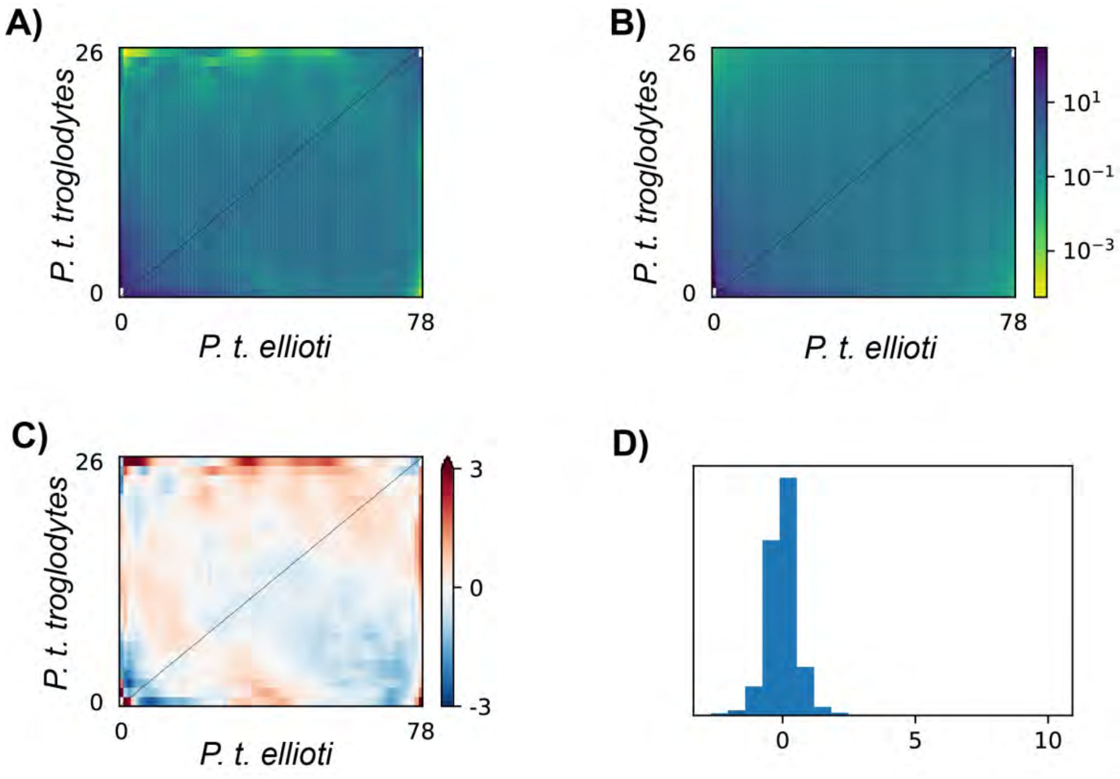


**S23 Fig. Posterior plots of model performance.**
(A) The observed Joint SFS for *P. t. troglodytes* and *P. t. ellioti*.
(B) the simulated Joint SFS for *P. t. troglodytes* and *P. t. ellioti* under the most likely asymmetric migration scenario obtained from *δaδi.*
(C) The residuals between the modeled and observed Joint SFS.
(D) A 1D histogram of the residual values between the model and the observed data.

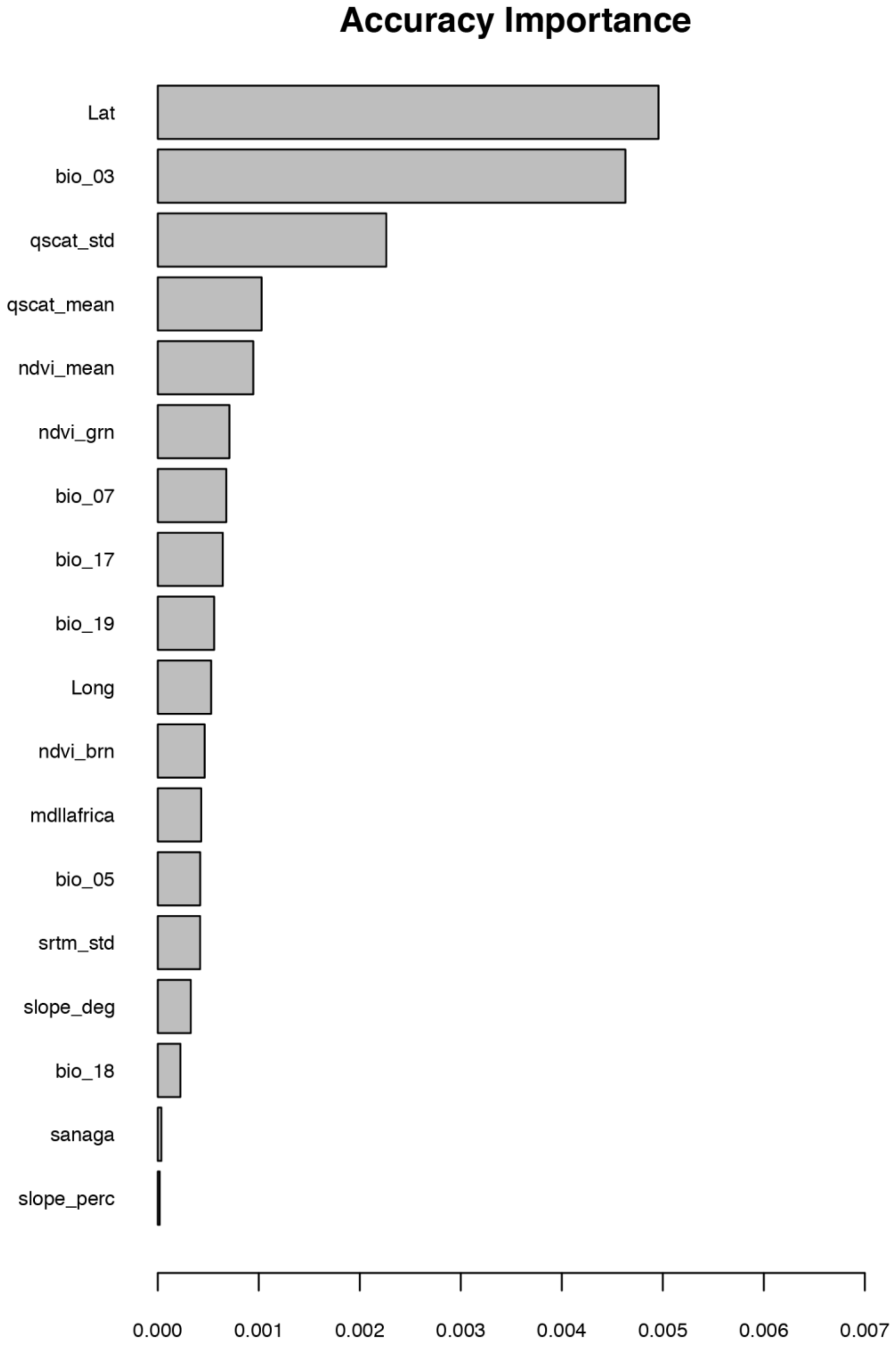


**S24 Fig. $R^2$ weighted importance of the environmental predictor variables to the Gradient Forest model of gene-environment relationships.**

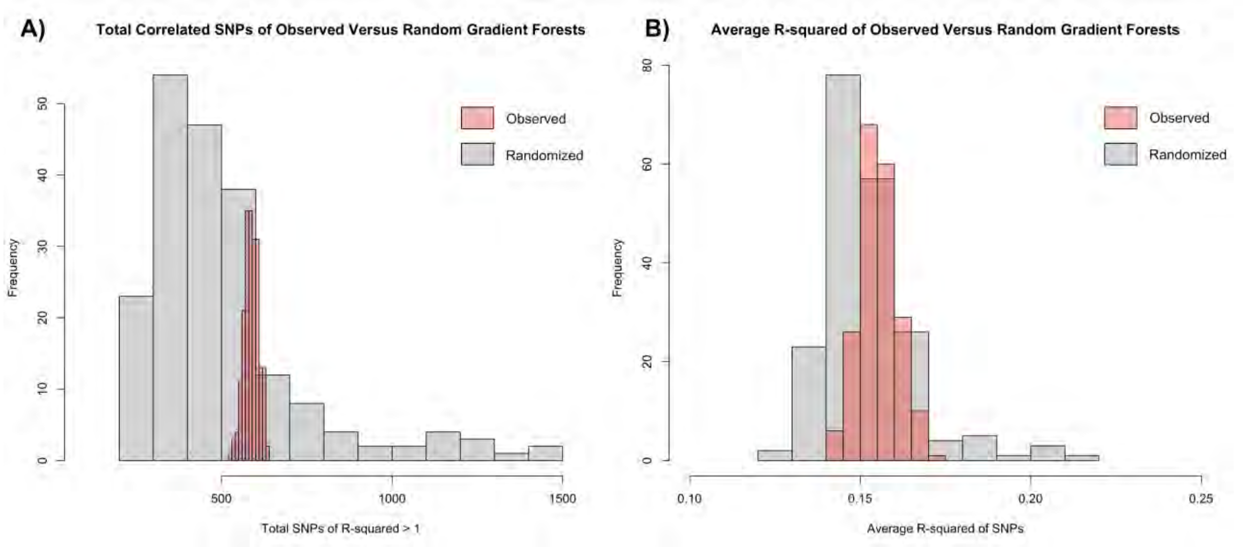


**S25 Fig. Results of randomized gradient forest models (n=200), as compared to results from the observed data (n=200).**

An average 588 of 7,878 SNPs demonstrated a positive $R^2$ with at least one environmental variable, with an average $R^2 = 0.155$ in the Observed data distribution (n=200, represented by the red histograms in A) and B) above). A significantly different average was obtained when randomizing the associations between the genomic data (SNPs) and environmental predictors for both total SNPs with a positive $R^2$ (average total = 504, t = 5.011(unequal variances df = 202.28), $p < 0.0001$), as well as for the average $R^2$ (average = 0.152, t = 2.806 (unequal variance df = 261.97), p = 0.0054) of the randomized gradient forests runs (n=200).

**Table S12. Enriched GO terms in the 'Biological Processes' domain for environmentally associated outliers (LFMM and gradient forest) in wild chimpanzees.**

| GO ID | Term Description | Genes[a] | p-value | Count |
|---|---|---|---|---|
| GO:0007411 | Axon guidance | SEMA3E, EPHA5, NRP1, LAMA1, CCDC141, EXT1, SEMA5A, CNTN6, CNTN4, PALLD, EPHA6, DCC, DSCAM, SEMA3F, DAG1, EPHA7, PTPRO, ROBO4, LGI1, CNTN5, PAX6, EFNB2, TUBB3 | 3.14E-06 | 23 of 129 |
| GO:0071560 | Cellular response to transforming growth factor beta stimulus | PDE3A, WNT2, TGFB1, EPB41L5, COL4A2, SOX5, WWOX | 0.001108359 | 7 of 22 |
| GO:0009611 | Response to wounding | EDNRA, GRIN2A, SLC1A2, TGFB1, TMPRSS4, PAX6, ODAM | 0.001814455 | 7 of 24 |
| GO:0070593 | Dendrite self-avoidance | DSCAM, CCDC141, EXT1, ROBO4, CNTN6, PALLD | 0.002017233 | 6 of 17 |
| GO:0035556 | Intracellular signal transduction | ARHGAP29, CILK1, KSR2, ADCY8, ASB17, DGKB, DGKI, DCDC2, MAP3K20, DCDC1, VAV3, BLNK, LYN, NRG3, ADCY1, RGS6, BRAF, DEPDC5, PLCL2, DEPDC1, PLCB1, MAST4, ARHGEF4, , DCLK3, PRKCA, TNIK, NRG2, CCDC68, PAK6, DAPK1 | 0.003408536 | 32 of 327 |
| GO:0051642 | Centrosome localization | NIN, NDEL1, NUBP1, SYNE2, NDE1 | 0.003514262 | 5 of 12 |
| GO:0022408 | Negative regulation of cell-cell adhesion | PODXL, TGFB1, TNR, JAG1, EPB41L5 | 0.003514262 | 5 of 12 |
| GO:0007156 | Homophilic cell adhesion via plasma membrane adhesion molecules | CCDC141, CDHR3, CNTN6, PALLD, SDK1, CLSTN2, CDH13, PTPRM, DSCAM, PTPRT, CDH20, NCAM2, ROBO1, PECAM1, ROBO4, KIRREL3 | 0.003738745 | 16 of 122 |
| GO:1903779 | Regulation of cardiac conduction | RYR3, TNNI3K, RYR2, ITPR1, ITPR2 | 0.004850741 | 5 of 13 |
| GO:0002027 | Regulation of heart rate | EDNRA, CASQ2, TNNI3K, RYR2, MYH6, BVES | 0.005493921 | 6 of 21 |
| GO:0048814 | Regulation of dendrite morphogenesis | HECW1, TNIK, HECW2, NEDD4L, SIPA1L1 | 0.006490105 | 5 of 14 |
| GO:0045109 | Intermediate filament organization | VIM, DSP, BFSP2, KRT2, KRT20 | 0.006490105 | 5 of 14 |
| GO:0030336 | Negative regulation of cell migration | PTPRU, PTPRT, DAG1, ROBO1, DACH1, DUSP3, SULF1, JAG1, PTPRK, OSBPL8, MITF | 0.00713265 | 11 of 72 |
| GO:0008217 | Regulation of blood pressure | TRHDE, EDNRA, UMOD, UTS2B, ACVRL1, EXT1, MYH6, NPY | 0.007598714 | 8 of 41 |

| | | | | |
|---|---|---|---|---|
| GO:0035249 | Sysnaptic transmission, glutamatergic | PRKN, GRIK1, GRID1, GRIK3, EXT1, GRID2 | 0.008271382 | 6 of 23 |
| GO:0060079 | Excitatory postsynaptic potential | GRIN2B, GRIN2A, DGKI, PPP3CA, GRID2 | 0.008458802 | 5 of 15 |
| GO:0045669 | Positive regulation of osteoblast differentiation | GNAS, WWTR1, SUCO, BMP6, JAG1, NELL1, SCUBE2, FBN2 | 0.009878242 | 8 of 43 |
| GO:0007162 | Negative regulation of cell adhesion | CDH13, DSCAM, ANGPT1, PODXL, PDE3B, RIPOR2 | 0.009970574 | 6 of 24 |
| GO:0030198 | Extracellular matrix organization | HPSE2, COL13A1, COL19A1, ADAMTS19, COL25A1, TNR, TNFRSF11B, ITGA2, ABI3BP, ITGA9, SERPINB5, ADAMTS5, ADAMTS17, NPHP3, ADAMTS18, SMOC1, COL4A1, COL4A2 | 0.010436808 | 18 of 162 |
| GO:0009410 | Response to xenobiotic stimulus | PTPRM, GNAS, UMOD, TGFBR2, GRIN2A, SLC1A2, EMX1, ATG5, VAV3 | 0.010554869 | 9 of 54 |
| GO:0051239 | Regulation of multicellular organismal process | TGFBR2, ACVRL1, AMHR2, ITPR1 | 0.011777424 | 4 of 9 |
| GO:0007154 | Cell communication | FREM1, SLC8A1, GJD2, GJD4, SLC8A3, FREM3 | 0.01189458 | 6 of 25 |
| GO:0043010 | Camera-type eye development | FBN1, GRHL2, CACNA1C, LAMA1, FBN2, MITF | 0.014055845 | 6 of 26 |
| GO:0030509 | BMP signaling pathway | USP15, SMPD3, BMP6, ACVRL1, TGFB1, EXT1, RGMA, BMP5, SMAD3 | 0.014456014 | 9 of 57 |
| GO:0003094 | Glomerular filtration | MYO1E, EDNRA, UMOD, SULF1 | 0.016124967 | 4 of 10 |
| GO:0007420 | Brain development | IMMP2L, GRHL2, NFIB, ADGRL3, EMX1, CCDC39, MACROD2, PLCB1, TGFBR2, EPHA7, CNTN5, PPT1, COL4A1 | 0.016705656 | 13 of 106 |
| GO:0010587 | miRNA catabolic process | SND1, PARN, LIN28B | 0.017831679 | 3 of 4 |
| GO:0001764 | Neuron migration | GPM6A, TOP2B, CEP85L, ASTN2, PAX6, KIRREL3, DCC, PRKG1, DCDC2, NDE1 | 0.018393559 | 10 of 71 |
| GO:0007492 | Endoderm development | TGFB1, EXT1, EPB41L5, SMAD3 | 0.021252347 | 4 of 11 |
| GO:0001974 | Blood vessel remodeling | EDNRA, ACVRL1, EXT1, ATG5, JAG1 | 0.023984008 | 5 of 20 |
| GO:0002503 | Peptide antigen assembly with MHC class II protein complex | PATR-DOB, HLA-DPB1, HLA-DPA1, HLA-DQB1 | 0.027165042 | 4 of 12 |
| GO:0001958 | Endochondral ossification | SMPD3, GNAS, COL13A1, BMP6, EXT1 | 0.028334617 | 5 of 21 |

| | | | | |
|---|---|---|---|---|
| GO:0007160 | Cell-matrix adhesion | TIAM1, FREM1, ITGBL1, COL13A1, CD96, HPSE, EPDR1, TBCEL | 0.031858728 | 8 of 54 |
| GO:0030335 | Positive regulation of cell migration | SEMA3E, EGFR, ATP8A1, TGFB1, SEMA5A, PDGFC, PPP3CA, SYNE2, CDH13, TIAM1, SEMA3F, PODXL, PECAM1, LAMC2, LYN | 0.032231929 | 15 of 143 |
| GO:0047496 | Vesicle transport along microtubule | NDEL1, TRAK1, BICDL1, NDE1 | 0.033859333 | 4 of 13 |
| GO:0060389 | Pathway-restricted SMAD protein phosphorylation | USP15, TGFBR2, TGFB1, AMHR2 | 0.033859333 | 4 of 13 |
| GO:0060218 | Hematopoietic stem cell differentiation | CHD2, EXT1, UFL1, MLLT3 | 0.033859333 | 4 of 13 |
| GO:0019933 | cAMP-mediated signaling | PDE3A, CAP1, PDE7B, ADGRG6 | 0.033859333 | 4 of 13 |
| GO:0050804 | Modulation of synaptic transmission | GRIK1, GRID1, GRIK3, NRG3, DGKB, AKAP12, GRID2 | 0.036558271 | 7 of 44 |
| GO:0045332 | Phospholipid translocation | ATP10D, ATP10B, ABCB4, ATP8A1, ABCB1 | 0.038355754 | 5 of 23 |
| GO:0008104 | Protein localization | NIN, PARD3, NBEAL2, CCDC68, NBEA, PARD3B, CEP128 | 0.040241136 | 7 of 45 |
| GO:0010467 | Gene expression | EDNRA, GRM3, TGFB1, EXT1, ROS1, PAX6, MLLT3 | 0.040241136 | 7 of 45 |
| GO:0051056 | Regulation of small GTPase mediated signal transduction | RALGAPA2, ARHGAP29, ARHGAP28, SIPA1L1, SIPA1L2, VAV3 | 0.041070945 | 6 of 34 |
| GO:0043652 | Engulfment of apoptotic cell | RHOBTB1, XKR4, MEGF10, XKR5 | 0.041323668 | 4 of 14 |
| GO:0061028 | Establishment of endothelial barrier | EDNRA, HPSE, PECAM1, ROBO4 | 0.049539884 | 4 of 15 |
| GO:0060441 | Epithelial tube branching involved in lung morphogenesis | DAG1, LAMA1, RSPO2, EXT1 | 0.049539884 | 4 of 15 |
| GO:0016601 | Rac protein signal transduction | CDH13, TIAM1, ELMO1, EPS8 | 0.049539884 | 4 of 15 |

[a]Enrichment analysis was performed using a background population of genes found outside of regions under natural selection.

**Table S13. Enriched KEGG pathways for environmentally associated outliers (LFMM and gradient forest) in wild chimpanzees.**

| KEGG ID | Term Description | Genes[a] | *p*-value | Count |
|---|---|---|---|---|
| ptr05414 | Dilated cardiomyopathy | PRKACB, CACNA2D3, ADCY8, CACNA1C, ADCY1, RYR2, LAMA1, TGFB1, ITGA2, ITGA9, GNAS, DAG1, CACNG6, SLC8A1, SLC8A3, TPM1, MYH6, IGF1 | 0.00001 | 18 of 84 |
| ptr04724 | Glutamatergic synapse | PRKACB, GRIN2B, GRIK3, GRM3, PLA2G4A, ADCY8, CACNA1C, ADCY1, SLC1A2, PPP3CA, ITPR1, PLCB1, GNAS, PRKCA, GRIK1, GRIN2A, GNG3, ITPR2 | 0.00008 | 18 of 102 |
| ptr04020 | Calcium signaling pathway | GRIN2B, PTGER3, PRKACB, ADCY8, EGFR, CASQ2, PDGFC, CACNA1E, RYR3, SLC8A3, CAMK1D, EDNRA, ADRA1B, CACNA1C, ADCY1, RYR2, NOS3, PPP3CA, ITPR1, PLCB1, GNAS, PRKCA, SLC8A1, GRIN2A, FGF3, GNA14, ITPR2 | 0.00059 | 27 of 223 |
| ptr05410 | Hypertrophic cardiomyopathy | CACNA2D3, CACNA1C, RYR2, LAMA1, TGFB1, ITGA2, ITGA9, DAG1, CACNG6, SLC8A1, SLC8A3, TPM1, MYH6, IGF1 | 0.00080 | 14 of 81 |
| ptr04720 | Long-term potentiation | PRKACB, GRIN2B, PRKCA, ADCY8, GRIN2A, CACNA1C, ADCY1, BRAF, PPP3CA, ITPR1, PLCB1, ITPR2 | 0.00085 | 12 of 62 |
| ptr04921 | Oxytocin signaling pathway | PRKACB, CACNA2D3, EGFR, PLA2G4A, ADCY8, CACNA1C, ADCY1, RYR2, NOS3, ROCK1, PPP3CA, ITPR1, PLCB1, GNAS, CACNG6, PRKCA, RYR3, CAMK1D, ITPR2 | 0.00096 | 19 of 136 |
| ptr04713 | Circadian entrainment | PRKACB, GRIN2B, ADCY8, CACNA1C, ADCY1, RYR2, ITPR1, PLCB1, PRKG1, GNAS, PRKCA, RYR3, GRIN2A, GNG3 | 0.00101 | 14 of 83 |
| ptr04015 | Rap1 signaling pathway | GRIN2B, MAGI2, EGFR, ADCY8, PARD3, ADCY1, ANGPT1, BRAF, RAPGEF5, PDGFC, DOCK4, PLCB1, PIK3CB, TIAM1, GNAS, PRKCA, GRIN2A, RAPGEF4, SIPA1L1, SIPA1L2, VAV3, FGF3, IGF1 | 0.00118 | 23 of 185 |

| | | | | |
|---|---|---|---|---|
| ptr02010 | ABC transporters | ABCA13, ABCB8, ABCG5, TAP2, ABCG8, ABCB4, ABCC11, ABCA6, ABCB1 | 0.00137 | 9 of 38 |
| ptr04730 | Long-term depression | GNAS, PRKCA, PLA2G4A, BRAF, ITPR1, PLCB1, ITPR2, GRID2, IGF1, LYN, PRKG1 | 0.00138 | 11 of 56 |
| ptr04970 | Salivary secretion | PRKACB, ADRA1B, ADCY8, ADCY1, ITPR1, SLC12A2, PLCB1, PRKG1, GNAS, PRKCA, RYR3, LYZ, ITPR2 | 0.00146 | 13 of 76 |
| ptr04540 | Gap junction | PRKACB, EGFR, ADCY8, ADCY1, GJD2, PDGFC, ITPR1, PLCB1, PRKG1, GNAS, PRKCA, ITPR2, TUBB3 | 0.00164 | 13 of 77 |
| ptr04261 | Adrenergic signaling in cardiomyocytes | KCNK2, PRKACB, CACNA2D3, ADRA1B, ADCY8, CACNA1C, ADCY1, RYR2, BVES, PLCB1, GNAS, CACNG6, PRKCA, SLC8A1, RAPGEF4, SLC8A3, TPM1, MYH6 | 0.00193 | 18 of 133 |
| ptr04070 | Phosphatidylinositol signaling system | IP6K3, MTMR6, DGKB, DGKI, INPP4B, ITPR1, PLCB1, PIK3CB, PRKCA, IP6K2, PIP4K2A, PIK3C2G, ITPR2, PI4KB | 0.00264 | 14 of 92 |
| ptr05146 | Amoebiasis | PRKACB, ADCY1, LAMA1, TGFB1, PLCB1, PIK3CB, GNAS, PRKCA, ARG2, LAMC2, CD1A, COL4A1, GNA14, COL4A2 | 0.00352 | 14 of 95 |
| ptr04510 | Focal adhesion | MYL10, EGFR, LAMA1, BRAF, ROCK1, PDGFC, TNR, COL6A3, ITGA2, , PIK3CB, ITGA9, CHAD, PRKCA, PAK6, LAMC2, VAV3, COL4A1, COL4A2, IGF1, PARVA | 0.00433 | 21 of 180 |
| ptr05412 | Arrhythmogenic right ventricular cardiomyopathy | ITGA9, DAG1, CACNG6, CACNA2D3, SLC8A1, CACNA1C, DSP, RYR2, LAMA1, SLC8A3, ITGA2 | 0.00539 | 11 of 67 |
| ptr04024 | cAMP signaling pathway | KCNK2, PTGER3, PRKACB, GRIN2B, EDNRA, PDE3A, ADCY8, CACNA1C, ADCY1, RYR2, BRAF, ROCK1, CNGA3, BVES, PIK3CB, NPY, TIAM1, GNAS, GRIN2A, RAPGEF4, VAV3, PDE3B | 0.00602 | 22 of 198 |
| ptr04512 | ECM-receptor interaction | ITGA9, FREM1, CHAD, DAG1, LAMA1, LAMC2, TNR, ITGA2, COL6A3, COL4A1, COL4A2, | 0.00678 | 12 of 80 |
| ptr04927 | Cortisol synthesis and secretion | PRKACB, KCNK2, GNAS, ADCY8, CACNA1C, ADCY1, ITPR1, PLCB1, ITPR2, | 0.00704 | 10 of 59 |

| | | | | |
|---|---|---|---|---|
| ptr04360 | Axon guidance | SEMA3E, EPHA5, NRP1, PARD3, ROCK1, SEMA5A, PPP3CA, EPHA6, PIK3CB, DCC, SEMA3F, PRKCA, ROBO1, PAK6, EPHA7, ABLIM1, RGMA, TRPC4, EFNB2 | 0.00706 | 19 of 163 |
| ptr04976 | Bile secretion | PRKACB, ABCG5, GNAS, AQP9, ABCG8, ADCY8, SLC10A1, ADCY1, ABCB4, SLCO1B1, ABCB1 | 0.00738 | 11 of 70 |
| ptr04371 | Apelin signaling pathway | PRKACB, ADCY8, ADCY1, RYR2, NOS3, JAG1, ITPR1, PLCB1, RYR3, SLC8A1, GNG3, SLC8A3, PDE3B, ITPR2, SMAD3 | 0.00843 | 15 of 117 |
| ptr05416 | Viral myocarditis | DAG1, PATR-DOB, HLA-DPB1, LAMA1, CD80, HLA-DPA1, HLA-DQB1, CASP9, MYH6 | 0.00916 | 9 of 51 |
| ptr04270 | Vascular smooth muscle contraction | PRKACB, EDNRA, ADRA1B, PLA2G4A, ADCY8, CACNA1C, ADCY1, BRAF, ROCK1, ITPR1, PLCB1, PRKG1, GNAS, PRKCA, ITPR2 | 0.01122 | 15 of 121 |
| ptr04926 | Relaxin signaling pathway | PRKACB, EGFR, ADCY8, ADCY1, NOS3, TGFB1, PLCB1, PIK3CB, GNAS, PRKCA, TGFBR2, GNG3, COL4A1, COL4A2 | 0.01207 | 14 of 110 |
| ptr04611 | Platelet activation | PRKACB, PLA2G4A, ADCY8, ADCY1, NOS3, ROCK1, ITGA2, ITPR1, PLCB1, PIK3CB, PRKG1, GNAS, ITPR2, LYN | 0.01297 | 14 of 111 |
| ptr04924 | Renin secretion | PRKACB, EDNRA, GNAS, PDE3A, CACNA1C, PPP3CA, ITPR1, PDE3B, PLCB1, ITPR2 | 0.01314 | 10 of 65 |
| ptr04750 | Inflammatory mediator regulation of TRP channels | PRKACB, GNAS, PRKCA, PLA2G4A, ADCY8, ADCY1, IL1RAP, ITPR1, PLCB1, ITPR2, PIK3CB, IGF1 | 0.01474 | 12 of 89 |
| ptr04022 | cGMP-PKG signaling pathway | EDNRA, ADRA1B, PDE3A, ADCY8, CACNA1C, ADCY1, NOS3, ROCK1, PPP3CA, ITPR1, PLCB1, PRKG1, SLC8A1, SLC8A3, PDE3B, MYH6, ITPR2 | 0.01559 | 17 of 151 |
| ptr05322 | Systemic lupus erythematosus | GRIN2B, PATR-DOB, HLA-DPB1, C7, GRIN2A, CD80, HLA-DPA1, HLA-DQB1, C2 | 0.01584 | 9 of 56 |
| ptr04923 | Regulation of lipolysis in adipocytes | PRKACB, PTGER3, GNAS, ADCY8, ADCY1, PDE3B, PIK3CB, NPY, PRKG1 | 0.01584 | 9 of 56 |

| | | | | |
|---|---|---|---|---|
| ptr04911 | Insulin secretion | PRKACB, GNAS, PRKCA, ADCY8, CACNA1C, RAPGEF4, ADCY1, RYR2, SNAP25, PLCB1, RIMS2 | 0.01821 | 11 of 80 |
| ptr04961 | Endocrine and other factor-regulated calcium reabsorption | PRKACB, GNAS, PRKCA, SLC8A1, ESR1, DNM3, SLC8A3, PLCB1 | 0.01891 | 8 of 47 |
| ptr04918 | Thyroid hormone synthesis | PRKACB, GNAS, PRKCA, ADCY8, DUOX1, ADCY1, GPX6, ITPR1, PLCB1, ITPR2 | 0.01897 | 10 of 69 |
| ptr04912 | GnRH signaling pathway | PRKACB, GNAS, PRKCA, EGFR, PLA2G4A, ADCY8, CACNA1C, ADCY1, ITPR1, PLCB1, ITPR2 | 0.01973 | 11 of 81 |
| ptr04659 | Th17 cell differentiation | CD247, IL17F, PATR-DOB, HLA-DPB1, TGFBR2, RORC, TGFB1, IL1RAP, PPP3CA, HLA-DPA1, HLA-DQB1, SMAD3 | 0.02147 | 12 of 94 |
| ptr05017 | Spinocerebellar ataxia | GRIN2B, MAP3K5, PSMB7, ATG13, OPA1, ATXN10, ITPR1, PLCB1, PIK3CB, PRKCA, GRIN2A, FGF14, PSMD3, PSMA1, ITPR2 | 0.02265 | 15 of 132 |
| ptr04915 | Estrogen signaling pathway | PRKACB, EGFR, ADCY8, ADCY1, NOS3, ITPR1, PLCB1, PIK3CB, GNAS, KRT24, ESR1, KRT20, ITPR2, KRT12 | 0.02655 | 14 of 122 |
| ptr05205 | Proteoglycans in cancer | PRKACB, HPSE2, ANK3, EGFR, BRAF, ROCK1, TGFB1, ITGA2, ITPR1, PIK3CB, TIAM1, PRKCA, ESR1, WNT2, HPSE, VAV3, ITPR2, IGF1 | 0.02663 | 18 of 174 |
| ptr01522 | Endocrine resistance | PRKACB, NOTCH4, GNAS, EGFR, ADCY8, ESR1, ADCY1, BRAF, JAG1, PIK3CB, IGF1 | 0.02673 | 11 of 85 |
| ptr04925 | Aldosterone synthesis and secretion | PRKACB, GNAS, PRKCA, ADCY8, CACNA1C, ADCY1, CAMK1D, ITPR1, PLCB1, ITPR2, | 0.02873 | 11 of 86 |
| ptr04971 | Gastric acid secretion | PRKACB, KCNK2, GNAS, PRKCA, ADCY8, ADCY1, ITPR1, PLCB1, ITPR2 | 0.03569 | 9 of 65 |
| ptr04935 | Growth hormone synthesis, secretion and action | PRKACB, GNAS, PRKCA, ADCY8, CACNA1C, GHR, ADCY1, ITPR1, PLCB1, ITPR2, PIK3CB, IGF1 | 0.04128 | 12 of 104 |
| ptr04972 | Pancreatic secretion | RAB27B, GNAS, PRKCA, ADCY8, ADCY1, RYR2, ITPR1, SLC12A2, PLCB1, ITPR2 | 0.04135 | 10 of 79 |

| | | | | |
|---|---|---|---|---|
| ptr04072 | Phospholipase D signaling pathway | GRM3, EGFR, PLA2G4A, ADCY8, ADCY1, DGKB, DGKI, PDGFC, PLCB1, PIK3CB, GNAS, PRKCA, RAPGEF4, DNM3 | 0.04380 | 14 of 131 |
| ptr05321 | Inflammatory bowel disease | IL17F, PATR-DOB, HLA-DPB1, RORC, TGFB1, HLA-DPA1, HLA-DQB1, SMAD3 | 0.04455 | 8 of 56 |
| ptr04310 | Wnt signaling pathway | PRKACB, BTRC, TLE4, VANGL1, DKK1, RSPO2, PPP3CA, PLCB1, SOST, PRKCA, WNT2, SIAH1, SFRP4, LGR4, SMAD3 | 0.04768 | 15 of 146 |

[a]Enrichment analysis was performed using a background population of genes found outside of regions under natural selection.

**Table S14. Enriched GO terms in the 'Biological Processes' domain for environmentally associated outliers (LFMM and gradient forest) in wild chimpanzees.**

| GO/KEGG ID | Term Description | Genes[a] | p-value | Count |
|---|---|---|---|---|
| GO:0007411 | Axon guidance | SEMA3E, EPHA5, NRP1, LAMA1, CCDC141, EXT1, CNTN6, CNTN4, PALLD, EPHA6, DCC, SEMA3F, DAG1, EPHA7, CNTN5, PAX6, EFNB2, TUBB3 | 0.008 | 18 of 26 |
| ptr04360 | Axon guidance | SEMA3E, EPHA5, NRP1, PARD3, ROCK1, EPHA6, PIK3CB, DCC, SEMA3F, PRKCA, ROBO1, EPHA7, ABLIM1, TRPC4, EFNB2 | 0.040 | 15 of 24 |

[a]Enrichment analysis was performed using a background population of genes found outside of regions under natural selection.

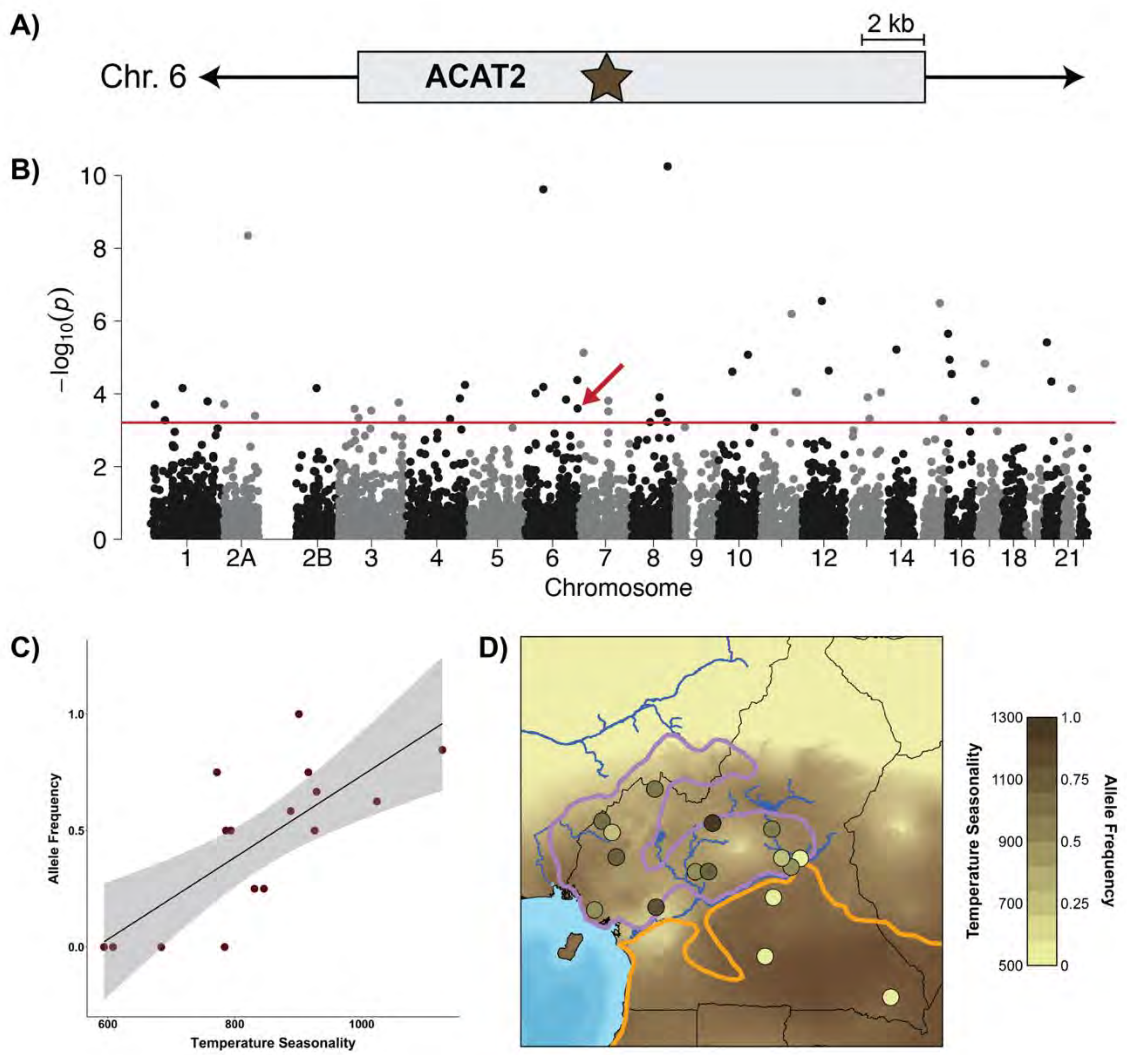


**S26 Fig. Evidence of selective pressures on acetyl-CoA acetyltransferase 2 (*ACAT2*).**

(A) Map of the *ACAT2* gene on chromosome 6 with brown star representing the SNP identified through outlier analysis between *P. t. troglodytes* and *P. t. elliotì*.

(B) Manhattan plot showing the significance (as the negative $\log_{10}$ p-value) of SNP associations with the environmental variable temperature seasonality. Grey colors distinguish different chromosomes. The red line represents the threshold for significant association ($p = 0.05$). The SNP contained in the ACAT2 gene is highlighted by the red arrow.

(C) Correlation between the allele frequency of the SNP contained in the ACAT2 gene and temperature seasonality values at each corresponding sampling location ($R^2 = 0.5615$, $p = 0.0005$).

(D) Allele frequencies of the SNP contained in the ACAT2 gene across Cameroon. Sampling sites are represented by circles that are shaded according to the frequency of the allele within the population. SNP frequencies are plotted against temperature seasonality across the region.

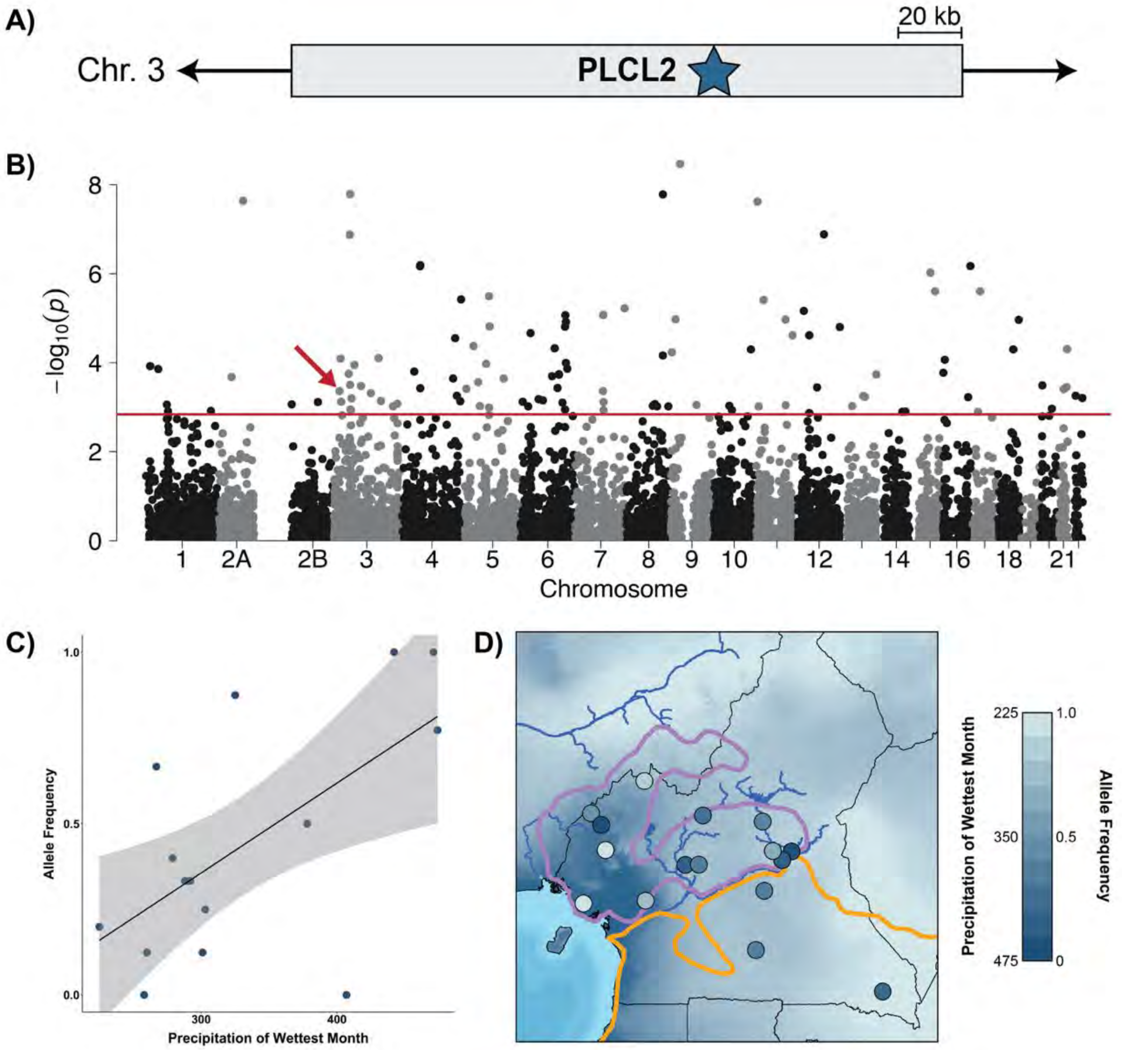


**S27 Fig. Evidence of selective pressures on phospholipase C like 2 (*PLCL2*).**
(A) Map of the *PLCL2* gene on chromosome 3 with brown star representing the SNP identified through outlier analysis between *P. t. troglodytes* and *P. t. elliotі*.
(B) Manhattan plot showing the significance (as the negative $\log_{10}$ p-value) of SNP associations with the environmental variable precipitation of wettest month. Grey colors distinguish different chromosomes. The red line represents the threshold for significant association (p = 0.05). The SNP contained in the PLCL2 gene is highlighted by the red arrow.
(C) Correlation between the allele frequency of the SNP contained in the PLCL2 gene and precipitation of wettest month values at each corresponding sampling location ($R^2$ = 0.3422, p = 0.0102).
(D). Allele frequencies of the SNP contained in the PLCL2 gene across Cameroon. Sampling sites are represented by circles that are shaded according to the frequency of the allele within the population. SNP frequencies are plotted against precipitation of wettest month across the region.
